# Resurrecting the Alternative Splicing Landscape of Archaic Hominins using Machine Learning

**DOI:** 10.1101/2022.08.02.502533

**Authors:** Colin M. Brand, Laura L. Colbran, John A. Capra

## Abstract

Alternative splicing contributes to adaptation and divergence in many species. However, it has not been possible to directly compare splicing between modern and archaic hominins. Here, we unmask the recent evolution of this previously unobservable regulatory mechanism by applying SpliceAI, a machine-learning algorithm that identifies splice altering variants (SAVs), to high-coverage genomes from three Neanderthals and a Denisovan. We discover 5,950 putative archaic SAVs, of which 2,186 are archaic-specific and 3,607 also occur in modern humans via introgression (244) or shared ancestry (3,520). Archaic-specific SAVs are enriched in genes that contribute to many traits potentially relevant to hominin phenotypic divergence, such as the epidermis, respiration, and spinal rigidity. Compared to shared SAVs, archaic-specific SAVs occur in sites under weaker selection and are more common in genes with tissue-specific expression. Further underscoring the importance of negative selection on SAVs, Neanderthal lineages with low effective population sizes are enriched for SAVs compared to Denisovan and shared SAVs. Finally, we find that nearly all introgressed SAVs in humans were shared across Neanderthals, suggesting that older SAVs were more tolerated in modern human genomes. Our results reveal the splicing landscape of archaic hominins and identify potential contributions of splicing to phenotypic differences among hominins.

## 1 Introduction

While the paleontological and archaeological records provide evidence about some phenotypes of extinct hominins, most ancient tissues have not survived to the present. The discovery and successful sequencing of DNA genome-wide from a Denisovan [1] and multiple Neanderthal genomes [2–4] enabled direct comparisons of the genotypes of these archaic hominins to one another and anatomically modern humans. These data also enable the potential for indirect phenotypic comparisons by predicting archaic phenotypes from their genomes [5]. Diverse molecular mechanisms collectively shape the similarities and differences between archaic hominins and modern humans. Given that the biology linking genotype to organism-level phenotype is complex and that the mapping may not generalize across human populations [6], predicting “low-level” molecular phenotypes from genetic information is a promising alternative. Recent work has successfully explored such differences in protein-coding sequence [7] and differences relevant to gene expression, such as divergent gene regulation [8], differential methylation [9], and divergent 3D genome contacts [10].

Variation in gene splicing could also underlie phenotypic differences between archaic hominins and modern humans, but archaic splicing patterns have not been comprehensively quantified. Alternative splicing enables the production of multiple protein isoforms from a single gene [11–13]. The resulting proteomic diversity is essential for many processes, including development and establishing tissue identity [14]. Defects in splicing underlie many human diseases (e.g., [15–22]), and variation in splicing contributes to differences in traits in nonhuman species (see [23], Table 1). Further, alternative splicing can evolve rapidly and respond to environmental factors—suggesting it often contributes to adaptation [23–25] and species differences [26–28].

Splicing patterns are directly influenced by the nucleotide sequences surrounding splice sites [29]. This has enabled the development of many algorithms to predict alternative splicing from RNA-seq [30–32] or DNA sequence [33–37]. Beyond human clinical applications, methods that require only DNA sequence can be leveraged to understand alternative splicing in extant species for which acquiring RNA-seq data may be difficult to impossible or in extinct taxa, such as archaic hominins.

Here, we resurrect the genome-wide alternative splicing landscape of archaic hominins using SpliceAI, an algorithm that predicts splicing patterns from sequence alone [35]. First, we assess the distribution of splice-altering variants (SAVs) among all four archaic individuals, identify which genes are affected, and describe how the transcripts are modified. Second, we quantify which SAVs are also present in modern humans due to shared ancestry or introgression. Third, we quantify SAV enrichment among gene sets that underlie modern human phenotypes. Fourth, we estimate the effects of SAVs on the resulting transcript or protein. Fifth, we explore how selection shaped alternative splicing in archaics. Sixth, we evaluate the expression and function of archaic SAVs that are also present in modern humans. Finally, we highlight a handful of archaic SAVs with potential evolutionary significance.

## 2 Results

We examined the alternative splicing landscape in archaic hominins using all four currently available high-coverage archaic genomes, representing three Neanderthals [2–4] and a Denisovan [1]. We applied the SpliceAI classifier to sites with high-quality genotype calls where at least one archaic individual exhibited at least one allele different from the human reference (hg19/GRCh37) using the built-in GENCODE, Human Release 24, annotations to identify variants in gene bodies (**Fig. 1A**). SpliceAI estimates Δ, the splice-altering probability (SAP), for each variant of: 1) an acceptor gain, 2) an acceptor loss, 3) a donor gain, and 4) a donor loss (**Fig. 1A**). SpliceAI also indicates the positions changed for each of these four effects in basepairs.

**Fig. 1:**
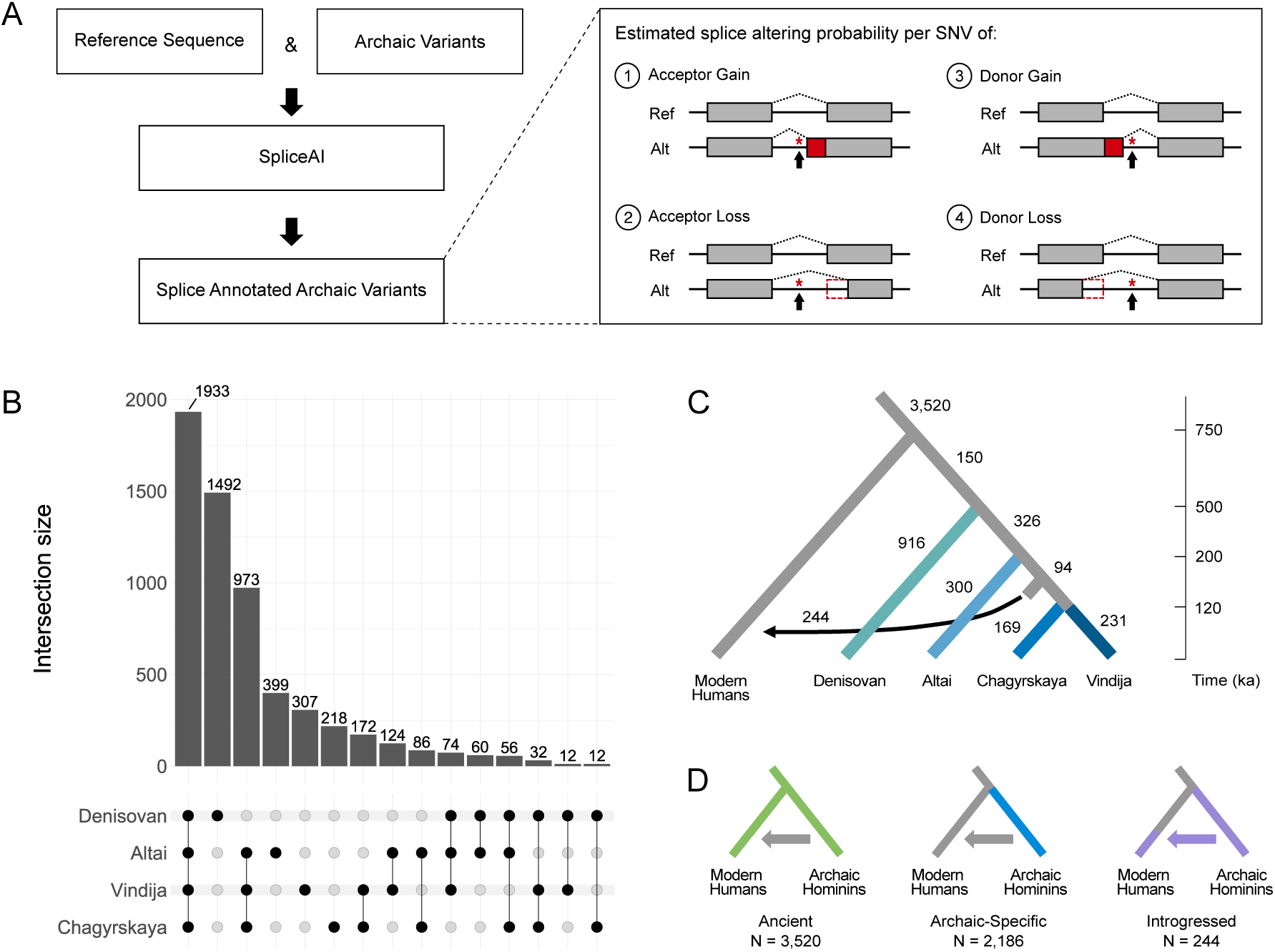
The identification, distribution, and origin of archaic SAVs. **(A)** We used SpliceAI to identify putative splice altering variants in archaic hominin genomes. We analyzed autosomal SNVs from four archaic hominins compared to the reference sequence (hg19/GRCh37). SpliceAI annotates each variant with splice altering probabilities (Δ scores) and position changes for each class of splicing alteration: 1) acceptor gain, 2) acceptor loss, 3) donor gain, or 4) donor loss. Here, we visualize one example consequence per splicing class alteration. See **Extended Data Fig. 7** for all possibilities. **(B)** The distribution of the presence of archaic SAVs across archaic individuals. The dot matrix indicates the number of SAVs per lineage(s). **(C)** The evolutionary origins of the archaic SAVs. From the distribution of SAVs across archaic and modern individuals, we inferred their origins using parsimony. We also identified introgressed archaic SAVs using two Neanderthal ancestry sets: [38] (shown here) and [39]. The divergence times and placement of the introgression arrow reflect estimates from [4] and [40]. We display data using Δ *≥* 0.2 here and these patterns are maintained when Δ *≥* 0.5. **(D)** We consider three main categories of archaic SAVs based on their origin and presence across populations. “Ancient” are archaic SAVs present in both modern and archaic hominin individuals and are inferred to have origins before the last common ancestor of these groups. “Archaic-Specific” are archaic SAVs that are present in archaic hominins, but absent or present at low frequency (allele frequency *<* 0.0001) in modern humans. “Introgressed” are archaic SAVs that were introgressed into Eurasian populations due to archaic admixture.

Alternative splicing occurs across nearly all eukaryotes and its molecular mechanisms are deeply conserved [41]. We therefore anticipated that the sequence patterns learned by SpliceAI in humans would generalize to archaics. To confirm this, we searched the 147 genes associated with the major spliceosome complex [42] for “archaic-specific” variants, i.e., archaic variants absent or at very low allele frequency (*<* 0.0001) from individuals sequenced by the 1000 Genomes Project (1KG) [43] and the Genome Aggregation Database (gnomAD) [44] (**Supplementary Data 1**). We annotated these variants using the Ensembl Variant Effect Predictor [45]. We found only two missense variants that were scored as likely to disrupt protein function by both PolyPhen and SIFT, neither of which were fixed in all four archaics (**Supplementary Data 1**). We observed a similar number of predicted deleterious variants in random sets of four diverse modern humans (0–3). Thus, there is near complete conservation of the proteins involved in splicing between archaic hominins and modern humans.

### 2.1 Thousands of splice altering variants (SAVs) are present in archaic hominins

We identified 1,567,894 autosomal positions with *≥* 1 non-reference allele among the archaic individuals (**Supplementary Table 1**). Many of these positions fell within a single GENCODE annotation; however, a handful were present in multiple annotated products (**Supplementary Table 2**). An individual variant that overlaps multiple annotations may have differential splicing effects on the different transcripts. Hereafter, we define a “variant” as one non-reference allele for a single annotated transcript at a given genomic position.

Among these variants, 1,049 had high splice altering probability (SAP; SpliceAI Δ *≥* 0.5) out of 1,607,350 archaic variants we analyzed. 5,950 archaic variants had moderate SAP (Δ *≥* 0.2). Hereafter, we refer to these variants as high-confidence splice altering variants (SAVs) and SAVs, respectively, and to maximize sensitivity, we focus on the SAVs in the main text.

The number of SAVs was similar among the four archaics, ranging from 3,482 (Chagyrskaya) to 3,705 (Altai) (**Supplementary Table 3**). These values fell within the range of SAVs observed in individual modern humans, estimated from one randomly sampled individual per 1KG population (**Supplementary Table 4**). SAVs were most commonly shared among all four archaic individuals (**Fig. 1B**; **Supplementary Fig. 1**). As expected from the known phylogeny, the Denisovan exhibited the most unique SAVs, followed by all Neanderthals, and each individual Neanderthal (**Fig. 1B**; **Supplementary Fig. 1**).

A total of 4,242 genes have at least one archaic SAV. 3,111 genes have only one SAV; however, 1,131 had multiple SAVs (**Supplementary Table 5**). Among the genes with the largest number of archaic SAVs are: *WWOX* (N = 9), which is involved in bone growth development [46], *HLA-DPA1* (N = 7) and *HLA-DPB1* (N = 10), essential components of the immune system [47], and *CNTNAP2* (N = 11), a nervous system protein associated with neurodevelopmental disorders that is also one of the longest genes in the human genome [48].

Many SAVs (47.8%) have a high SAP for only one of the four classes splicing change (acceptor gain, acceptor loss, donor gain, and donor loss) (**Supplementary Fig. 2**), and as expected, the overall association between the probabilities of different change types was negative (*ρ* = -0.34 – -0.14) for variants with at least one SAP greater than 0 (**Supplementary Fig. 3**). Donor gains were the most frequent result of splice altering variants for both thresholds (29.5% and 35.1% of variant effects, respectively) (**Supplementary Fig. 2**). While this may reflect the true distribution, we cannot rule out that the classifier has greater power to recognize donor gains compared to acceptor gains, acceptor losses, and donor losses.

### 2.2 37% of archaic SAVs are archaic-specific

We inferred the origin of archaic variants based on parsimony. We identified 2,186 (37%) “archaic-specific” SAVs. These archaic SAVs are absent among modern humans in 1KG and gnomAD or occur in gnomAD at a very low (*<* 0.0001) allele frequency (**Fig. 1C**). Such low frequency variants are likely to be recurrent mutations identical by state rather than identical by descent.

The remaining 63% of archaic SAVs are present in modern humans. Archaic hominins and modern humans last shared a common ancestor approximately 570–752 ka [40]. SAVs present in both archaic and modern humans may be the result of either introgression, shared ancestry, or recurrent mutation. To identify SAVs present in 1KG due to introgression, we used two datasets on archaic introgression into modern humans [38, 39]. While modern human genomes retain Denisovan and Neanderthal ancestry, most 1KG samples have minimal (*<* 1%) Denisovan ancestry [38, 39]. Therefore, we focused on Neanderthal introgression and classified 244 SAVs identified by [38] in 239 genes (**Fig. 1D**; **Supplementary Fig. 4**) and 386 SAVs identified by [39] in 361 genes as “introgressed” (**Supplementary Fig. 4**). Despite only modest overlap between the two introgression datasets (**Supplementary Fig. 5**), we observed qualitatively similar results in downstream analyses. Hereafter, we present results using the [38] introgressed variants in the main text and include results using the [39] set in the supplemental text.

Non-introgressed variants present in both archaic and modern humans likely evolved prior to our most recent common ancestor. We refer to these SAVs as “ancient”, and we consider the archaic SAVs with an allele frequency *≥* 0.05 in at least two 1KG superpopulations “high-confidence ancient”. This decreases the probability of recurrent mutation or misclassification of introgressed alleles. Hereafter, “ancient” refers to these high-confidence ancient variants unless otherwise specified. We identified 2,252 such variants based on [38] among 1,896 genes (**Fig. 1D**; **Supplementary Fig. 4**) and 2,195 variants based on [39] among 1,856 genes (**Supplementary Fig. 4**).

### 2.3 Archaic-specific SAVs are enriched in genes with diverse phenotypes

To identify the potential phenotypic consequences of archaic-specific SAVs, we tested for enrichment of functional annotations among genes with archaic-specific SAVs. Following [10], we considered links between genes and phenotypes from two sources: GWAS Catalog 2019 [49] and the Human Phenotype Ontology [50], that capture common and rare disease annotations, respectively. Structural properties of genes, such as the number of exons, influence the probability that they have SAVs (**Supplementary Table 6**; **Supplementary Fig. 6**). To account for these different probabilities, we generated a permutation-based empirical null distribution (Methods) and used it to estimate enrichment for each phenotype and control the false-discovery rate (FDR).

Given that we cannot directly observe archaic individuals, functions associated with genes with archaic-specific SAVs are of particular interest. We found enrichment for many phenotypes among the 1,907 genes with archaic-specific SAVs (**Fig. 2**; **Supplementary Data 2**). Only two common traits were significantly enriched for these SAVs: blood metabolite levels and blood metabolite ratios (**Fig. 2A**). There were substantially more rare phenotypes enriched among genes with archaic-specific SAVs (**Fig. 2B**), and these included traits that are known to differentiate archaic hominins and modern humans, including skeletal traits such as lumbar hyperlordosis and several cranial features (**Supplementary Data 2**). At least one significantly enriched trait occurred in every system across the Human Phenotype Ontology, except for the endocrine system.

**Fig. 2:**
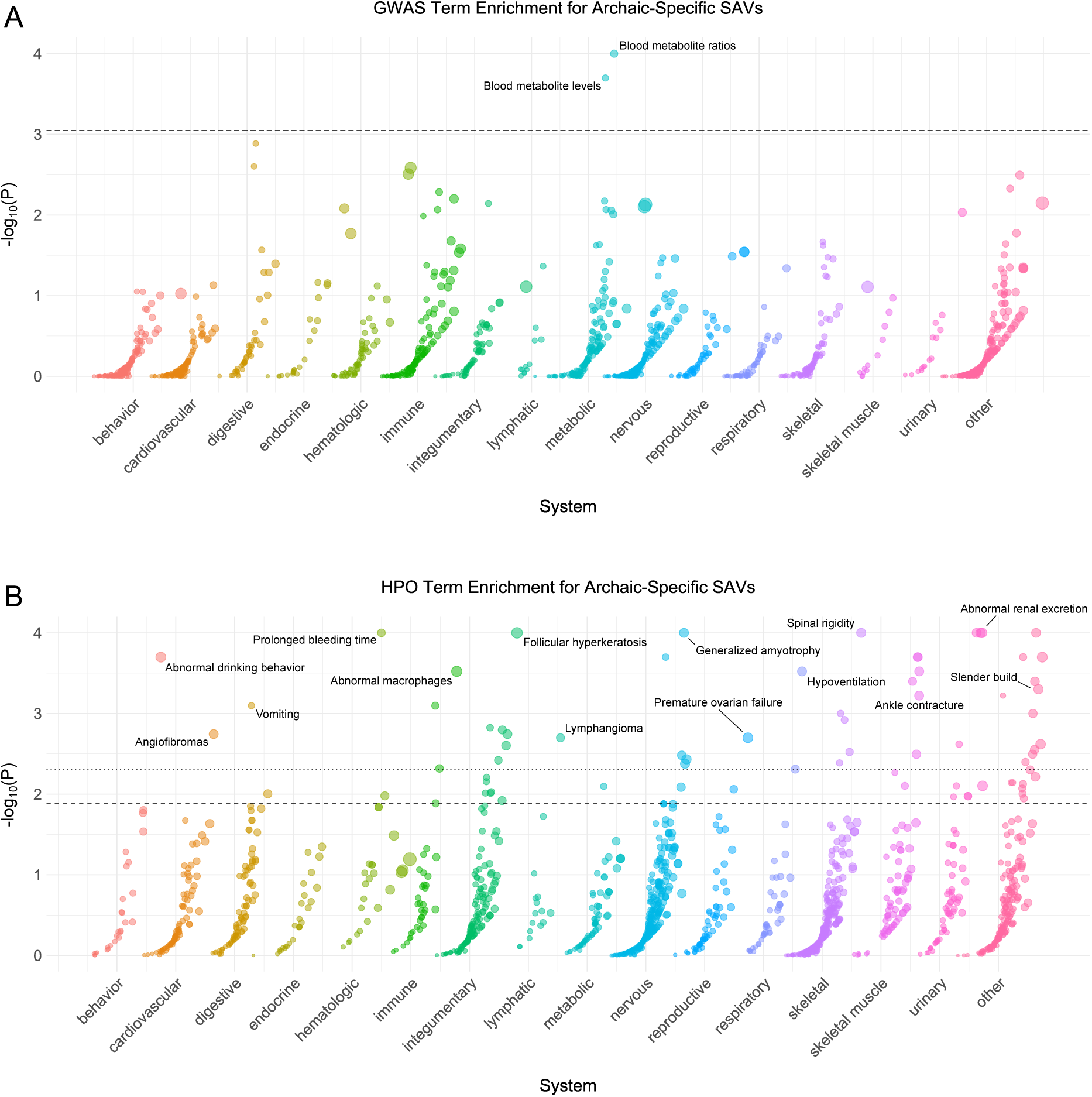
Genes with archaic-specific SAVs are enriched for roles in many phenotypes. **(A)** Phenotype associations enriched among genes with archaic-specific SAVs based on annotations from the 2019 GWAS Catalog. Phenotypes are ordered by increasing enrichment within manually curated systems. Circle size indicates enrichment magnitude. Enrichment and p-values were calculated from an empirical null distribution generated from 10,000 shuffles of maximum Δ across the entire dataset (Methods). Dotted and dashed lines represent false discovery rate (FDR) corrected p-value thresholds at FDR = 0.05 and 0.1, respectively. One example phenotype with a p-value *≤* the stricter FDR threshold (0.05) is annotated per system. **(B)** Phenotypes enriched among genes with archaic-specific SAVs based on annotations from the Human Phenotype Ontology (HPO). Data were generated and visualized as in **A**. See **Supplementary Data 2** for all phenotype enrichment results.

Next, we sought to characterize similarities and differences among the archaic hominin individuals. We assessed phenotype enrichment among genes that contained shared, Neanderthal-specific, and lineage-specific SAVs (**Supplementary Data 2**). We found minimal enrichment among the 106 genes with shared SAVs (**Extended Data Fig. 1**). However, there was considerable enrichment across various systems for Neanderthal- and lineage-specific SAVs (**Extended Data Figs. 2**–**6**). For example, all Neanderthals were enriched for SAVs in genes underlying skin conditions including abnormal skin blistering and fragile skin (**Extended Data Fig. 5**). The Denisovan exhibited enrichment for SAVs in genes associated with many skeletal and skeletal muscle system traits including skull defects, spinal rigidity, abnormal skeletal muscle fiber size, increased muscle fiber diameter variation, and type 1 muscle fiber predominance (**Extended Data Fig. 4**). No traits were enriched in multiple different sets of lineage-specific SAVs at FDR-adjusted significance levels.

### 2.4 Most SAVs result in isoforms that trigger nonsense mediated decay or yield altered transcripts and proteins

Each SAV can result in a number of effects on the mRNA product, including having little to no impact. Therefore, the above analysis captures the extent of potential phenotypic consequences as inferred using gene ontologies. Next, we sought to characterize the possible functional effects of archaic SAVs using an *in silico* approach.

We predicted the effect of each SAV on the resulting transcript by constructing a canonical transcript using the GENCODE exon annotations. Next, we generated a novel transcript using the variant, indicated splicing alteration class (e.g., acceptor gain), and Δ position for that alteration (**Extended Data Fig. 7**). If multiple alteration classes passed our SAP threshold, we modeled the class with the largest Δ. We compared the length and composition of the resulting transcripts and proteins for all but six SAVs with disagreements between the annotated transcript and genome sequences (**Supplementary Data 3**).

When considering the most likely effect per SAV, the majority (60%) of SAVs result in a different transcript or protein sequence (**Fig. 3A**). Among these consequential SAVs, the most prevalent effect was a longer protein that included premature termination codons (**Fig. 3A**). Many such isoforms would trigger nonsense mediated decay (NMD). The remaining SAVs resulted in altered transcripts or proteins that would not induce NMD but may yield a different or differentially stable protein.

**Fig. 3:**
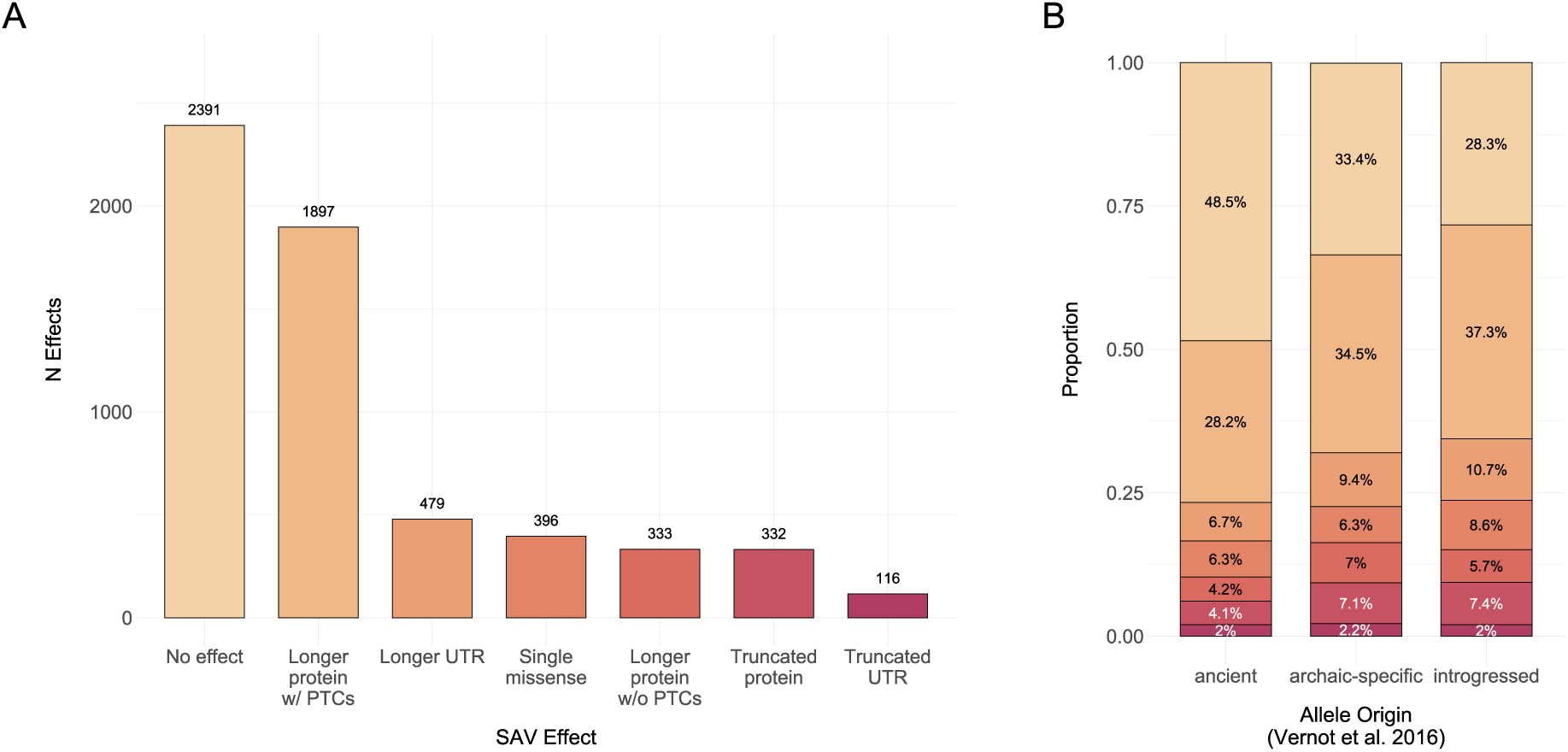
Most SAVs result in isoforms that trigger nonsense mediated decay or yield altered transcripts and proteins. **(A)** The number of SAVs that result in one of seven effects based on the single, largest splicing effect per SAV. We excluded six SAVs for which the genomic and transcriptomic annotations did not match. PTCs = premature termination codons, UTR = 5’ or 3’ untranslated region. **(B)** The number of protein effects stratified by allele origin [38]. Colors indicate the transcript or protein effect as in **A**.

When stratifying SAVs by allele origin, the proportion of these effects was generally similar for most classes (**Fig. 3B**). However, ancient SAVs more frequently resulted in no effect, whereas archaic-specific and introgressed SAVs were more likely to alter the canonical protein or UTRs (G Test of Independence, G = 138, P = 1.73 *×* 10*^−^*^23^). This pattern was also observed when variant allele origin was classified per [39] (G Test of Independence, G = 121, P = 3.45 *×* 10*^−^*^20^) (**Supplementary Table 7**). We also note that many SAVs are predicted to result in multiple splicing alteration classes (e.g., a donor gain and a donor loss), and thus by focusing on a single class per SAV, we may miss some biologically relevant effects.

### 2.5 Site-level evolutionary conservation varies across SAV origin

Genes vary in their tolerance to mutation and SAVs often disrupt gene function and contribute to disease [20–22]. To evaluate if the presence of archaic SAVs is associated with evolutionary constraint on genes, we compared the per gene tolerance to missense and loss-of-function variants from gnomAD [44] among ancient, archaic-specific, introgressed, and non-splice altered genes. In addition to constraint at the gene level, evolutionary constraint can be quantified at nucleotide level by methods like phyloP that quantify deviations from the expected neutral substitution rate at the site-level between species [51]. Thus, to explore the constraint on splicing altering variant sites themselves, we also compared their phyloP scores.

While we found a significant difference in the observed/expected number of missense variants per transcript among genes with different classes of SAV (Kruskal-Wallis, H = 18.079, P = 0.0004), the effect size was minimal (**Supplementary Fig. 7A**). Furthermore, genes with SAVs of different origins did not significantly differ in the observed/expected number of loss-of-function variants per transcript (Kruskal-Wallis, H = 1.533, P = 0.675) (**Supplementary Fig. 7B**). Variants classified per [39] exhibited the same pattern (**Supplementary Figs. 7C**, **7D**). These results suggest that genes with alternative splicing in archaics are similar in their gene-level constraint to other genes.

In contrast, phyloP scores were significantly different between ancient SAVs, archaic-specific SAVs, introgressed SAVs, and non-SAVs (Kruskal-Wallis, H = 877.429, P = 6.963 *×* 10*^−^*^190^) (**Fig. 4**). All of the variant sets exhibited a wide range of phyloP scores, indicating diverse pressures on SAVs of each type. However, ancient SAVs and non-SAVs exhibited largely negative phyloP scores suggesting substitution rates faster than expected under neutral evolution. In contrast, archaic-specific and introgressed SAVs had higher median phyloP scores suggesting that more of these loci experienced negative constraint. However, 84.3% occurred within the range consistent with neutral evolution (|phyloP| = 1.3). Variants classified per [39] exhibited similar patterns (**Supplementary Fig. 8**); however, archaic-specific, rather than introgressed, variants had a larger mean phyloP score.

**Fig. 4:**
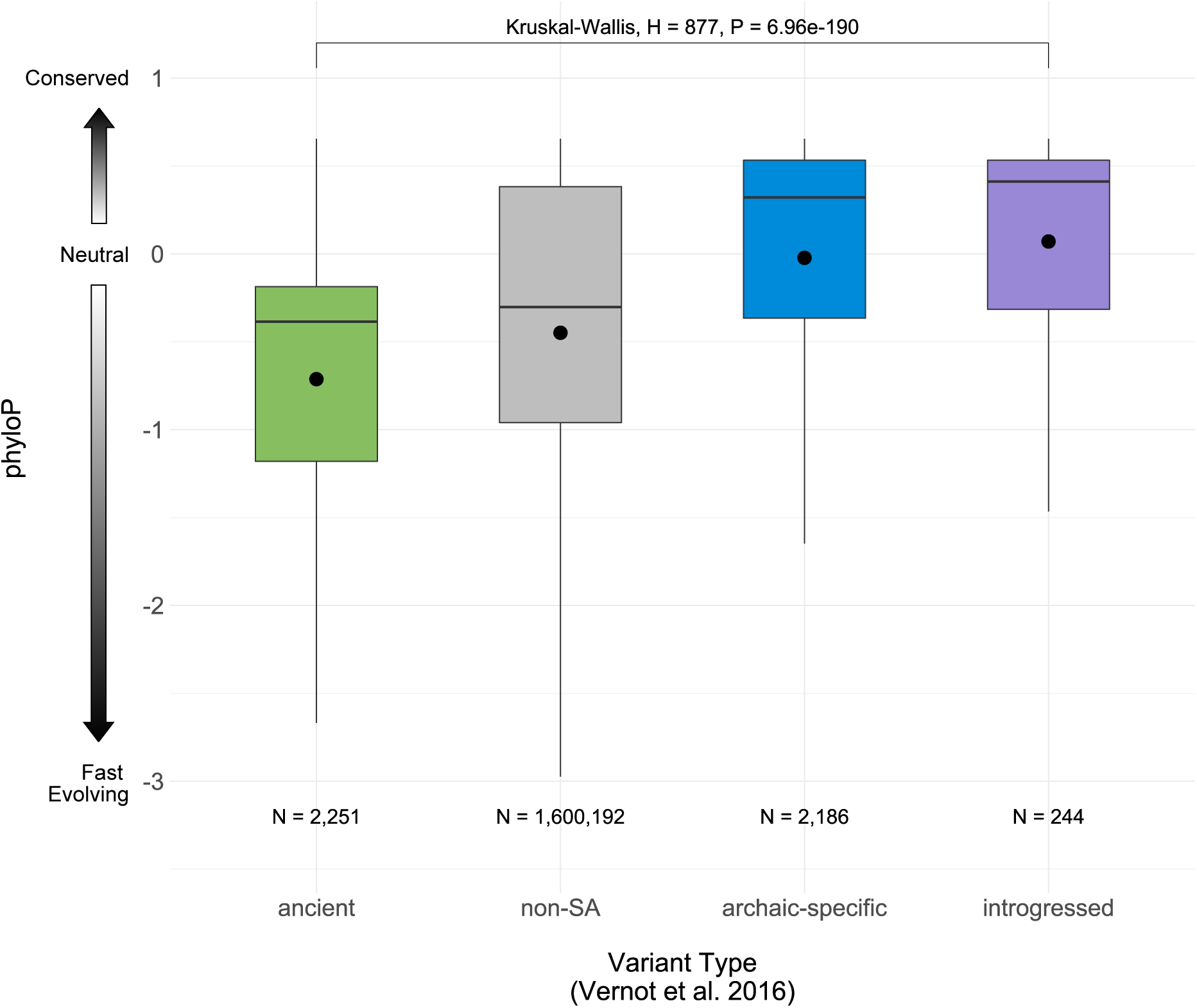
Site-level evolutionary conservation varies across SAVs with different origins. Evolutionary conservation score (phyloP) distributions for archaic SAVs of different origins and non-SAVs. Positive scores indicate substitution rates slower than expected under neutral evolution (conserved evolution), while negative scores indicate higher substitution rates than expected (fast evolution). The boxplots give the first quantile, median, and third quantile of the distributions, and the mean is noted by the black point. Ns are the number of variants per set. SAV classifications were based on [38] introgression calls.

### 2.6 The prevalence of SAVs across lineages is consistent with purifying selection on most SAVs

Variants that disrupt splicing and/or produce new isoforms are expected to be under strong negative selection [52, 53]. However, given differences in ages of archaic SAVs and the effective population sizes (*N_e_*) of the lineages in which they arose, different SAVs were likely exposed to different strengths of selection for different periods. Thus, we hypothesized that the probability a given SAV would survive to the present would vary based on its origin. For example, SAVs that arose in the ancestor all archaic lineages were likely subject to purifying selection over a longer time scale than lineage-specific SAVs, especially those that arose in lineages with low *N_e_*.

Shared archaic variants are depleted for SAVs compared to lineage-specific variants, and this depletion increased with higher splice altering probability thresholds (**Fig. 5A**, **Fig. 5B**, **Supplementary Table 8**). This result is consistent with the hypothesis that most splicing variants are deleterious and that the longer exposure to negative selection for older variants results in a smaller fraction of remaining SAVs. It is also with concordant with the site-level constraint results.

**Fig. 5:**
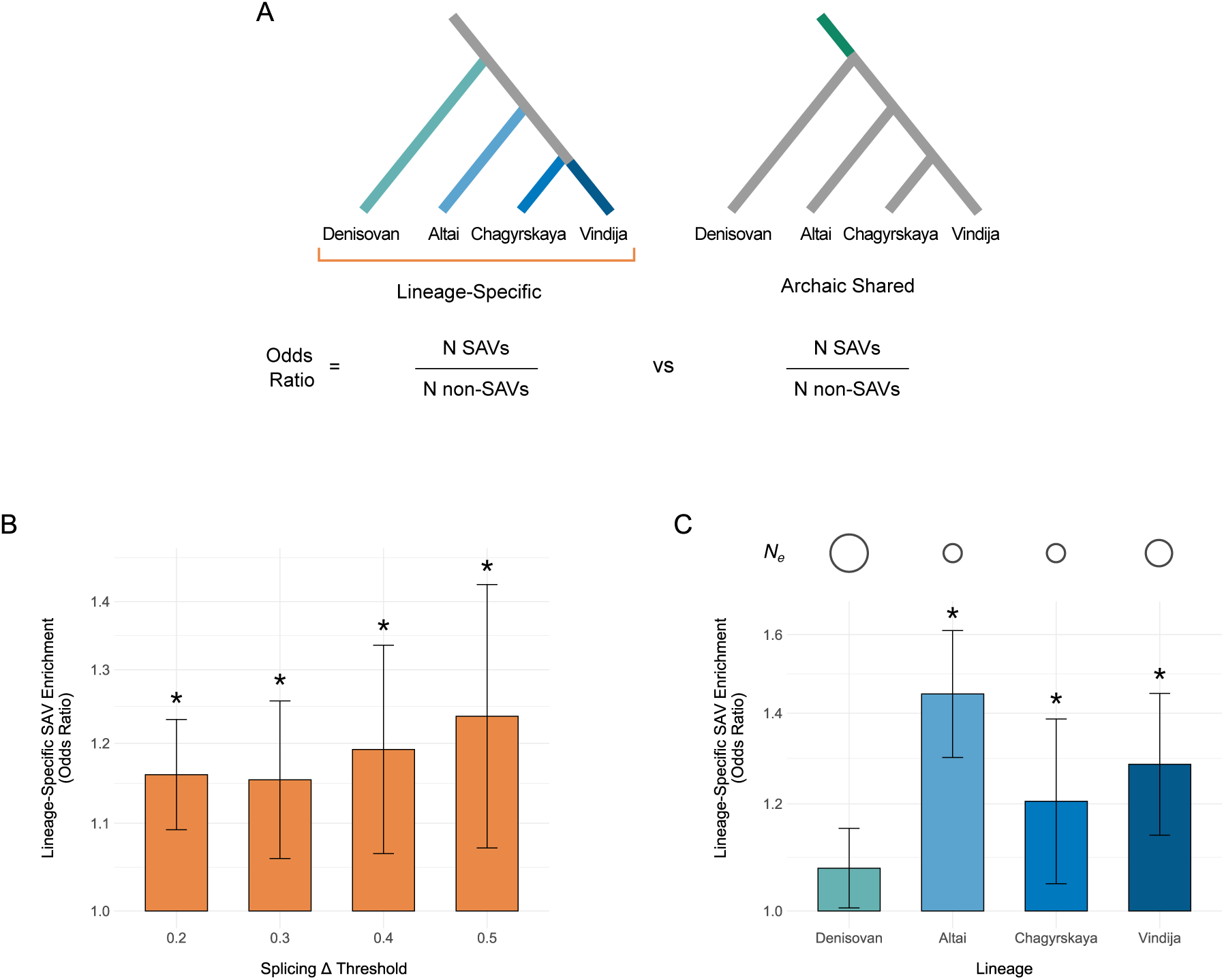
Lineage-specific archaic variants are enriched for SAVs compared to shared archaic variants. **(A)** We hypothesized that lineage-specific archaic variants (left) would be enriched for SAVs compared to variants shared among the four archaic individuals (right), due to less exposure to strong negative selection. To test this, we computed the odds ratio for being an SAV over all variants unique to each lineage (turquoise and blue edges) compared to variants shared among all four lineages (dark green edge). We also hypothesized that lineage-specific archaic variants would vary in their SAV enrichment compared to shared variants, based on the different effective population sizes and lengths of each branch. To test this, we computed the odds ratio for being an SAV over variants unique to each lineage individually compared to variants shared among all four lineages. **(B)** Archaic variants with origins in a specific archaic lineage are enriched for SAVs compared to variants shared among all four archaic lineages. The enrichment increases at increasing splice altering probability (Δ) thresholds. Note the y-axis is log10 transformed. **(C)** Lineage-specific archaic variants vary in their enrichment for SAVs. The Neanderthal lineage-specific variants have stronger SAV enrichment than Denisovan-specific variants. Estimated *Ne* per lineage is denoted by a circle above each lineage, with increasing size reflecting larger *Ne*. *Ne* estimates are from [4]. In all panels, asterisks reflect significance of Fisher’s exact tests using a Bonferroni corrected *α* (0.0125). Error bars denote the 95% CI. The number of variants used in each enrichment test are listed in **Supplementary Table 8** and Supplementary Table 9. Note the y-axis is log10 transformed.

Given that the population histories for each archaic lineage were likely different, we also compared within lineages. Neanderthals are thought to have lived in smaller groups and exhibited a lower *N_e_* than Denisovans [4]. We tested this hypothesis by repeating the SAV enrichment test for variants specific to each individual archaic lineage (**Fig. 5A**). All three Neanderthals were significantly enriched for unique SAVs compared to shared archaic variants after Bonferroni correction (OR = 1.205-1.447, **Fig. 5C**, **Supplementary Table 9**). In contrast, variants on the longer and higher *N_e_* Denisovan lineage were not significantly enriched for SAVs (OR = 1.075). At the stricter high-confidence SAV threshold, both the Altai and Vindija Neanderthals remained significantly enriched with increased odds ratios (**Supplementary Fig. 9**). These results are consistent with experimental results that found modern humans are depleted for SAVs with strong splicing effects compared to archaics [53].

### 2.7 Introgressed SAVs found in modern humans were present across archaics

We hypothesized that the evolutionary history of SAVs might also influence their prevalence in modern human populations. For example, introgressed variants experienced strong negative selection in the generations immediately after interbreeding [54], so archaic SAVs that survived stronger and longer-term selection would be more likely to survive in modern humans. To test this, we first considered the distribution of remaining introgressed variants among the archaics.

Most introgressed SAVs were present in all Neanderthals (N = 143) or present in all archaics (N = 68; **Supplementary Table 10**). No SAVs private to Vindija or Chagyrskaya nor shared between both late Neanderthals were identified as introgressed, even though Neanderthal ancestry in most modern humans is most closely related to Vindija and Chagyrskaya [3, 4]. This is consistent with weaker selection on lineage-specific SAVs and previous work suggesting that older introgressed archaic variants were more tolerated in humans [55–57].

To further test this, we compared the expected origin distribution for introgressed SAVs (based on the distribution of archaic-specific SAVs) to the observed distribution for introgressed SAVs. Fewer Altai-specific SAVs occur among introgressed variants whereas shared Neanderthal SAVs are more prevalent than expected (**Fig. 6A**). This pattern remains for highconfidence and [39] SAVs (**Supplementary Fig. 10**). These patterns suggest that older SAVs, either those that evolved prior to the Neanderthal common ancestor or prior to the Denisovan and Neanderthal common ancestor were the most tolerated after introgression.

**Fig. 6:**
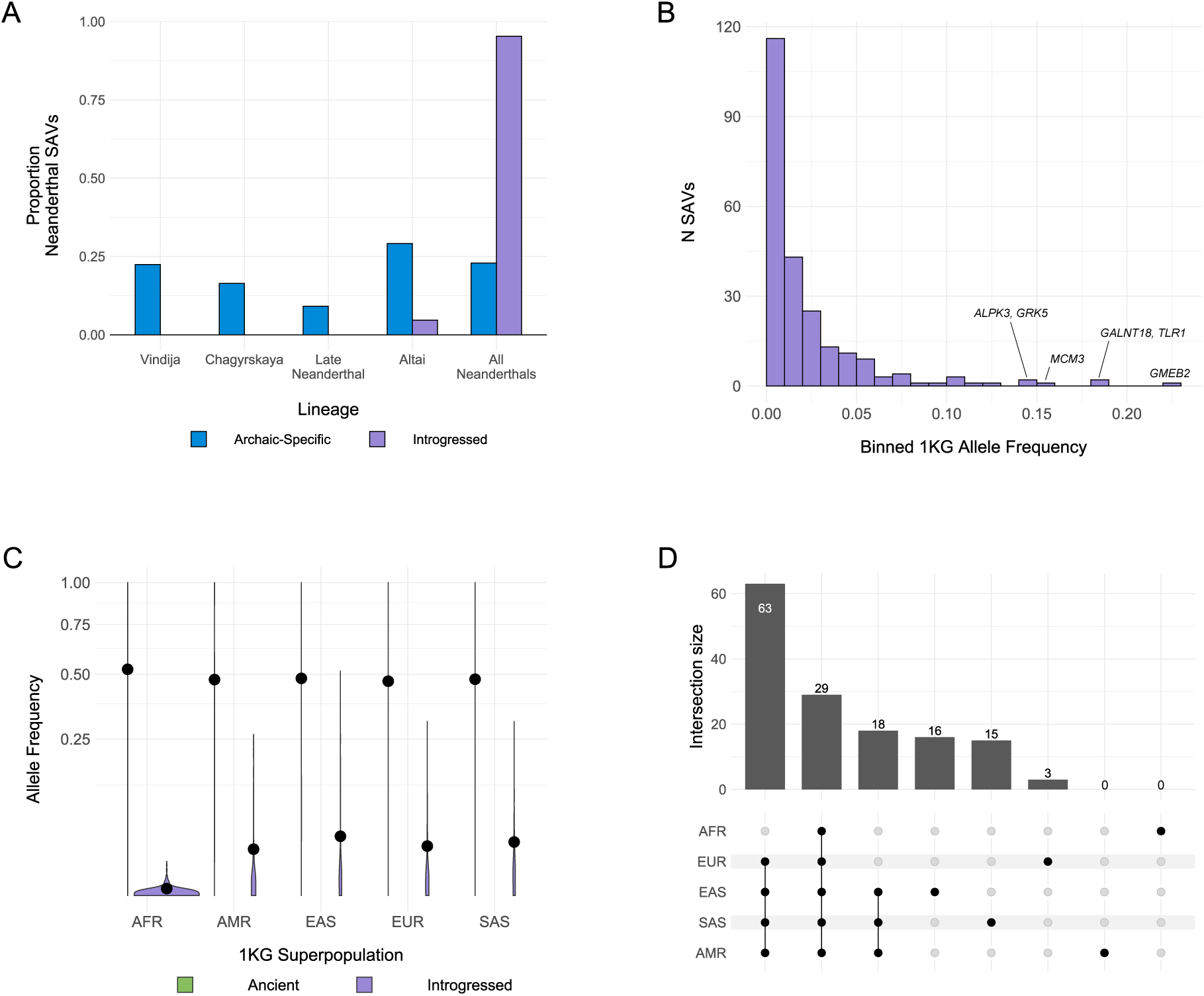
Introgressed SAVs present in modern humans were shared across archaic individuals. **(A)** Histograms comparing distributions of the presence of all Neanderthal SAVs (blue) and introgressed SAVs (purple) in different sets of Neanderthal individuals. Introgressed SAVs are significantly older than expected from all Neanderthal SAVs. We focused on Neanderthal lineages because of low power to detect introgressed Denisovan SAVs. All data are presented in **Supplementary Table 10**. **(B)** Allele frequency distribution for introgressed SAVs per [38] with Δ *≥* 0.2. Allele frequencies represent the mean from the 1KG African, American, East Asian, European, and South Asian superpopulation frequencies. **(C)** Allele frequency distributions for SAVs present in both archaic and modern individuals stratified by 1KG superpopulation and origin (ancient vs. introgressed). AFR = African, AMR = American, EAS = East Asian, EUR = European, SAS = South Asian. The black dot represents the mean allele frequency for each set. The y-axis is square root transformed. **(D)** The number of introgressed SAVs with a minimum allele frequency of at least 0.01 in each modern human population. We display all individual populations, the non-African set, and Asian/American set here. See **Supplementary** Fig. 15 for all sets. In all panels introgressed SAVs and frequencies are as defined by [38].

Consistent with known introgression patterns, introgressed SAVs occurred at lower overall frequencies (**Fig. 6B**). However, a small number of introgressed SAVs occur at modest to high frequencies among genes including *GMEB2*, *GALNT18*, and *TLR1* (**Fig. 6B**). The latter occurs in an adaptively introgressed locus spanning three toll-like receptors—key components of the innate immune system [58] and this SAV has been confirmed to generate an isoform using a massively parallel splicing reporter assay [53].

In contrast, ancient SAVs occur at high frequencies in all five 1KG superpopulations (**Supplementary Figs. 12**, **13**), and their frequencies are significantly higher among Africans (*µ* = 0.522) than non-Africans (*µ* = 0.476) (Mann-Whitney U, U = 10,963,956, P = 2.66 *×* 10*^−^*^09^) (**Fig. 6C**, **Supplementary Fig. 11**). Introgressed SAVs have significantly lower frequencies in all superpopulations (**Fig. 6B**) and are less likely to be shared among multiple populations (**Fig. 6C**, **Supplementary Figs. 14**, **15**).

It is possible these patterns reflect the general attributes of introgressed variants, rather than splicing effects of SAVs. We therefore examined the relationship between allele frequency in 1KG and splice altering probability (Δ max) for introgressed SAVs. 1KG populations did not generally differ in Δ max for either ancient or introgressed SAVs, although introgressed SAVs had a higher Δ max (**Supplementary Fig. 16**). We anticipated, however, introgressed SAVs that were more likely to be splice altering would occur at lower frequencies. Indeed, we found a significant negative association between allele frequency and Δ max for Δ *≥* 0.2 (Spearman, *ρ* = -0.2378, P = 0.0002) (**Extended Data Fig. 8**). This pattern is likely absent among ancient variants due to purifying selection on deleterious variants that occurred prior to the divergence of archaics and moderns. Further, our prediction that ancient SAVs were more likely to have no effect on the resulting transcript and protein (**Fig. 3B**) is consistent with this hypothesis.

### 2.8 Introgressed SAVs have functional associations

We tested whether any functional annotations (GWAS or HPO terms) were enriched among the 361 genes with [39] introgressed SAVs and 239 genes with [38] introgressed SAVs (**Supplementary Data 2**). Two terms were significantly enriched among genes with [38] introgressed SAVs: adverse response to breast cancer chemotherapy (GWAS) and Oligohydramnios (HPO) (**Extended Data Fig. 9**). Four HPO terms related to hip-girdle, pelvic, and shoulder muscles were enriched among genes with [39] introgressed SAVs (**Extended Data Fig. 10A**). However, 19 GWAS terms met our FDR-corrected significance threshold including *H. pylori* serologic status and systemic sclerosis (**Extended Data Fig. 10A**). Overall, these results suggest that SAVs surviving in modern human populations influence several immune, skeletal, and reproductive phenotypes.

We further considered the potential functional effects of introgressed SAVs by intersecting them with Neanderthal variants exhibiting allele-specific expression (ASE) in modern humans [59]. We identified 16 SAVs out of 1,236 ASE variants, including variants in *GSDMC*, *HSPG2*, and *RARS* (**Supplementary Table 11**). The SAV in *HSPG2*, predicted to create a donor gain, was recently validated using a massively parallel splicing reporter assay [53]. We also note that a handful of the [59] ASE variants fell just under our SAV threshold. Among these is a Neanderthal variant (rs950169) in *ADAMTSL3* that results in a truncated protein [59]. SpliceAI correctly predicted the loss of the upstream acceptor (Δ = 0.19), although it did not indicate the downstream acceptor gain.

### 2.9 Introgressed SAVs are more tissue-specific than ancient SAVs

Given their different histories of selective pressures, we hypothesized that introgressed SAVs would be more tissue-specific than ancient SAVs in their effects. To explore this, we identified 1,381 archaic SAVs with sQTL data from GTEx across 49 tissues.

Introgressed sQTL SAVs were significantly associated with tissue specific gene expression compared to ancient sQTL SAVs (Mann-Whitney U, U = 35,123.5, P = 0.027) (**Fig. 7A**). On average, introgressed SAVs influenced splicing in 4.92 fewer tissues than ancient SAVs. Further, all sQTL SAVs with broad effects (*>* 40 tissues) were ancient (107 high-confidence and 5 low-confidence). 74 of these were shared among all four archaics (**Supplementary Table 12**), suggesting that core sQTL SAVs were more likely to evolve in the deep past. These patterns were also observed among the [39] variants (**Supplementary Fig. 17**). Collectively, 30% of sQTL SAVs (N = 427) were associated with tissue specific effects on splicing (1 or 2 tissues) (**Supplementary Fig. 18**). Consistent with known gene expression patterns, testis had the most unique sQTL among SAVs, followed by skeletal muscle and thyroid.

**Fig. 7:**
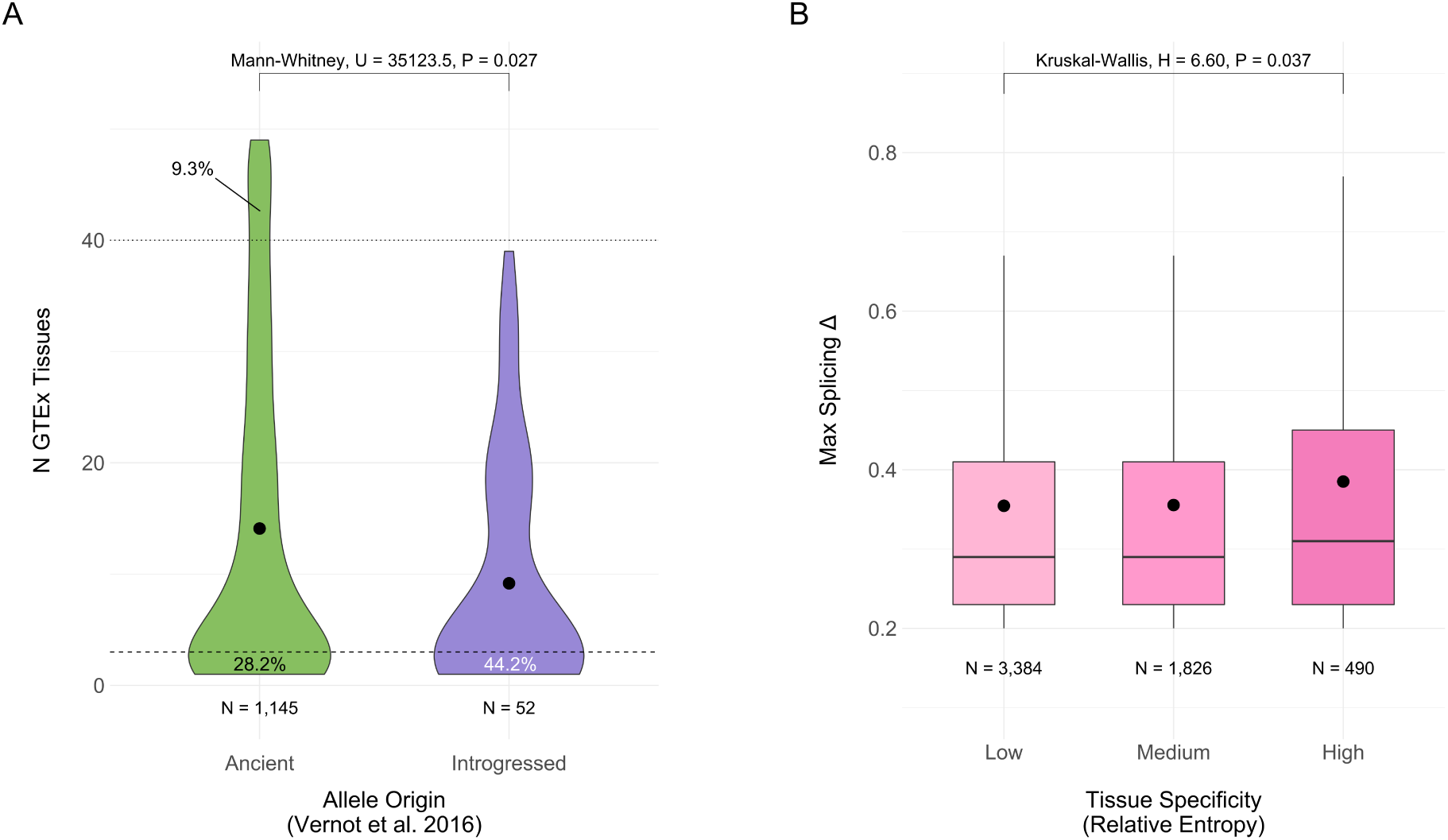
Increased tissue-specificity is associated with introgressed SAVs and with increased probability of alternative splicing. **(A)** The distribution of the number of GTEx tissues in which an ancient or introgressed SAV (per [38]) was identified as an sQTL. Introgressed variants are significantly more tissue-specific. We defined “tissue-specific” variants as those occurring in 1 or 2 tissues and “core” sQTLs as those occurring in > 40 of the 49 tissues. The dashed and dotted lines represent these definitions, respectively. The proportion of SAVs below and above these thresholds are annotated for each allele origin. **(B)** The distributions of maximum splice altering probability (Δ) for SAVs in 4,061 genes binned by tissue-specificity of expression. We quantified the tissue-specificity of each gene as the relative entropy of expression levels across 34 tissues from GTEx compared to genes overall. Low tissue specificity reflects relative entropy *≤* 0.1, medium tissue specificity reflects relative entropy *>* 0.1 and *≤* 0.5, and high tissue specificity reflects relative entropy *>* 0.5 (**Supplementary** Fig. 19).

Variation in gene expression among tissues may also influence the efficacy of negative selection to remove deleterious SAVs. For example, [60] demonstrated that more ubiquitously expressed genes in *Paramecium tetraurelia* exhibited less alternative splicing compared to genes with more tissue-specific expression due to the differences in the efficacy of negative selection. We predicted that tissue-specificity of expression would be associated with the number of SAVs per gene or maximum Δ. We quantified tissue-specificity of expression using the relative entropy of the TPM count for each gene across tissues compared to the expression distribution across tissues for all genes in GTEx. This metric ranges from 0 to 1, with higher values reflecting greater tissue specificity. Most genes exhibited broad expression (**Supplementary Fig. 19**), so we divided genes into low, medium, and high tissue-specificity based on the relative entropy.

Genes with the most tissue-specific expression had significantly higher median maximum splice altering probability (Δ) than genes with broader expression patterns (**Fig. 7B**; Kruskal-Wallis, H = 6.599, P = 0.037). This could indicate greater selection against SAVs likely to influence expression across many tissues; however, we note that the effect was small in magnitude. The distribution of the number of archaic SAVs per gene did not differ significantly between entropy classes (Kruskal-Wallis, H = 1.89, P = 0.388) (**Supplementary Fig. 20**); all had a median of 1 SAV.

### 2.10 Archaic SAVs with potential evolutionary significance

Many archaic SAVs influence genes with known or previously hypothesized significance to the evolutionary divergence between archaic hominins and modern humans. For example, the 2’-5’ oligoadenylate synthetase *OAS* locus harbors an adaptively introgressed splice altering variant at chr12: 113,357,193 (G *>* A) that disrupts an acceptor site and results in multiple isoforms and leads to reduced activity of the antiviral enzyme [61, 62]. This ancestral variant was reintroduced in to modern Eurasian populations by Neanderthal introgression [63]. SpliceAI correctly predicted the acceptor loss at this site (Δ = 0.89). This locus harbors 92 additional archaic variants (N = 92). We found one additional SAV at chr12: 113,355,275 in *OAS1* that potentially results in an acceptor gain (Δ = 0.26). This allele was unique to the Denisovan, is derived, and was present in only one of 2,054 1KG samples as a heterozygote. This suggests potential further splice variant evolution of this locus, with possible Denisovan-specific effects.

We also identified several variants at other well-studied loci. Variation in human populations at the *EPAS1* locus includes a Denisovan-introgressed haplotype thought to contribute to adaptation to living at high altitude among Tibetans [64]. Of 184 archaic variants occurring at this locus, we identified two as candidate SAVs. One variant (chr2: 46,584,859; rs372272284) is fixed in the Denisovan whereas all Neanderthals have the human reference allele (**Fig. 8A**). The variant is introgressed and present at low frequency in East Asians in 1KG, and is also the lead variant in an observed association of the introgressed haplotype with decreased hemoglobin levels in Tibetans [65]. This SAV strengthens a canonical 5’ splice site (CAA|GT to CAG|GT) [29], resulting in a donor gain (Δ = 0.37) (**Fig. 8A**). If used, this splice site would introduce multiple stop codons, resulting in NMD (**Supplementary Data 3**). This would result in the same molecular effect (decreased circulating *EPAS1* RNA) that is thought to contribute to hypoxia adaptation [66]. The other candidate SAV (chr2: 46,610,904) is absent from 1KG/gnomAD and occurs as a heterozygyote in the Altai Neanderthal, and is near the end of the last intron of the gene, making it much less likely to fundamentally alter the mRNA product.

**Fig. 8:**
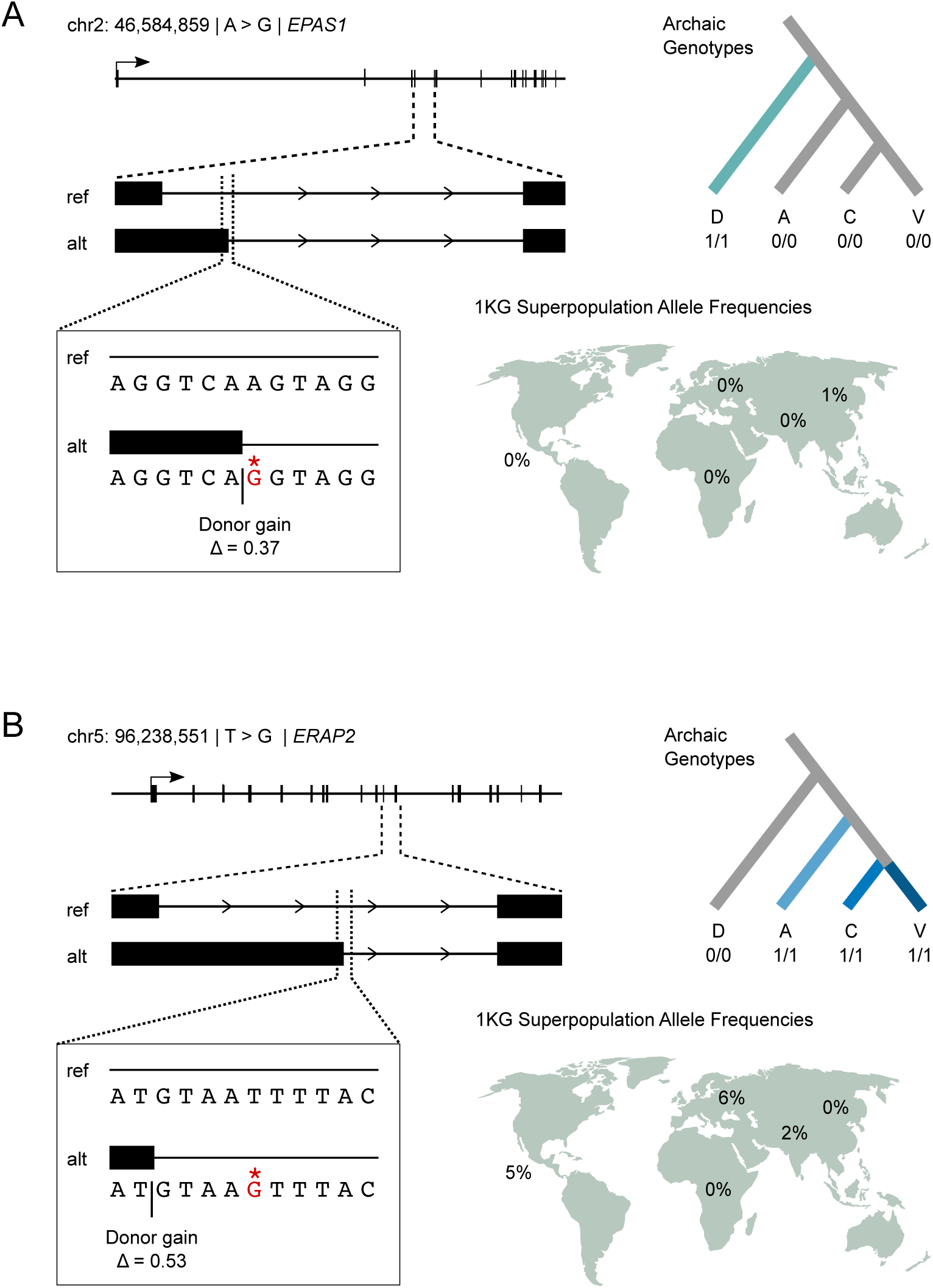
Example archaic SAVs leading to nonsense mediated decay in loci with evidence of recent adaptive evolution. **(A)** A Denisovan-specific homozygous SAV results in a donor gain in *EPAS1*, hypoxia-inducible factor-2alpha, between the fourth and fifth exon. The transcript resulting from the SAV introduces six premature termination codons (**Supplementary Data 3**), which likely results in the elimination of the transcript via nonsense mediated decay. This SAV potentially contributes to the effects of the introgressed haplotype in Tibetan adaptation to living at high altitude. This variant is present as a heterozygote in 12 individuals from 1KG: 8 from the East Asian superpopulation and 4 from the South Asian superpopulation. **(B)** Three archaic SAVs, including a Neanderthal-specific variant, occur in *ERAP2*, a MHC presentation gene with evidence of strong balancing selection in human populations. Consistent with this, the SNV occurs at low frequency in three of the five 1KG superpopulations. As in the *EPAS1* example, this variant results in a donor gain between the 11th and 12th exons, which introduces nine premature termination codons **Supplementary Data 3**.

We also identified three archaic SAVs within *ERAP2*, a gene subject to strong and consistent balancing selection in different human populations [67]. SpliceAI correctly identified a previously characterized human variant (chr5: 96,235,896; rs2248374), which causes a donor loss (Δ = 0.51) and results in a truncated protein and subsequent NMD of the mRNA. However, we identified an additional Neanderthal SAV, which is also introgressed and occurs at low frequencies among Americans (5%), Europeans (6%), and South Asians (2%) in 1KG (**Fig. 8B**). This SAV, rs17486481, is a donor gain (Δ = 0.53) that introduces a canonical 5’ splice site (AT|GTAAT to AT|GTAAG) and would similarly result in NMD (**Fig. 8B**) (**Supplementary Data 3**). However this allele always occurs with the non-truncated version of rs2248374 (while being much rarer), and the need to maintain the non-truncated allele is most likely why it remains uncommon. The third variant (chr5: 96,248,413) was archaic-specific—occurring as a heterozygote in the Altai Neanderthal—and results in an acceptor gain (Δ = 0.24).

## 3 Discussion

Alternative splicing plays a critical role in organismal biology, particularly during development and establishing tissue identity [14]. Thus, alternative splicing often contributes to adaptation and phenotypic differences between closely related species [23–28]. The development of machine learning algorithms that can predict alternative splicing from sequence alone now enables analysis of alternative splicing in populations for which transcriptomic data are difficult or impossible to generate, including archaic hominins. Here, we use SpliceAI to uncover the previously unobservable genome-wide alternative splicing landscape of archaic hominins.

We identify thousands of putative splice altering variants from the high-coverage genomes of three Neanderthals and a Denisovan. We find that many of these variants do not occur in modern humans, and we propose that they are implicated in specific phenotypic differences between archaic hominins and modern humans. Additionally, many SAVs are shared with modern humans and are ancient, evolving before the common ancestor of archaic hominins and modern humans. Furthermore, a few hundred SAVs are present in human populations due to introgression, and these surviving introgressed SAVs are almost entirely shared across Neanderthals. We also observe multiple lines of evidence supporting the role of negative selection in shaping SAV patterns.

Given that introgressed and ancient SAVs are present in modern humans, their splicing patterns have the potential to be directly studied to further understanding of their phenotypic effects. We found that 36.7% of non-archaic-specific SAVs were identified in modern humans in GTEx as sQTLs. There are several reasons why archaic SAVs might not have been detected as sQTL. Splicing is often tissue-specific, and GTEx assayed only a small fraction of tissues and contexts. Furthermore, splicing is influenced by sequence, but is also a somewhat stochastic process influenced by other cellular dynamics, such as polymerase pausing [29]. These, along with limited statistical power in many GTEx tissues, particularly for SAVs at low frequency in Europeans, mean that many splice-altering variants should not necessarily be detected. Indeed, we observe higher fractions of SAVs as sQTL when analyzing high frequency variants (**Supplementary Fig. 21**). The tissue-specificity of archaic SAVs is of great interest, but the degree to which SAV tissue-specificity in modern humans reflects specificity in archaic hominins is unknown without further experimental study. However, such studies are challenging because the genomic and archaic cellular context cannot be perfectly replicated (i.e., testing an archaic SAV in a Neanderthal genome background in a Neanderthal tissue) [5, 68].

Our results offer new insight into an essential molecular mechanism and previously unstudied attributes of archaic hominins; however, we note some limitations of our approach. First, we did not include structural variants (InDels) or variants from the sex chromosomes in this analysis, both of which warrant further study. For example, the X chromosome exhibits high levels of alternative splicing [69] and splicing can occur in a sex-specific manner [26, 70]. However, for now, we only have high-coverage archaic hominin sequences from females. Future development and application of models with sex-specific transcriptomic data may offer additional phenotypic insights. Second, the tag SNPs and modern human samples used in this analysis are best suited to identifying Neanderthal rather than Denisovan introgression [38, 39]. Our conservative approach for identifying introgressed haplotypes means that the number of introgressed SAVs reported here is an underestimate and does not include Denisovan-derived SAVs. Multiple modern human populations contain considerable Denisovan ancestry, therefore, future work should consider these variants.

In summary, our approach of combining machine learning with ancient DNA and modern population genetic data identifies thousands of archaic variants that potentially alter splicing, including many that appear to be specific to archaic hominins. Genes affected by archaic SAVs are enriched for roles in a variety of phenotypes and several influence loci with known relevance to recent human evolution. For example, two archaic SAVs that we highlight likely cause in NMD of the resulting *EPAS1* and *ERAP2* transcripts. Downregulation of *EPAS1* is thought to underlie high-altitude adaptation in Tibetans [66]. In *ERAP2*, another variant in human populations that induces NMD has experienced strong balancing selection [67]. These examples underscore that phenotypic effects from alternative splicing are not limited to expanded proteomic diversity, but also downregulation of gene expression via NMD [71, 72]. Further work is needed to understand the functional effects of these and other archaic SAVs. [53] recently used a massively parallel splicing reporter assay to assess exonic SAVs in archaics and modern humans, validating a number of predictions from the present study. However, this assay is limited to testing only a subset of exonic variants and additional assays are required to test the other types of exonic and any intronic SAVs [53]. Nonetheless, our results suggest that alternative splicing played a role in hominin divergence and offers specific molecular hypotheses for testing. The identification of archaic-specific splice variants here will enable future analysis of human-specific splice variants. We also anticipate that our sequence-based approach will enable study of alternative splicing in other extinct or difficult to sample taxa.

## 4 Methods

### 4.1 Archaic Genomic Data

We retrieved four high-coverage publicly available archaic hominin genomes representing three Neanderthals [2–4] and a Denisovan [1].

We excluded sites that were invariant among the archaic individuals (“ALT = .”) and variants with low site quality (“QUAL *<* 30”). Further, low quality genotypes were set to missing based on read depth (“FMT/DP *<* 10”) and genotype quality (“FMT/GQ *<* 30”). We also normalized InDels and split multi-allelic records into separate entries for positions with multiple variants (“norm -m -”). All filtering was completed using bcftools, version 1.13 [73].

All genomic coordinates presented in the manuscript and supplementary material refer to hg19/GRCh37.

### 4.2 Variant Annotation

We annotated variants for putative alternative splicing using SpliceAI, version 1.3.1 [35]. Briefly, SpliceAI uses a deep residual neural network to estimate the splice altering probability and position change of each variant from DNA sequence alone making it ideal for studying archaic hominins, for which we cannot obtain transcript-level data. The model considers 5 kbp flanking the variant in both directions. The output includes four splice altering probabilities (Δs) for 1) acceptor gain (AG), 2) acceptor loss (AL), 3) donor gain (DG), and 4) donor loss (DL) as well as the position changes for each of the four deltas. Δs range from 0 to 1 and represent the likelihood a variant is splice altering for one or more of the four categories. We implemented SpliceAI in a Conda package using keras, version 2.3.1 [74] and tensorflow, version 1.15.0 [75]. After filtering, we ran SpliceAI on sets of 5 *×* 10^3^ variants using the hg19 reference genome using the GENCODE, Human Release 24, annotations [76] included with the package. We concatenated all results and further split variants with multiple annotations. Among all variants, 32,105 exhibited multiple annotations with different effects on splicing (**Supplementary Table 2**). While we included InDels and variants on the X chromosome in this scan, we restricted all downstream analyses to autosomal SNVs (Discussion).

### 4.3 Defining SAVs

For each variant, we identified the maximum SAP (Δ) among all four classes: AG, AL, DG, and DL. We then defined SAVs using two Δ thresholds: Δ max *≥* 0.2 and 0.5, “SAVs” and “high-confidence SAVs”, respectively.

We determined whether the number of SAVs identified in each archaic individual was different than expected by randomly selecting a sample from 24 1KG populations. We annotated all variants present among these individuals using SpliceAI and the hg38 annotations included with the package. We then analyzed the variants as for the archaics (i.e., splitting multi-allelic sites and variants with multiple GENCODE annotations). We filtered for variants with a Δ max *≥* 0.2 and summed the number of variants per 1KG sample that had at least one alternate allele present.

### 4.4 Archaic Variants in Modern Humans

We noted the distribution of each variant among the archaics using eight classes: 1) Altai, 2) Chagyrskaya, 3) Denisovan, 4) Vindija, 5) Late Neanderthal (Chagyrskaya and Vindija), 6) Neanderthal (Altai, Chagyrskaya, and Vindija), 7) Shared (all four archaics) and 8) Other (all remaining possible subsets). The assignment was based on the presence of at least one allele with an effect.

We assessed whether any variants present among the archaics are also present in modern humans using biallelic SNVs and InDels from 1KG, Phase 3 [43] and SNVs from gnomAD, version 3 [44]. We used LiftOver [77] to convert archaic variants from hg19 to hg38. We then normalized variants (“norm -m - -f hg38.fa”) and subset variants to those within gene bodies (“view -R genes.bed”). We queried these variants for allele count, allele number, and allele frequencies (“query -f”). Further, for 1KG variants, we retrieved allele frequency per 1KG superpopulation: Africa (AFR), Americas (AMR), East Asia (EAS), Europe (EUR), and South Asia (SAS). These precomputed values had been rounded to two decimal places in the VCFs. Normalization, filtering, and querying was done using bcftools [73]. After using LiftOver to convert back to hg19 coordinates, we merged the 1KG and gnomAD variants with the archaic variants ensuring the archaic and modern reference and alternate alleles matched. We recalculated the 1KG allele frequency for Africans as the annotated frequency included samples from an admixed African population: African ancestry in SW USA (ASW). We subset samples from Esan (ESN), Mandinka (GWD), Luhya (LWK), Mende (MSL), and Yoruban (YRI) and calculated allele frequency as allele count divided by allele number per site.

We used two datasets to identify introgressed variants: [38] and [39]. These datasets differ in their approach to recognizing introgressed sequences and partly overlap the archaic variants considered in this study. [38] used the S* statistic to classify human sequences as introgressed. S* leverages high linkage-disequilibrium among variants in an admixed target population that are absent in an unadmixed reference population [78, 79]. Introgressed haplotypes are then identified by maximizing the sum of scores among all SNP subsets at a particular locus [78, 79]. Tag SNPs are those variants that match an archaic allele and occur with at least two other tag SNPs in a 50 Kb window. Haplotypes were defined as regions encompossing *≥* 5 tag SNPs in LD within a given human population (*R*^2^ *≥* 0.8). We collated tag SNPs from all four populations: East Asian (ASN), European (EUR), Melanesian (PNG), and South Asian (SAS). We retained all metadata from [38]. A handful of tag SNPs encompass multiple haplotypes that reflect differences in haplotype size between modern human populations; we retained the first record per variant. [39] developed a modified S* statistic, Sprime, which employs a scoring method that adjusts the score based on the local mutation and recombination rates, allows for low frequency introgression in the unadmixed outgroup, and avoids windowing to identify introgressed segments. We collated introgressed variants for 20 non-African populations and filtered for those that matched the Altai Neanderthal at high-quality loci.

A handful of sequences in the hg19 reference genome are introgressed from archaic hominins. Therefore, we maximized the number of introgressed sites we could analyze by defining sites rather than variants as introgressed if either the reference or alternate allele for each SAV matched any Neanderthal base at a matching position. We ensured that SpliceAI predictions were similar for these allele pairs, regardless of which was the reference and alternate, by generating a custom hg19 sequence where introgressed reference alleles (N = 7,977) from [38] were replaced by the alternate allele using a custom script. We then applied SpliceAI to the introgressed reference alleles, now considered to be the alternate. We found that 24 of the 26 variants were classified as SAVs (**Supplementary Table 13**). One of the remaining two variants was nearly identical in splicing probability (Δ max = 0.19 and Δ max = 0.2), whereas the other variant’s predictions were different (Δ max = 0.16 and Δ max = 0.31) (**Supplementary Table 13**). Given this overall similarity, we maintained the original predictions for introgressed [38] reference alleles in our dataframe but provide the predictions when these nucleotides are the alternate allele for all SAVs and non-SAVs in the project GitHub repository. We recalculated allele frequencies for all introgressed variants to account for sites where the reference sequence contained introgressed alleles, as the precomputed 1KG allele frequency would be incorrect. The [39] metadata designate whether the reference or alternate allele is introgressed. Therefore, we used the 1KG allele frequency for sites with an introgressed alternate allele and subtracted the 1KG allele frequency from 1 for sites with an introgressed reference allele. For [38] introgressed variants, we calculated an average from the metadata, which included the allele frequencies in various populations for the introgressed allele. We took the mean of the AFR, AMR, EAS, EUR, and SAS frequencies for all introgressed positions.

The presence of introgressed alleles in the human reference results in previously excluded human polymorphisms due to our filtering criteria. We quantified these potential ancient or introgressed SAVs by intersecting sites that were fixed among the archaics for the human reference with introgressed alleles from [39] and [38] using bedtools2, version 2.30 [80]. We repeated the above procedure, inserting the alternate alleles from intersected sites into the hg19 reference and running SpliceAI on the reference alleles formatted as alternate alleles. This yielded 12,003 variants from 11,833 positions, of which 41 variants had a Δ max *≥* 0.2. We do not include these variants in the main text but the SpliceAI predictions for all SAVs and non-SAVs from this set are available in the project GitHub repository.

We categorized each variant’s “origin” based on presence in 1KG and gnomAD as well as whether or not the variant was introgressed. Further, we classified each variant’s allele origin based on introgressed variants identified by [38] “Vernot allele origin” and [39] “Browning allele origin” due to the incomplete overlap among variants in those datasets. Variants that did not occur in 1KG or gnomAD were defined as “archaic-specific”. Low frequency variants in modern humans are also highly likely to be the result of recurrent mutation rather than shared ancestry. In support of this hypothesis, we found CpGs were enriched among rare variants (allele frequency < 0.0001) vs non-CpG common variants (allele frequency *≥* 0.01) (Fisher’s exact test, OR = 1.88, p < 0.0001). Therefore, we also designated gnomAD variants whose allele frequency was *<* 0.0001 as “archaic-specific”. Sample sizes in 1KG and GTEx do not permit this level of sensitivity; therefore, allele frequency was only considered for gnomAD variants. Variants that were present in 1KG or gnomAD at an allele frequency *≥* 0.0001 and introgressed were defined as “introgressed”. Variants that were present in 1KG or gnomAD at an allele frequency *≥* 0.0001 but not introgressed were considered “ancient” at two confidence levels. “High-confidence ancient” variants were present in at least two 1KG superpopulations at an allele frequency *≥* 0.05, while “low-confidence ancient” variants did not meet this threshold. We report analyses on the high-confident ancient set; this helps to remove cases of potential convergent mutation. We did not retrieve population-level allele frequency data for gnomAD variants; therefore, common variants present in gnomAD and absent from 1KG were classified as “low-confidence ancient”. We restricted analyses including allele frequency as a variable to 1KG variants with population-level allele frequencies.

### 4.5 Gene Characteristics, Mutation Tolerance, and Conservation

We used the SpliceAI annotation file for hg19 from GENCODE, Human Release 24 [76], to count the number of exons per gene and calculate the length in bp of the gene body and the coding sequence. The number of isoforms per gene were retrieved from GENCODE, Human Release 40. We retrieved missense and loss-of-function (LoF) observed/expected ratios from gnomAD [44] to quantify each gene’s tolerance to mutation. We also considered conservation at the variant level. We used the primate subset of the 46 way multi-species alignment [51]. Positive phyloP scores indicate conservation or slower evolution than expected, whereas negative phyloP scores indicate acceleration or faster evolution than expected based on a null hypothesis of neutral evolution.

### 4.6 Phenotype Enrichment

We followed the approach of [10] to assess enrichment for SAVs in genes implicated in different human phenotypes. Many gene enrichment analyses suffer from low power to detect enrichment because an entire genome is used as the null distribution. Relatedly, SAVs are unevenly distributed throughout archaic genomes. We addressed this issue by generating a null distribution from the observed data. We first retrieved phenotypes and the associated genes per phenotype from Enrichr [81–83]. We used both the 2019 GWAS Catalog and the Human Phenotype Ontology (HPO). The GWAS Catalog largely considers common disease annotations and has 1,737 terms with 19,378 genes annotated [49], whereas HPO largely considers rare disease annotations and has 1,779 terms with 3,096 genes annotated [50]. All 3,516 terms were manually curated into one of 16 systems: behavioral, cardiovascular, digestive, endocrine, hematologic, immune, integumentary, lymphatic, metabolic, nervous, other, reproductive, respiratory, skeletal, skeletal muscle, and urinary.

We considered nine different gene sets, generated using SAVs with Δ *≥* 0.2, for our enrichment analyses: 1) genes with lineage-specific Altai SAVs (N = 283), 2) genes with lineagespecific Chagyrskaya SAVs (N = 165), 3) genes with lineage-specific Denisovan SAVs (N = 859), 4) genes with lineage-specific Vindija SAVs (N = 228), 5) genes with SAVs present in all three Neanderthals (N = 227), 6) genes with SAVs shared among all four archaics (N = 106), 7) genes with all archaic-specific SAVs (N = 1,907), 8) genes with introgressed SAVs per [38] (N = 239), and 9) genes with introgressed SAVs per [39] (N = 361). The shared set only included variants present in all four archaics and excluded those that were inferred from parsimony. We retained duplicated gene names to reflect genes with multiple SAVs.

We identified which genes were present in both the GWAS Catalog and HPO per set using a boolean to calculate the observed gene counts per term per ontology. We then removed GWAS and HPO terms per set that did not include at least one gene from the set. This resulted in 631 Altai, 1,407 archaic-specific, 761 Browning et al. 2018 introgressed, 412 Chagyrskaya, 1,023 Denisovan, 515 Neanderthal, 295 shared, 627 Vernot et al. 2016 introgressed, and 474 Vindija terms for the 2019 GWAS Catalog and 622 Altai, 1,490 archaic-specific, 720 Browning et al. 2018 introgressed, 391 Chagyrskaya, 1,152 Denisovan, 528 Neanderthal, 306 shared, 651 Vernot et al. 2016 introgressed, and 522 Vindija terms for the Human Phenotype Ontology. The max Δ was then shuffled across all 1,607,350 variants without modifying the annotation, allele origin, or distribution data. The distribution of genes for both ontologies was then recorded. We repeated this process 1 *×* 10^4^ times per set and calculated enrichment as the number of observed genes divided by the mean empirical gene count per term. p-values were calculated as the proportion of empiric counts + 1 *≥* the observed counts + 1. We adjusted our significance level due to multiple testing by correcting for the false discovery rate (FDR). We used a subset (N = 1 *×* 10^3^) of the empirical null observations and selected the highest p-value threshold that resulted in a *V* /*R < Q* where *V* is the mean number of expected false discoveries and *R* is the observed discoveries [10]. We calculated adjusted significance levels for each set for *Q* at both 0.05 and 0.1.

### 4.7 Novel Transcripts and Proteins

We constructed a novel transcript per SAV to assess downstream effects on the resulting protein. We generated a canonical transcript per gene using the exons defined from GENCODE. Next, we constructed a novel transcript using the splicing alteration class (acceptor gain, acceptor loss, donor gain, or donor loss) and associated position information per SAV. For SAVs with multiple splicing class alterations, e.g., a variant that results in both an acceptor gain and an acceptor loss, we modeled the alteration class per SAV with the largest Δ.

For acceptor and donor gains, we identified the relevant exon and added or removed sequence based on the SpliceAI prediction (**Extended Data Fig. 7**). For acceptor losses, we removed the subsequent exon from the transcript if the effect occurred at the upstream exon boundary (**Extended Data Fig. 7**). Similarly, we added the intronic sequence to the canonical transcript for donor losses that occurred at the exon boundary (**Extended Data Fig. 7**). For each of these scenarios, we included the variant when appropriate but otherwise kept the canonical sequence. We then retrieved a single start codon position from the associated GFF, prioritizing the longest and experimentally-validated (Ensembl) vs. computationally-predicted (HAVANA) start sites when there were multiple start positions.

We compared the canonical and novel transcripts and the resulting protein (**Supplementary Data 3**). We assigned each SAV as resulting in one of seven effects based on transcript/protein length and composition: 1) the canonical and novel transcripts/proteins are identical (“no effect”), 2) the novel protein is longer than the canonical and includes premature termination codons (“longer w/ PTCs”), 3) the novel protein is longer and does not include premature termination codons (“longer w/o PTCs”), 4) the 5’ or 3’ untranslated region (UTR) is longer than the canonical transcript, 5) the novel protein is the same length as the canonical but contains a single missense variant (“single missense”), 6) the novel protein is shorter than canonical protein (“truncated protein”), and 7) the 5’ or 3’ UTR is truncated than the canonical transcript (“truncated UTR”).

### 4.8 Gene Expression, Tissue Specificity, and sQTL

We used TPM counts for each gene from GTEx, version 8, to analyze expression. We quantified tissue specificity as the relative entropy of each gene’s expression profile across 34 tissues compared to the median across genes overall. Thus, a gene with expression only in a small number of tissues would have high relative entropy and a gene with expression across many tissues would have low relatively entropy to this background distribution. The 34 tissues were selected based on groupings from the Human Protein Atlas to minimize the amount of sharing between distinct tissues, e.g., since brain tissues are over-represented in GTEx. We used median expression across tissues as the null and calculated relative entropy using the entropy function from the SciPy statistics package [84]. Based on the observed distribution of relative entropy scores (**Supplementary Fig. 19**), we designated genes with scores *≤* 0.1 as “low tissue-specificity”, genes with scores > 0.1 and *≤* 0.1 as “medium tissue-specificity”, and genes with scores > 0.5 as “high tissue-specificity”. We compared both the number of SAVs per gene and the maximum Δ for SAVs among the three relative entropy categories.

We downloaded sQTL data from GTEx, version 8. We collated significant variant-gene associations (N = 24,445,206) across all 49 tissues and intersected these with SAVs, using LiftOver [77] to convert the SAVs to hg38 and then back to hg19 after intersecting.

### 4.9 Major Spliceosome Complex

We characterized differences in the major spliceosome complex between archaics and modern humans by identifying missense variants in the 147 genes associated with the complex. We identified 1,746 variants that did not occur in 1KG nor gnomAD or were present at low frequency in gnomAD (i.e., “archaic-specific”). We ran these variants through the Ensembl Variant Effect Predictor (VEP) [45] using the GRCh37.p13 assembly and all default options.

We repeated the above analysis on all four archaics and the randomly sampled 1KG individuals (Defining SAVs) using all variants that occurred in the spliceosome genes and analyzed them with VEP using the appropriate assembly.

### 4.10 Analysis

All data analyses were performed using Bash and Python scripts, some of which were implemented in Jupyter notebooks. We used samtools, version 1.16, to index custom FASTAs [85]. We employed non-parametric tests to analyze data including Fisher’s exact test, Kruskal-Wallis tests, Mann-Whitney U tests, and Spearman correlation, implemented with SciPy [84]. Partial correlations were run using the Pigouin package, version 0.5.2 [86]. Some additional metrics were calculated using custom functions. All reported p-values are two-tailed.

### 4.11 Visualization

Results were visualized using Inkscape, version 1.1 [87] and ggplot, version 3.3.6 [88] implemented in R, version 4.1.2 [89]. Additional packages used to generate figures include complexupset, version 1.3.3 [90]; cowplot, version 1.1.1; eulerr, version 6.1.1 [91]; reshape2, version 1.4.4 [92]; and tidyverse, version 1.3.2 [93].

### 4.12 Data availability

We used publicly available data for all analyses. Archaic InDel data are from the following repository: http://ftp.eva.mpg.de/neandertal/Vindija/VCF/indels/. Archaic SNV data are from the following repositories: Altai Neanderthal (http://ftp.eva.mpg.de/neandertal/Vindija/VCF/Altai/), Chagyrskaya (http://ftp.eva.mpg.de/neandertal/Chagyrskaya/VCF/), Denisova (http://ftp.eva.mpg.de/neandertal/Vindija/VCF/Denisova/), and Vindija (http://ftp.eva.mpg.de/neandertal/Vindija/VCF/Vindija33.19/). Modern human data are from the Thousand Genomes Project (http://hgdownload.soe.ucsc.edu/gbdb/hg38/1000Genomes/) and gnomAD (https://gnomad.broadinstitute.org/downloads#v3-variants). Introgressed tag SNPs from [38] were retrieved from: https://drive.google.com/drive/folders/0B9Pc7_zItMCVM05rUmhDc0hkWmc?resourcekey=0-zwKyJGRuooD9bWPRZ0vBzQ. Introgressed variants from [39] were retrieved from: https://data.mendeley.com/datasets/y7hyt83vxr/1. gnomAD constraint data were retrieved from: https://storage.googleapis.com/gcp-public-data--gnomad/release/2.1.1/constraint/gnomad.v2.1.1.lof_metrics.by_gene.txt.bgz. phyloP data for the primate subset were retrieved from: http://hgdownload.soe.ucsc.edu/goldenPath/hg19/phyloP46way/primates.phyloP46way.bw. ASE variants were retrieved from: https://drive.google.com/file/d/10ebWfA-sboAL1SDplmIrvH-xK4x9iohV/view. TPM data were retrieved from the Human Protein Atlas (https://www.proteinatlas.org/download/rna_tissue_gtex.tsv.zip). sQTL data were retrieved from GTEx, version 8 (https://storage.googleapis.com/gtex_analysis_v8/single_tissue_qtl_data/GTEx_Analysis_v8_sQTL.tar). Genes associated with the major spliceosome complex were retrieved from the HUGO Gene Nomenclature Committee (https://www.genenames.org/data/genegroup/#!/group/1518). The compiled dataset used in our analyses is available on Dryad (DOI: 10.7272/Q6H993F9).

### 4.13 Code availability

All code used to conduct analyses and generate figures is publicly available on GitHub (https://github.com/brandcm/Archaic_Splicing).

## 4.14 Acknowledgements

We thank Mary Lauren Benton for kindly sharing data on tissue specificity and Ziyue Gao for helpful discussion on recurrent mutations. Evonne McArthur and David Rinker provided comments that improved this manuscript. We also thank members of the Capra Lab for feedback on figures. This research greatly benefited from access to the Wynton high-performance compute cluster at UCSF. L.L.C. was funded by National Institutes of Health grant T32HG009495 to the University of Pennsylvania. J.A.C. and C.M.B. were funded by National Institutes of Health grant R35GM127087.

## 4.15 Author Contributions

Conceptualization, C.M.B, L.L.C, and J.A.C.; Formal Analysis, C.M.B and L.L.C; Writing – Original Draft, C.M.B, L.L.C, and J.A.C.; Writing – Review & Editing, C.M.B, L.L.C, and J.A.C.

## 4.17 Extended Data Figures

**Extended Data Fig. 1.**
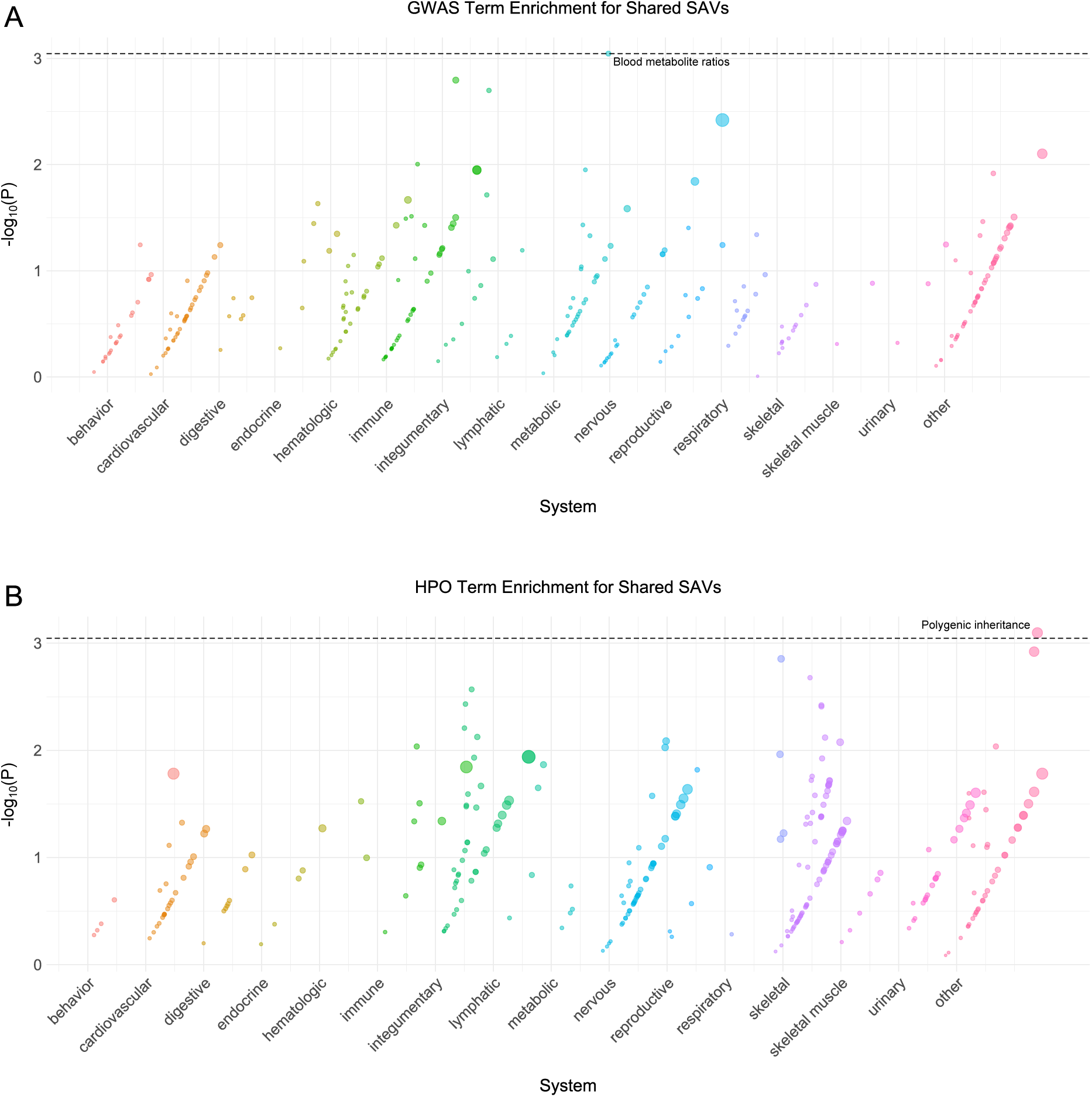
Shared phenotype enrichment. **(A)** Phenotype associations enriched among genes with archaic-specific shared SAVs based on annotations from the 2019 GWAS Catalog. Phenotypes are ordered by increasing enrichment within manually curated systems. Circle size indicates enrichment magnitude. Enrichment and p-values were calculated from an empirical null distribution generated from 10,000 shuffles of maximum Δ across the entire dataset (Methods). Dotted and dashed lines represent false discovery rate (FDR) corrected p-value thresholds at FDR = 0.05 and 0.1, respectively. At least one example phenotype with a p-value *≤* the stricter FDR threshold (0.05) is annotated per system. **(B)** Phenotypes enriched among genes with archaic-specific shared SAVs based on annotations from the Human Phenotype Ontology (HPO). Data were generated and visualized as in **A**. See **Supplementary Data 2** for all phenotype enrichment results.

**Extended Data Fig. 2.**
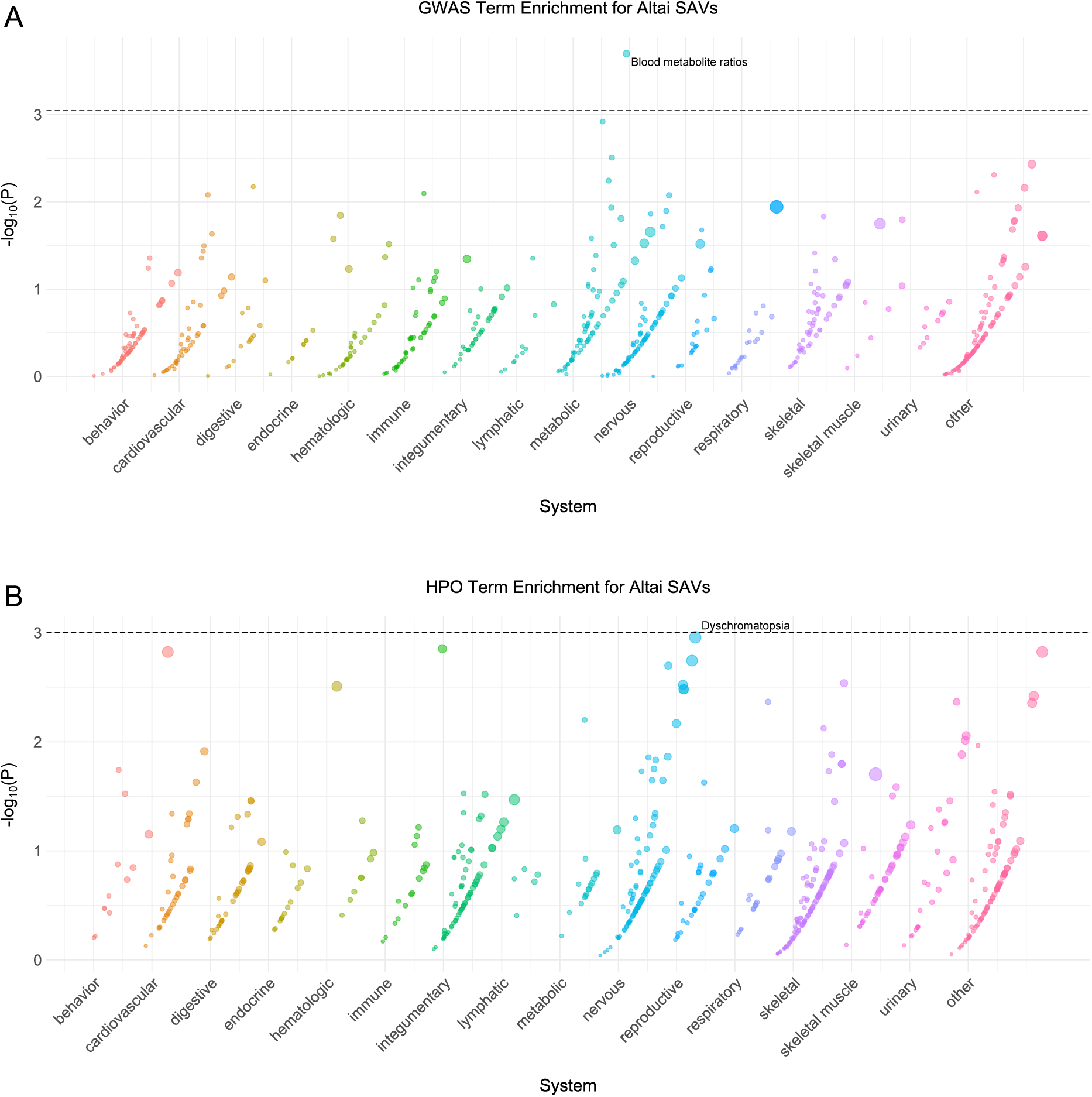
Altai phenotype enrichment. **(A)** Phenotype associations enriched among genes with archaic-specific Altai SAVs based on annotations from the 2019 GWAS Catalog. Phenotypes are ordered by increasing enrichment within manually curated systems. Circle size indicates enrichment magnitude. Enrichment and p-values were calculated from an empirical null distribution generated from 10,000 shuffles of maximum Δ across the entire dataset (Methods). Dotted and dashed lines represent false discovery rate (FDR) corrected p-value thresholds at FDR = 0.05 and 0.1, respectively. At least one example phenotype with a p-value *≤* the stricter FDR threshold (0.05) is annotated per system. **(B)** Phenotypes enriched among genes with archaic-specific Altai SAVs based on annotations from the Human Phenotype Ontology (HPO). Data were generated and visualized as in **A**. See **Supplementary Data 2** for all phenotype enrichment results.

**Extended Data Fig. 3.**
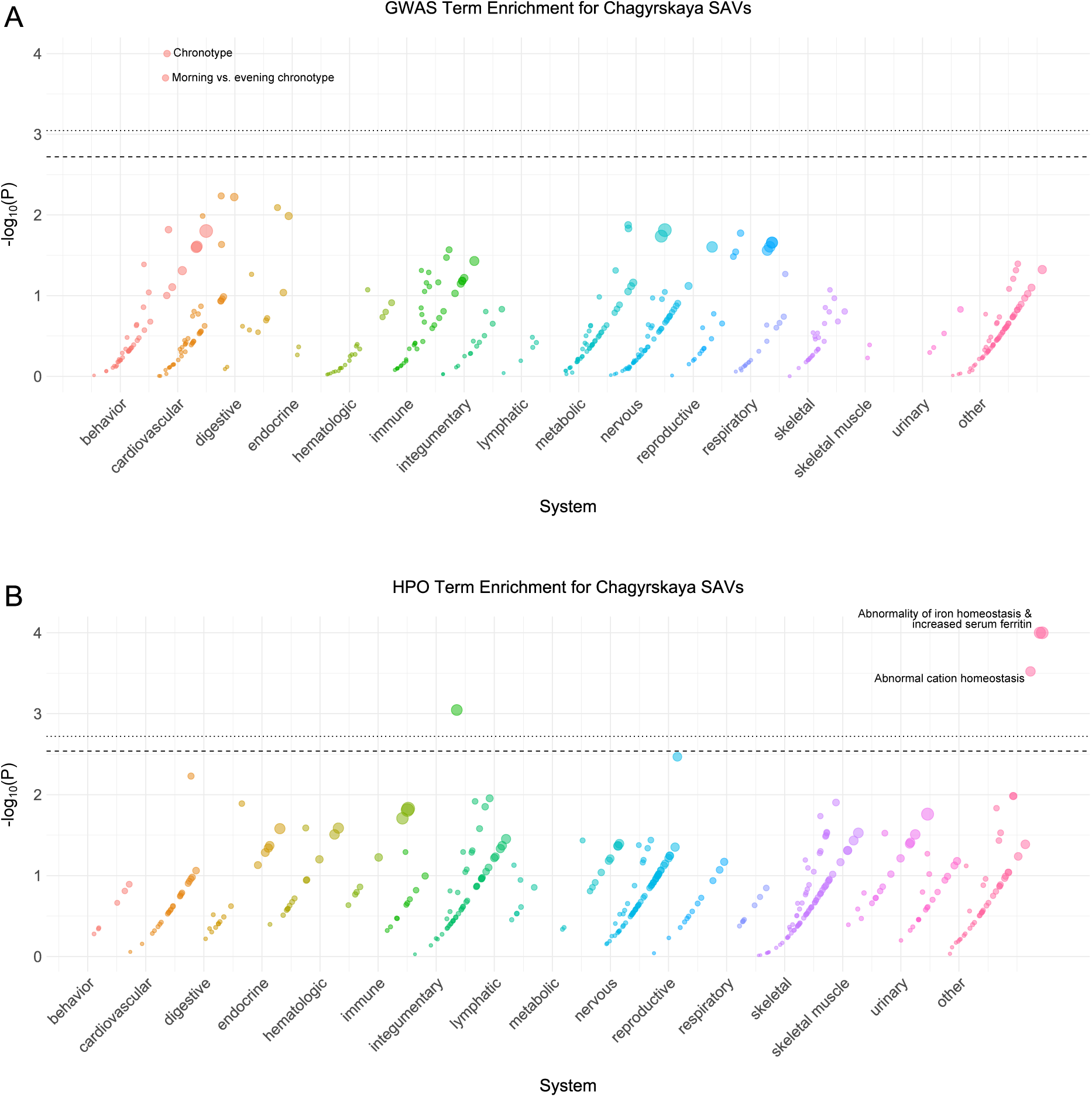
Chagyrskaya phenotype enrichment. **(A)** Phenotype associations enriched among genes with archaic-specific Chagyrskaya SAVs based on annotations from the 2019 GWAS Catalog. Phenotypes are ordered by increasing enrichment within manually curated systems. Circle size indicates enrichment magnitude. Enrichment and p-values were calculated from an empirical null distribution generated from 10,000 shuffles of maximum Δ across the entire dataset (Methods). Dotted and dashed lines represent false discovery rate (FDR) corrected p-value thresholds at FDR = 0.05 and 0.1, respectively. At least one example phenotype with a p-value *≤* the stricter FDR threshold (0.05) is annotated per system. **(B)** Phenotypes enriched among genes with archaic-specific Chagyrskaya SAVs based on annotations from the Human Phenotype Ontology (HPO). Data were generated and visualized as in **A**. See **Supplementary Data 2** for all phenotype enrichment results.

**Extended Data Fig. 4.**
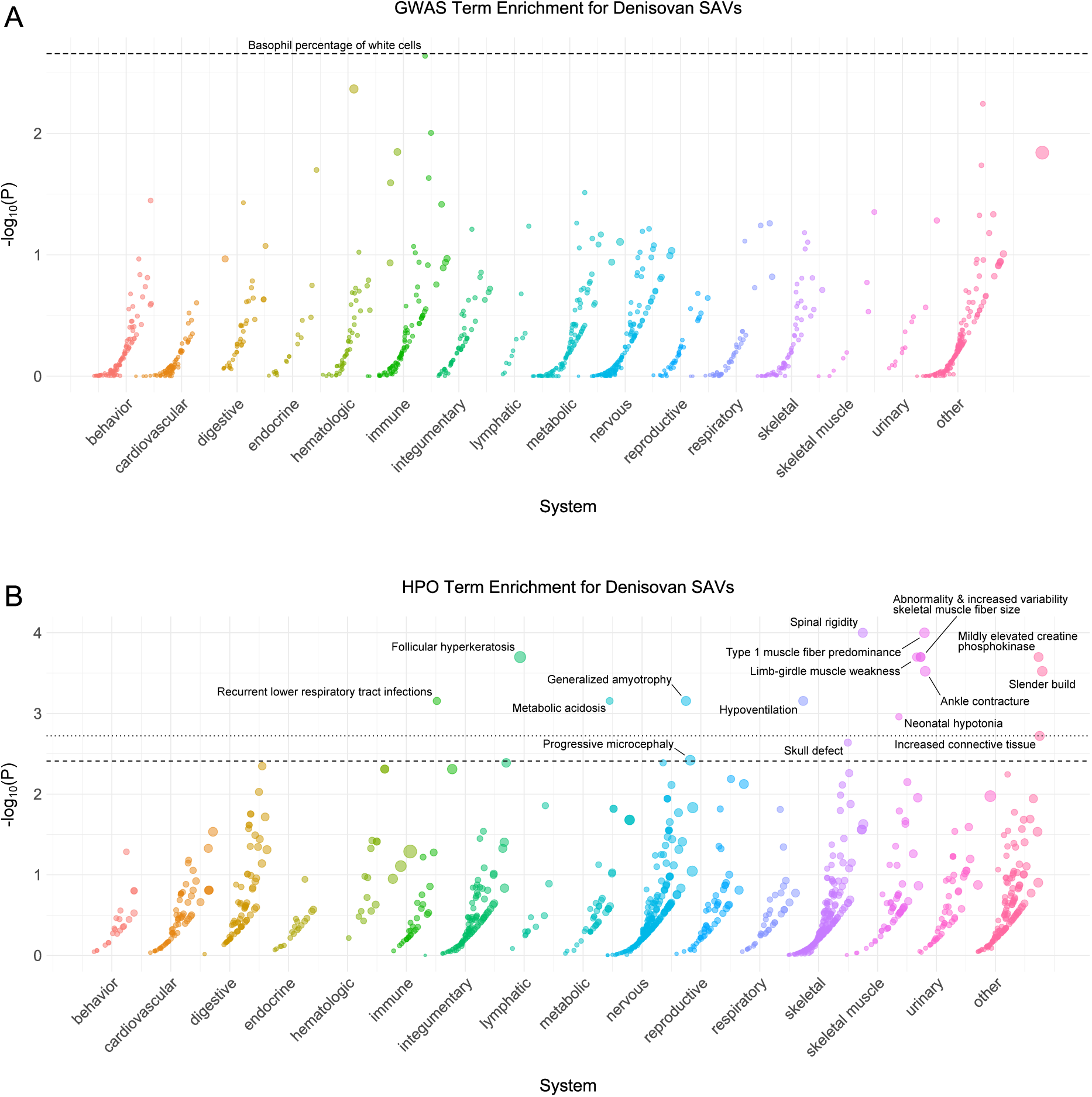
Denisovan phenotype enrichment. **(A)** Phenotype associations enriched among genes with archaic-specific Denisovan SAVs based on annotations from the 2019 GWAS Catalog. Phenotypes are ordered by increasing enrichment within manually curated systems. Circle size indicates enrichment magnitude. Enrichment and p-values were calculated from an empirical null distribution generated from 10,000 shuffles of maximum Δ across the entire dataset (Methods). Dotted and dashed lines represent false discovery rate (FDR) corrected p-value thresholds at FDR = 0.05 and 0.1, respectively. At least one example phenotype with a p-value *≤* the stricter FDR threshold (0.05) is annotated per system. **(B)** Phenotypes enriched among genes with archaic-specific Denisovan SAVs based on annotations from the Human Phenotype Ontology (HPO). Data were generated and visualized as in **A**. See **Supplementary Data 2** for all phenotype enrichment results.

**Extended Data Fig. 5.**
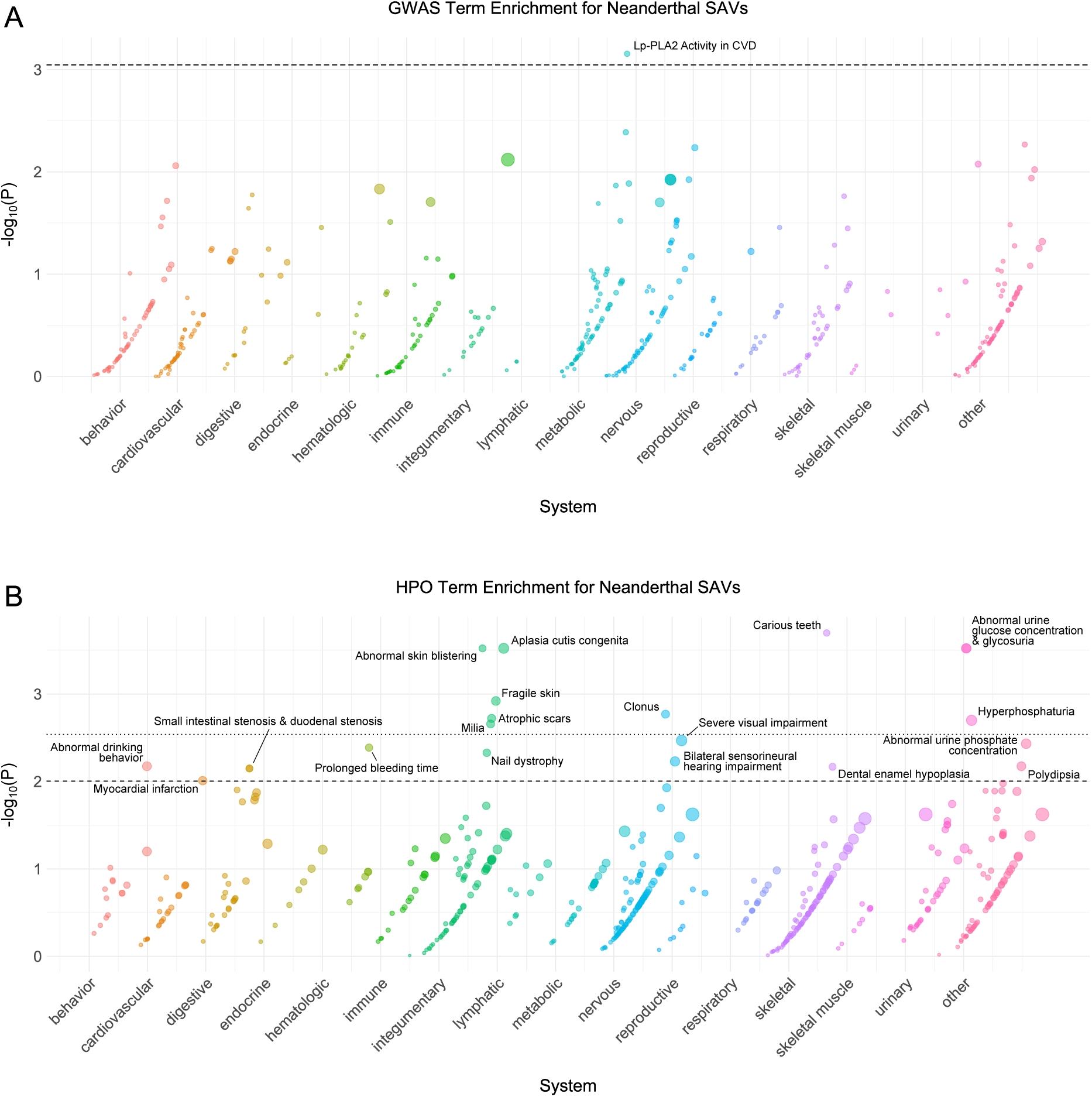
Neanderthal phenotype enrichment. **(A)** Phenotype associations enriched among genes with archaic-specific Neanderthal SAVs based on annotations from the 2019 GWAS Catalog. Phenotypes are ordered by increasing enrichment within manually curated systems. Circle size indicates enrichment magnitude. Enrichment and p-values were calculated from an empirical null distribution generated from 10,000 shuffles of maximum Δ across the entire dataset (Methods). Dotted and dashed lines represent false discovery rate (FDR) corrected p-value thresholds at FDR = 0.05 and 0.1, respectively. At least one example phenotype with a p-value *≤* the stricter FDR threshold (0.05) is annotated per system. Lp-PLA2 = Lipoprotein phospholipase A2. **(B)** Phenotypes enriched among genes with archaic-specific Neanderthal SAVs based on annotations from the Human Phenotype Ontology (HPO). Data were generated and visualized as in **A**. See **Supplementary Data 2** for all phenotype enrichment results.

**Extended Data Fig. 6.**
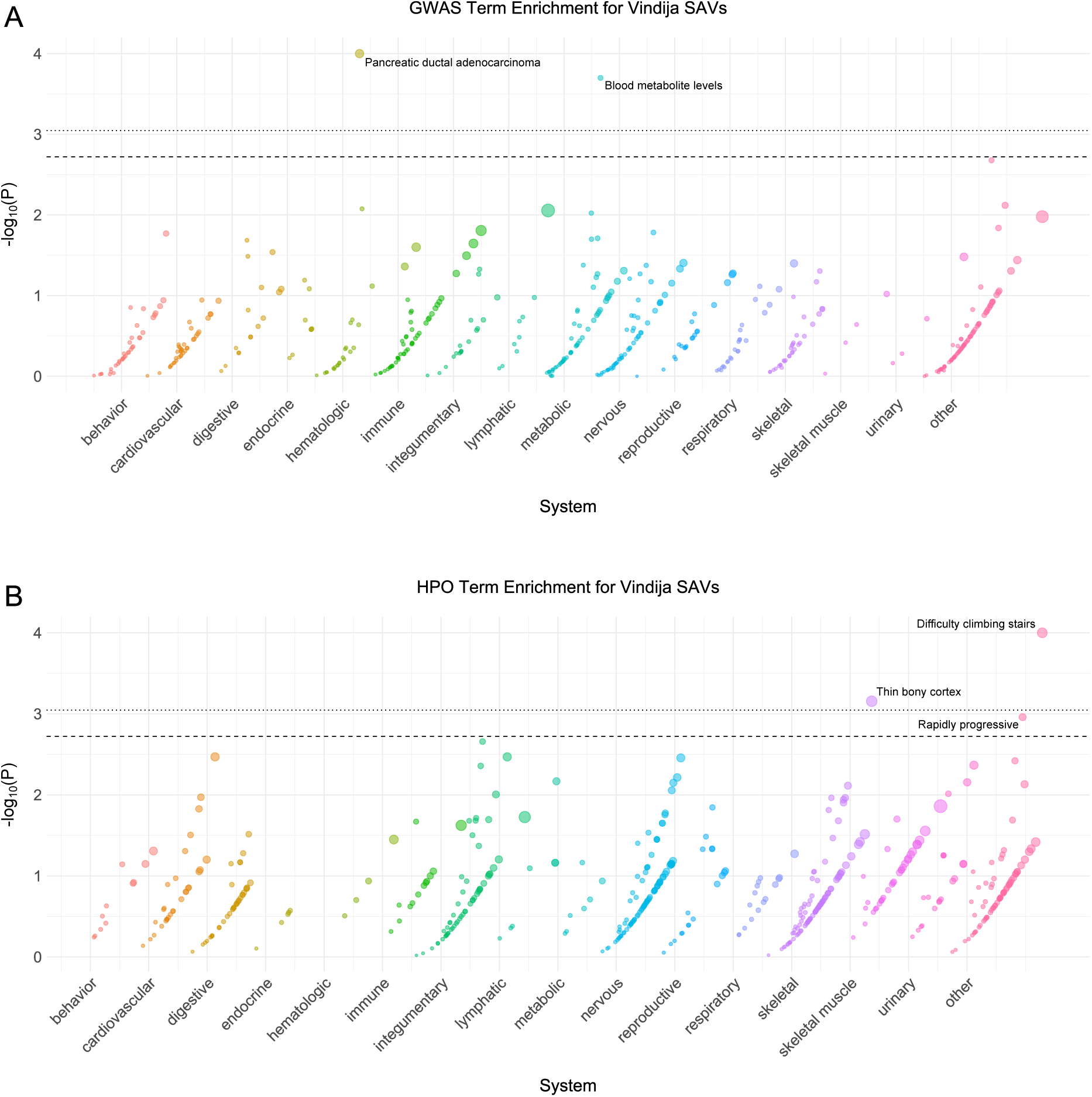
Vindija phenotype enrichment. **(A)** Phenotype associations enriched among genes with archaic-specific Vindija SAVs based on annotations from the 2019 GWAS Catalog. Phenotypes are ordered by increasing enrichment within manually curated systems. Circle size indicates enrichment magnitude. Enrichment and p-values were calculated from an empirical null distribution generated from 10,000 shuffles of maximum Δ across the entire dataset (Methods). Dotted and dashed lines represent false discovery rate (FDR) corrected p-value thresholds at FDR = 0.05 and 0.1, respectively. At least one example phenotype with a p-value *≤* the stricter FDR threshold (0.05) is annotated per system. **(B)** Phenotypes enriched among genes with archaic-specific Vindija SAVs based on annotations from the Human Phenotype Ontology (HPO). Data were generated and visualized as in **A**. See **Supplementary Data 2** for all phenotype enrichment results.

**Extended Data Fig. 7.**
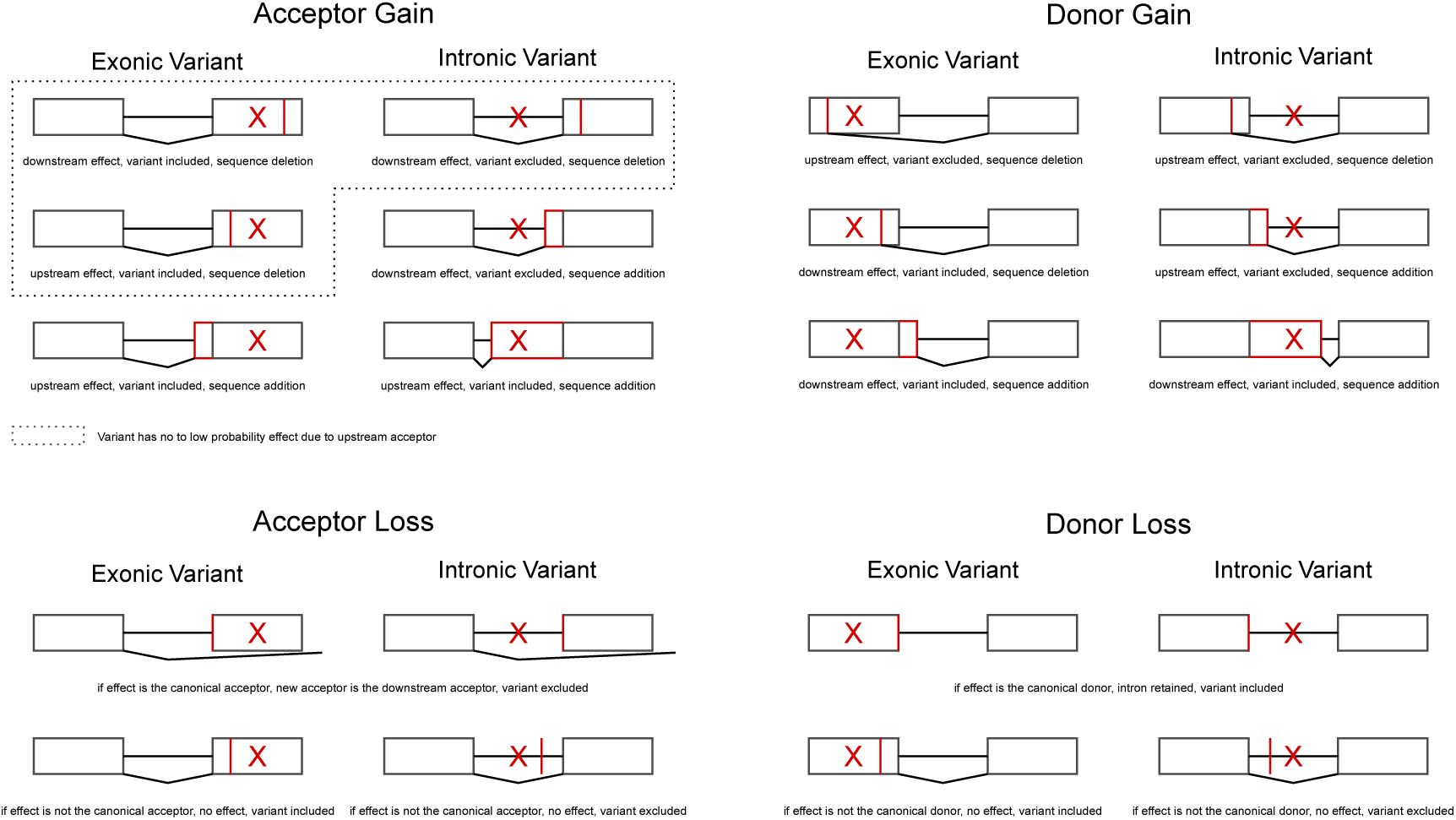
Modeling SAV effects on the canonical transcript. We used the SpliceAI output to construct a novel transcript per SAV by modifying the canonical transcript for that gene. We considered only one effect per SAV (e.g., either an acceptor gain, acceptor loss, donor gain, or donor loss) based on the effect with the largest Δ. Therefore, we did not model multiple effects for a single SAV (e.g., an acceptor gain and acceptor loss). Here, we illustrate all the possible consequences of a SAV for each of the four classes. We indicate the variant position with a red “X” and the position of the effect with a red vertical line (sequence deletion) or box (sequence addition). Each scenario includes a two exon gene (boxes) with a single intron (horizontal line).

**Extended Data Fig. 8.**
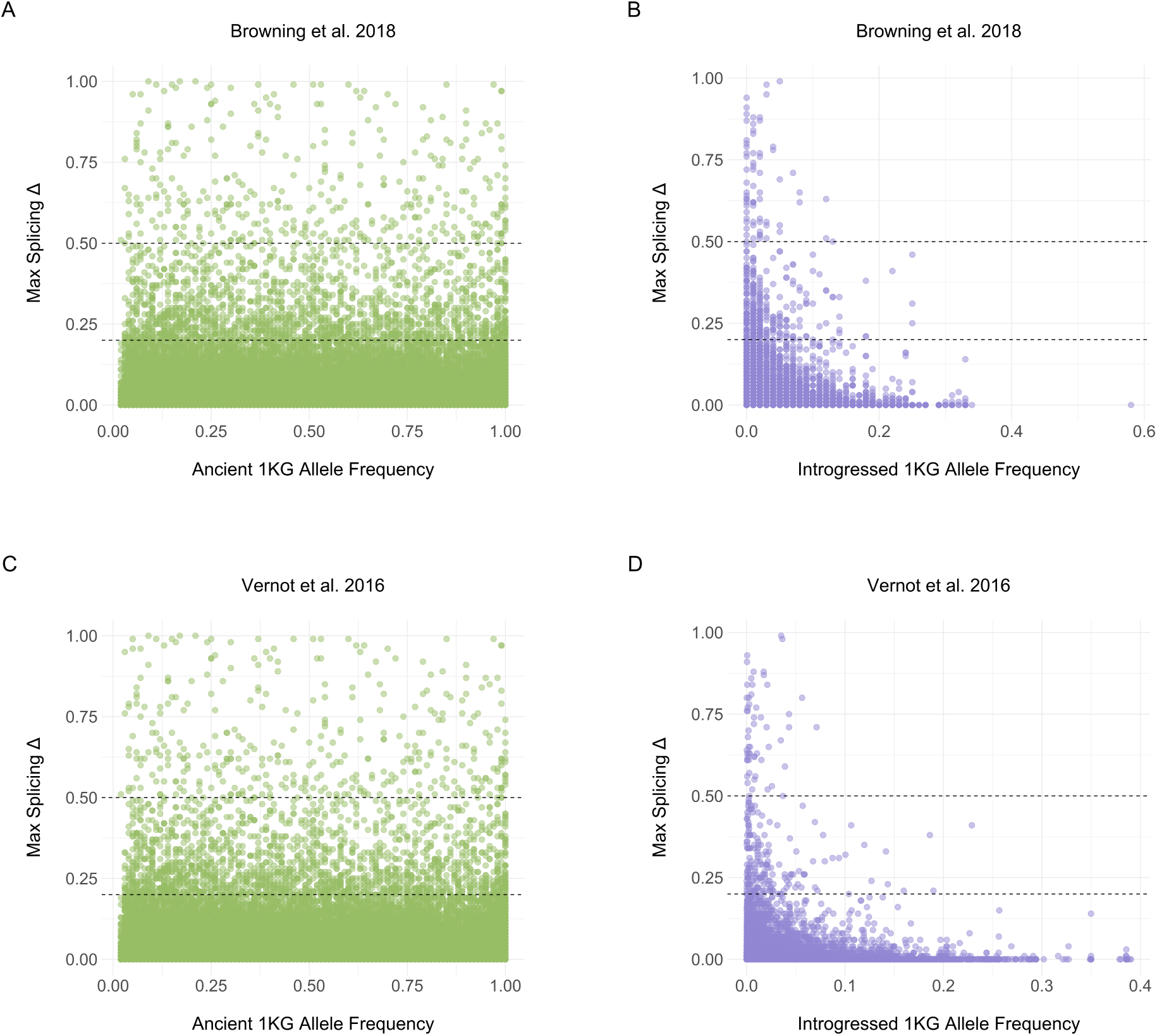
Δ max exhibits a variable relationship to 1KG allele frequency. **(A)** 1KG allele frequency and Δ max for all ancient variants per [39]. Allele frequencies are from 1KG. Dashed lines reflect both Δ thresholds. **(B)** 1KG allele frequency and Δ max for all introgressed variants per [39]. Allele frequencies are from 1KG. If the introgressed allele was the reference allele, we subtracted the 1KG allele frequency from 1. **(C)** 1KG allele frequency and Δ max for all ancient variants per [38]. Allele frequencies are from 1KG. **(D)** 1KG allele frequency and Δ max for all introgressed variants per [38]. Allele frequencies represent the mean from the AFR, AMR, EAS, EUR, SAS frequencies from the [38] metadata.

**Extended Data Fig. 9.**
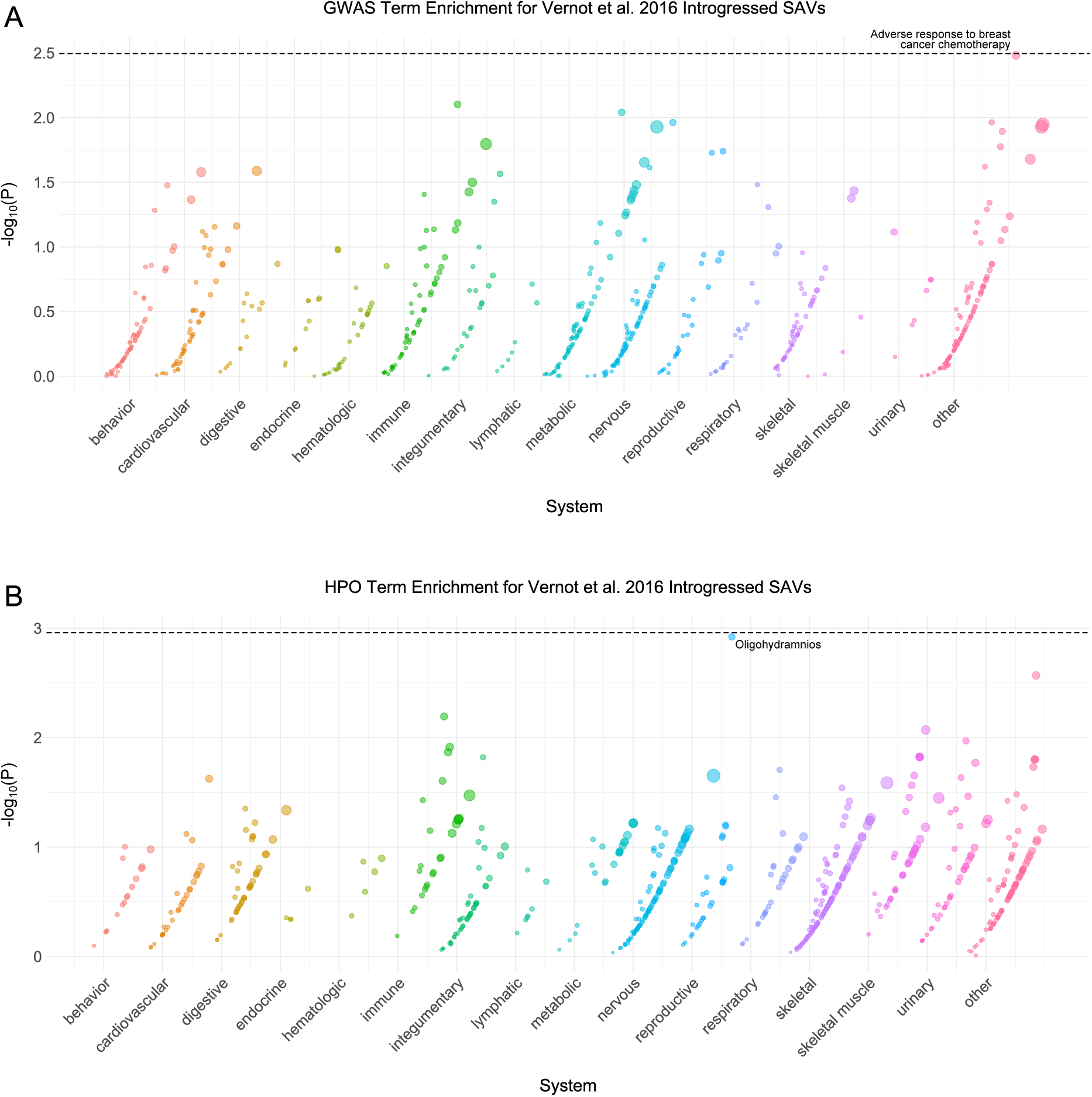
Vernot et al. 2016 introgressed phenotype enrichment. **(A)** Phenotype associations enriched among genes with [38] introgressed SAVs based on annotations from the 2019 GWAS Catalog. Phenotypes are ordered by increasing enrichment within manually curated systems. Circle size indicates enrichment magnitude. Enrichment and p-values were calculated from an empirical null distribution generated from 10,000 shuffles of maximum Δ across the entire dataset (Methods). Dotted and dashed lines represent false discovery rate (FDR) corrected p-value thresholds at FDR = 0.05 and 0.1, respectively. At least one example phenotype with a p-value *≤* the stricter FDR threshold (0.05) is annotated per system. **(B)** Phenotypes enriched among genes with [38] introgressed SAVs based on annotations from the Human Phenotype Ontology (HPO). Data were generated and visualized as in **A**. See **Supplementary Data 2** for all phenotype enrichment results.

**Extended Data Fig. 10.**
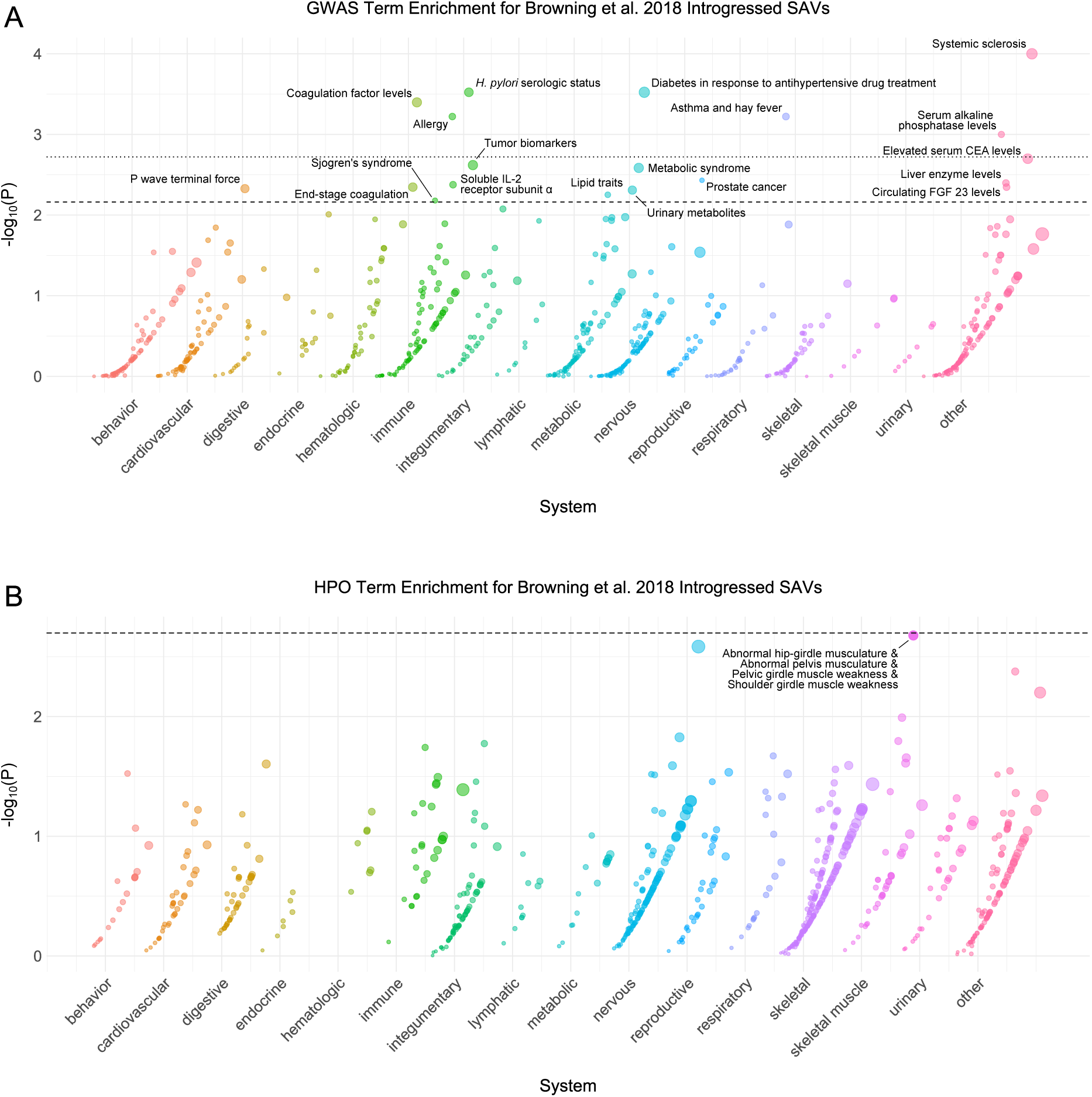
Browning et al. 2018 introgressed phenotype enrichment. **(A)** Phenotype associations enriched among genes with [39] introgressed SAVs based on annotations from the 2019 GWAS Catalog. Phenotypes are ordered by increasing enrichment within manually curated systems. Circle size indicates enrichment magnitude. Enrichment and p-values were calculated from an empirical null distribution generated from 10,000 shuffles of maximum Δ across the entire dataset (Methods). Dotted and dashed lines represent false discovery rate (FDR) corrected p-value thresholds at FDR = 0.05 and 0.1, respectively. At least one example phenotype with a p-value *≤* the stricter FDR threshold (0.05) is annotated per system. CEA = carcinoembryonic antigen, FGF = fibroblast growth factor. **(B)** Phenotypes enriched among genes with [39] introgressed SAVs based on annotations from the Human Phenotype Ontology (HPO). Data were generated and visualized as in **A**. See **Supplementary Data 2** for all phenotype enrichment results.

## 5 Supplementary Information

### 5.1 Supplementary Text

#### 5.1.1 The major spliceosome complex is strongly conserved between archaic hominins and modern humans

Alternative splicing occurs across nearly all eukaryotes and its molecular mechanisms are deeply conserved [41]. Thus, we anticipated that the sequence patterns learned by SpliceAI in humans would generalize to archaic hominins. To test for any significant differences, we first compared the sequences of 147 genes associated with the major spliceosome complex [42] between archaic hominins and modern humans. Analyzing all archaic variants absent in the 1000 Genomes Project (1KG) [43], we identified 19 non-synonymous archaic-specific variants, only 5 possibly or probably damaging by PolyPhen 2 and 8 of which were predicted to be deleterious by SIFT; [45], (**Supplementary Data 1**). Additionally, only two variants were predicted to disrupt protein function by both PolyPhen and SIFT (**Supplementary Data 1**). Furthermore, both deleterious variants were unique to the Altai Neanderthal.

We also considered the extent to which modern humans harbor deleterious variants in spliceosome associated genes. We subset variants that fell within spliceosome genes in each of the randomly sampled 1KG individuals and repeated the above procedure. We also repeated this for the archaics and included all spliceosome variants, regardless of whether or not they were archaic-specific. Modern humans exhibited between 0 and 3 deleterious variants, whereas the Chagyrskaya Neanderthal, Denisovan, and Vindija Neanderthal had 0 and the Altai Neanderthal had the two aforementioned variants (**Supplementary Table 14**).

Therefore, the major spliceosome complex is nearly identical between archaic hominins and modern humans, and there is no evidence that the sequence determinants of splicing have diverged.

#### 5.1.2 Physical characteristics of a gene are associated with the number of SAVs

Alternative splicing requires genes have at least one intron and the extent of splice altering variants may further be related to gene length. We considered the relationship between the the number of splicing variants (N = 0–11) in each gene and three gene traits: 1) number of exons, 2) length of gene body, and 3) length of coding sequence. These characteristics were positively associated at both Δ thresholds (**Supplementary Fig. 6**) (**Supplementary Table 6**). We also considered the relationship between the number of known isoforms per gene and the number of SAVs. We found a significant, positive association at both thresholds (**Supplementary Table 6**). However, this relationship is likely driven by the number of exons. When we conducted a partial correlation, controlling for the number of exons, both associations were non-significant with minimal effect size (Δ *≥* 0.2: partial Spearman, *ρ* = -0.003, P = 0.702, Δ *≥* 0.5: partial Spearman, *ρ* = 0.0008, P = 0.9174).

#### 5.1.3 Sprime identifies more introgressed SAVs present in all five 1KG superpopulations

Comparing our results using two different sets of introgressed variants [38, 39] revealed very similar patterns. For example, most ancient SAVs occurred at *≥* 0.05 frequency in all five 1KG superpopulations (**Supplementary Fig. 12**). The next most frequent were SAVs that were present in all but east Asians. SAVs shared among non-Africans and those present in all but Europeans were also common.

However, there were a few differences between the datasets in the distribution of introgressed SAVs across populations. We found that among those introgressed SAVs classified per [38], many were shared among non-Africans (**Fig. 6D**). However, introgressed SAVs classified per [39] were most commonly present in both Africans and non-Africans, followed by a smaller set of non-African SAVs (**Supplementary Fig. 14**). This difference likely reflects low frequency introgressed variants that are allowed to occur in the reference population in Sprime. While introgression between modern humans and archaics occurred outside of Africa, many Africans have low levels of Neanderthal ancestry due to backflow from Eurasians into Africans [94]. Among non-African subsets, both datasets were largely similar in their distribution (**Supplementary Figs. 14**, **15**).

#### 5.1.4 Ancient and introgressed sQTL SAVs have similar tissue activity patterns in GTEx

To test for differences in the tissues of activity for SAVs of different origin, we compared the distribution of ancient and introgressed SAVs identified in GTEx (**Supplementary Table 15**). The number of sQTLs in each tissue was positively correlated between ancient and introgressed variants, and this held for both sets of introgressed variants (Browning et al. 2018: Spearman, *ρ* = 0.94, P = 3.8 *×* 10*^−^*^24^; Vernot et al. 2016: Spearman, *ρ* = 0.92, P = 1.09 *×* 10*^−^*^20^) (**Supplementary Fig. 22**). However, the ability to detect sQTL is strongly influenced by sample size, and we found that the number of individuals sampled for each tissue is positively correlated with the number of sQTL SAVs in a tissue for both ancient variants (Browning et al. 2018: Spearman, *ρ* = 0.91, P = 5.73 *×* 10*^−^*^20^; Vernot et al. 2016: Spearman, *ρ* = 0.91, P = 8.43 *×* 10*^−^*^20^) and introgressed variants (Browning et al. 2018: Spearman, *ρ* = 0.89, P = 2.47 *×* 10*^−^*^17^; Vernot et al. 2016: Spearman, *ρ* = 0.85, P = 1.34 *×* 10*^−^*^14^) (**Supplementary Figs. 23**, **24**). We also note that the proportion of sQTL SAVs compared to all significant GTEx sQTLs detected per tissue was largely similar for most tissues (**Supplementary Fig. 25**). These results suggest that tissues of activity between SAVs of different allele origins are similar; however, these analyses are limited by the coverage and power of GTEx sQTL data.

We also considered tissue-level gene expression differences by comparing the number of genes expressed (TPM > 1) in each tissue with at least one SAV of a given origin. We found no significant differences among allele origins for either allele origin classification (**Supplementary Figs. 26**, **27**). As for the sQTL, these patterns were mainly driven by differences in sample size and number of expressed genes in each GTEx tissue.

### 5.2 Supplementary Data

**Supplementary Data 1.** Variant effect predictor (VEP) output from Ensembl for non-synonymous, archaic-specific variants in genes associated with the major spliceosome complex.

**Supplementary Data 2.** This file contains the outputs from the gene ontology analysis for GWAS Catalog 2019 and Human Phenotype Ontology terms.

**Supplementary Data 3.** This file contains information on the novel isoforms constructed for each SAV.

### 5.3 Supplementary Figures

**Supplementary Fig. 1.**
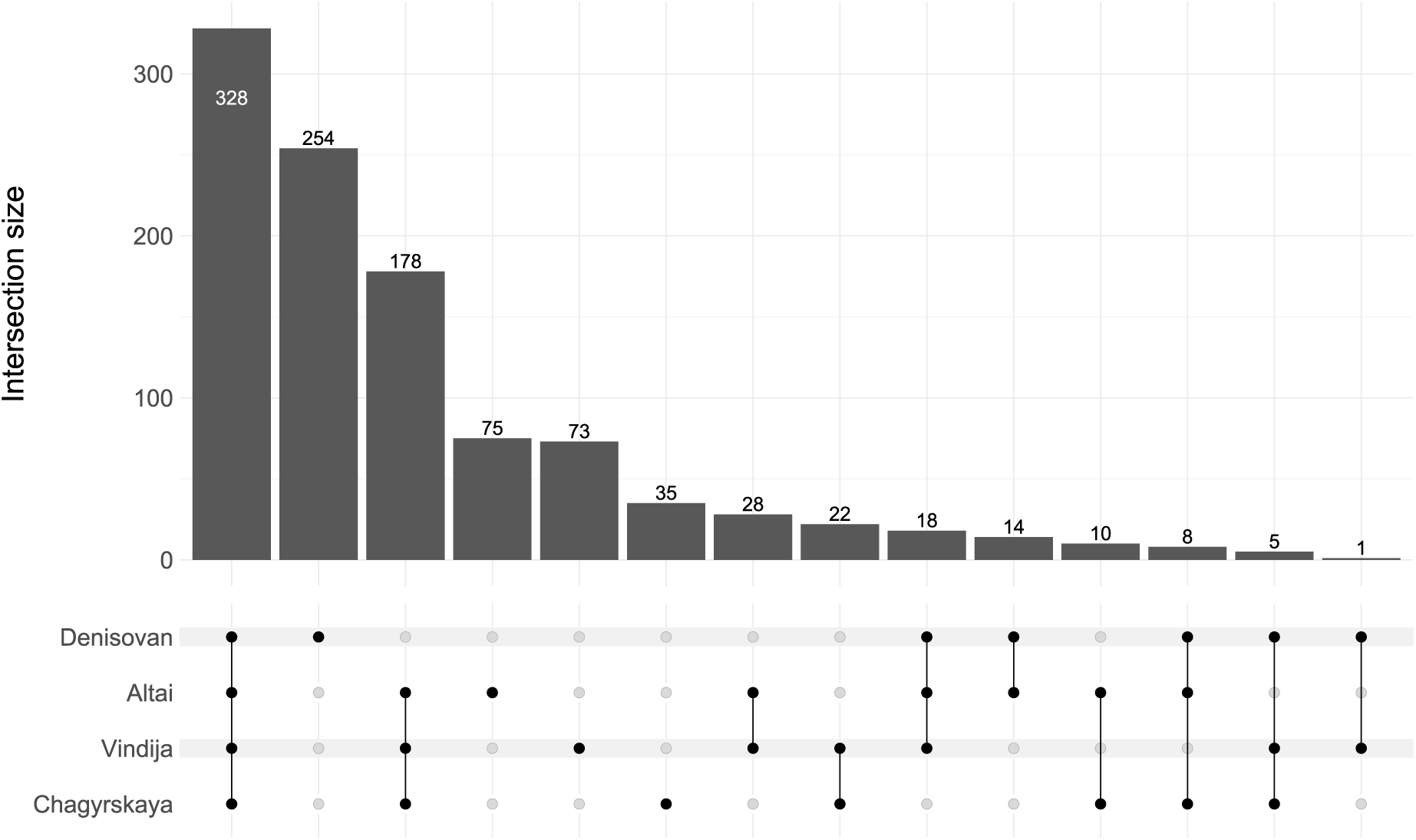
The distribution of SAVs is consistent with the archaic hominin phylogeny. Unique and shared SAVs at Δ *≥* 0.5.

**Supplementary Fig. 2.**
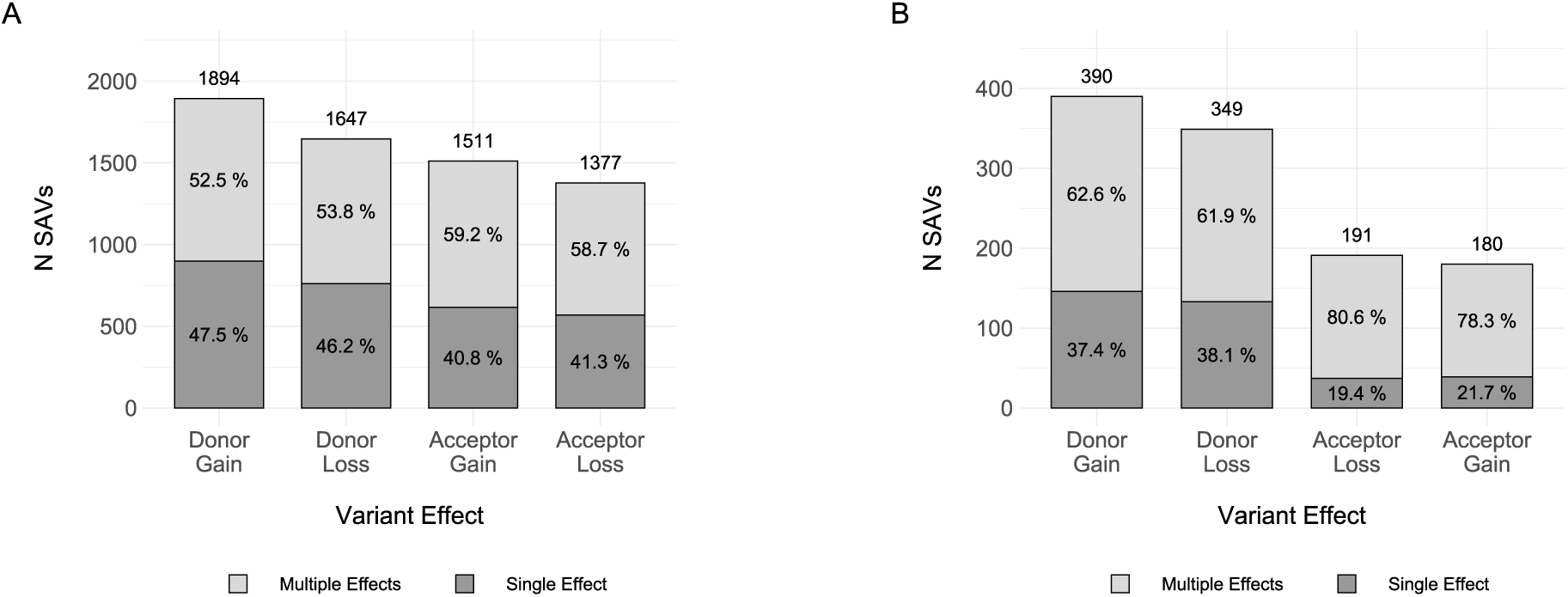
SAVs most commonly result in a donor gain. **(A)** The distribution of variant effects for all variants at the Δ *≥* 0.2 threshold. The N above each column indicates the number of variants with that effect above the threshold. Some variants result in multiple effects; therefore, the sum of these classes *̸*= 5,950. The color in each stacked bar indicates whether the effect is unique (dark grey) or one of many effects (light grey). **(B)** The distribution of variant effects for all variants at the Δ *≥* 0.5 threshold. As in **A**, some variants result in multiple effects; therefore, the sum of these classes *̸*= 1,049. The bar stacks are colored as in **A**.

**Supplementary Fig. 3.**
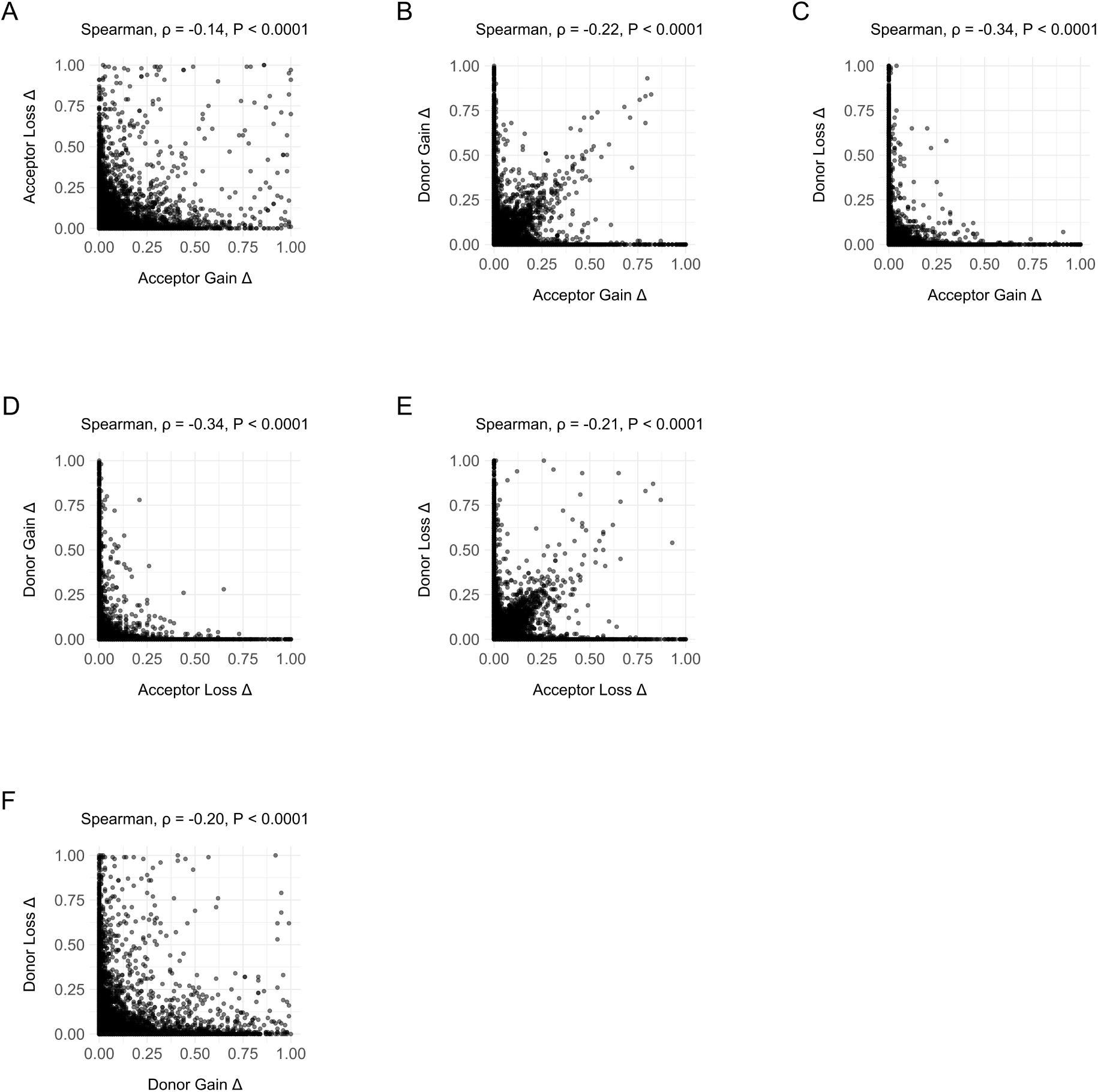
Delta scores between classes are negatively associated. **(A)** Association between acceptor gain Δs and acceptor loss Δs for all variants with at least one Δ > 0. **(B)** Association between acceptor gain Δs and donor gain Δs for all variants with at least one Δ > 0. **(C)** Association between acceptor gain Δs and donor loss Δs for all variants with at least one Δ > 0. **(D)** Association between acceptor loss Δs and donor gain Δs for all variants with at least one Δ > 0. **(E)** Association between acceptor loss Δs and donor loss Δs for all variants with at least one Δ > 0. **(F)** Association between donor gain Δs and donor loss Δs for all variants with at least one Δ > 0.

**Supplementary Fig. 4.**
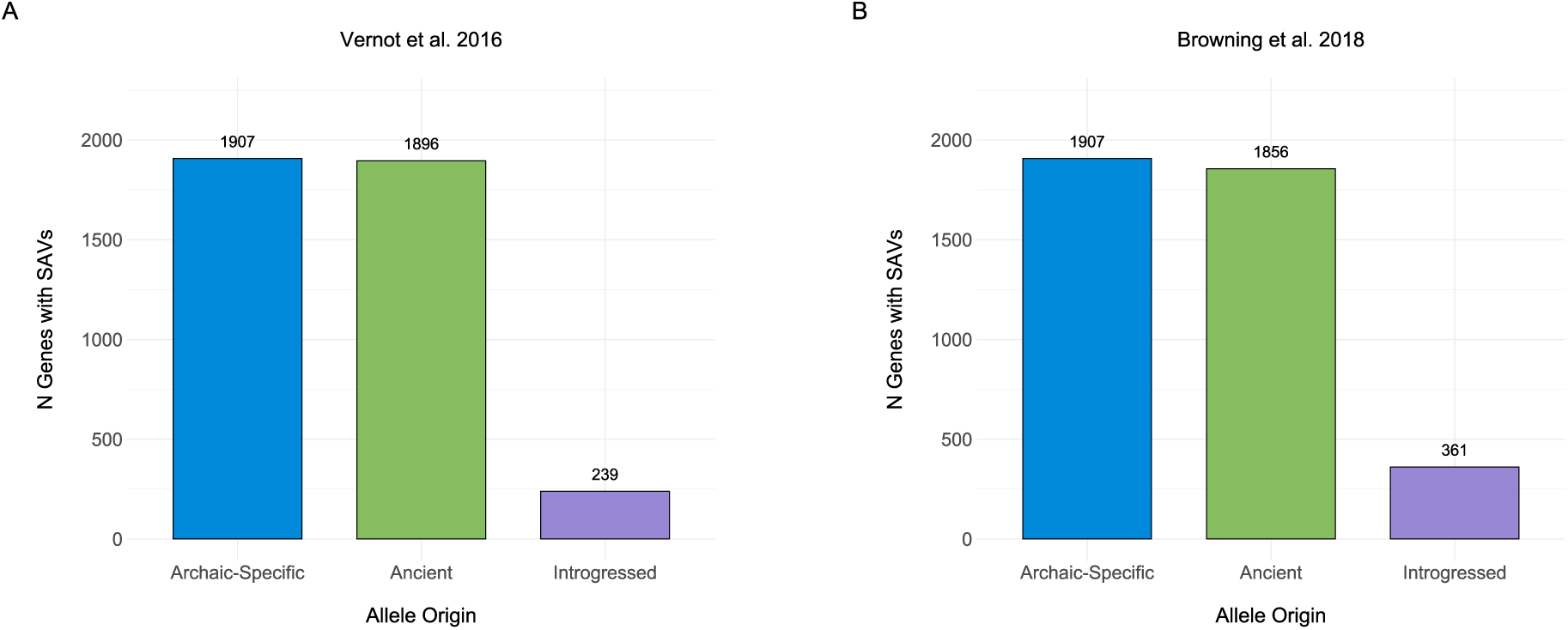
N genes with SAVs by allele origin. **(A)** The number of genes encompassed by each SAV allele origin per [39] at Δ *≥* 0.2. Some genes may occur in multiple categories. **(B)** The number of genes encompassed by each SAV allele origin per [38] at Δ *≥* 0.2. Some genes may occur in multiple categories.

**Supplementary Fig. 5.**
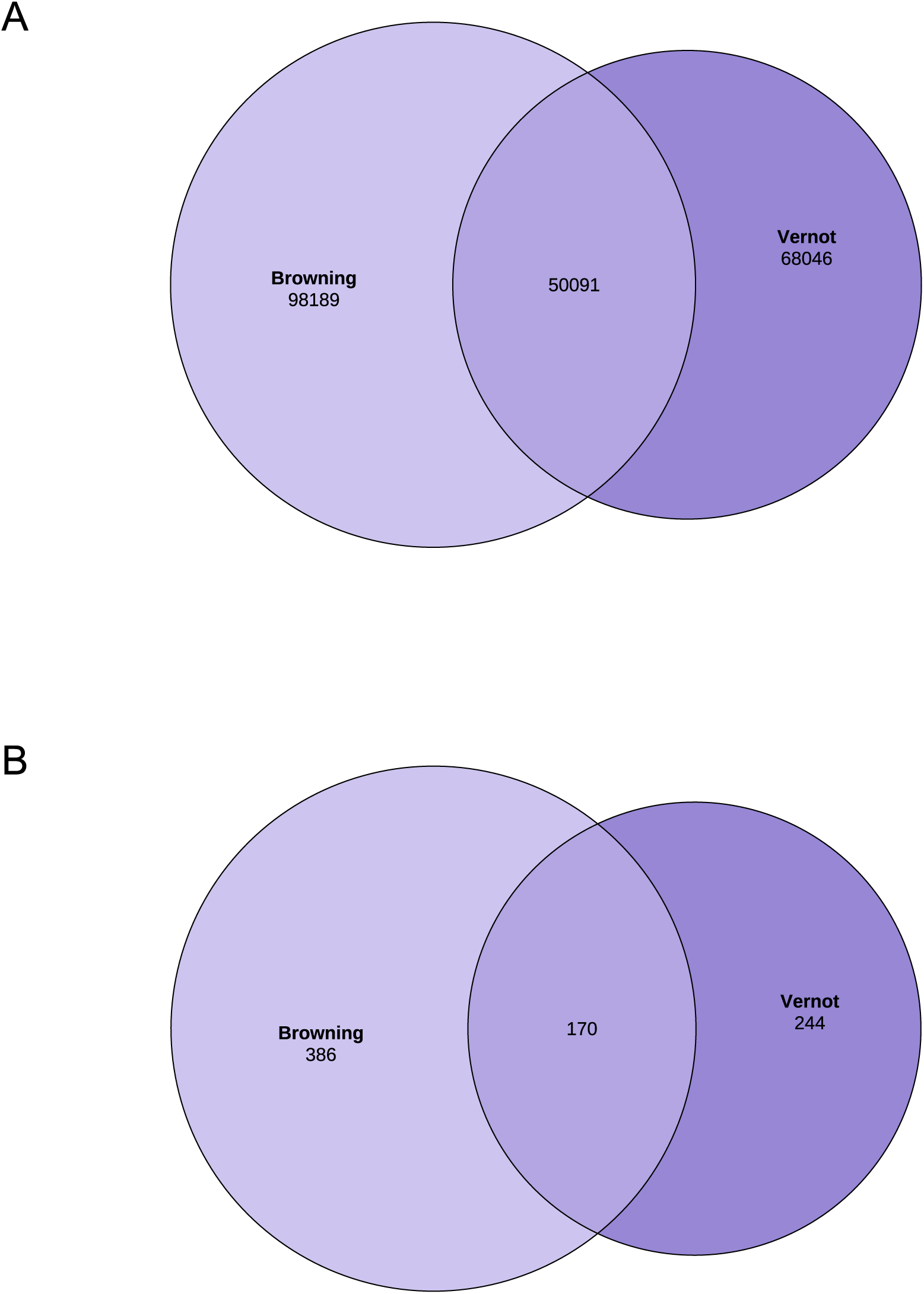
Two different sets of introgressed variants modestly overlap. **(A)** The overlap between introgressed variants from [39] and [38] that match a quality filtered locus in this study and are present in 1KG and/or gnomAD. **(B)** A subset of the overlap for variants with Δ *≥* 0.2.

**Supplementary Fig. 6.**
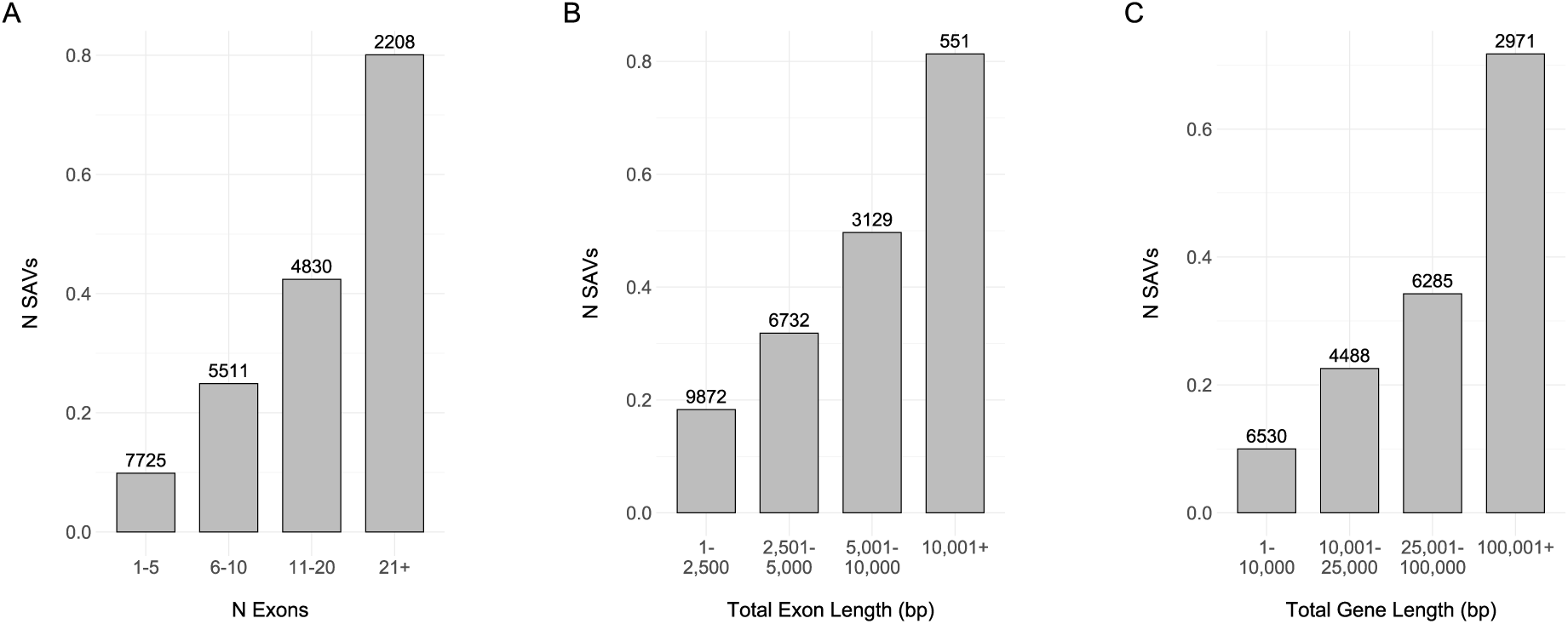
The number of exons and gene length are associated with more SAVs. **(A)** Binned number of exons and the mean number of SAVs at Δ *≥* 0.2 per bin. N reflects the number of genes per bin. **(B)** Binned exon length in bp and the mean number of SAVs at Δ *≥* 0.2 per bin. N reflects the number of genes per bin. **(C)** Binned gene length in bp and the mean number of SAVs at Δ *≥* 0.2 per bin. N reflects the number of genes per bin.

**Supplementary Fig. 7.**
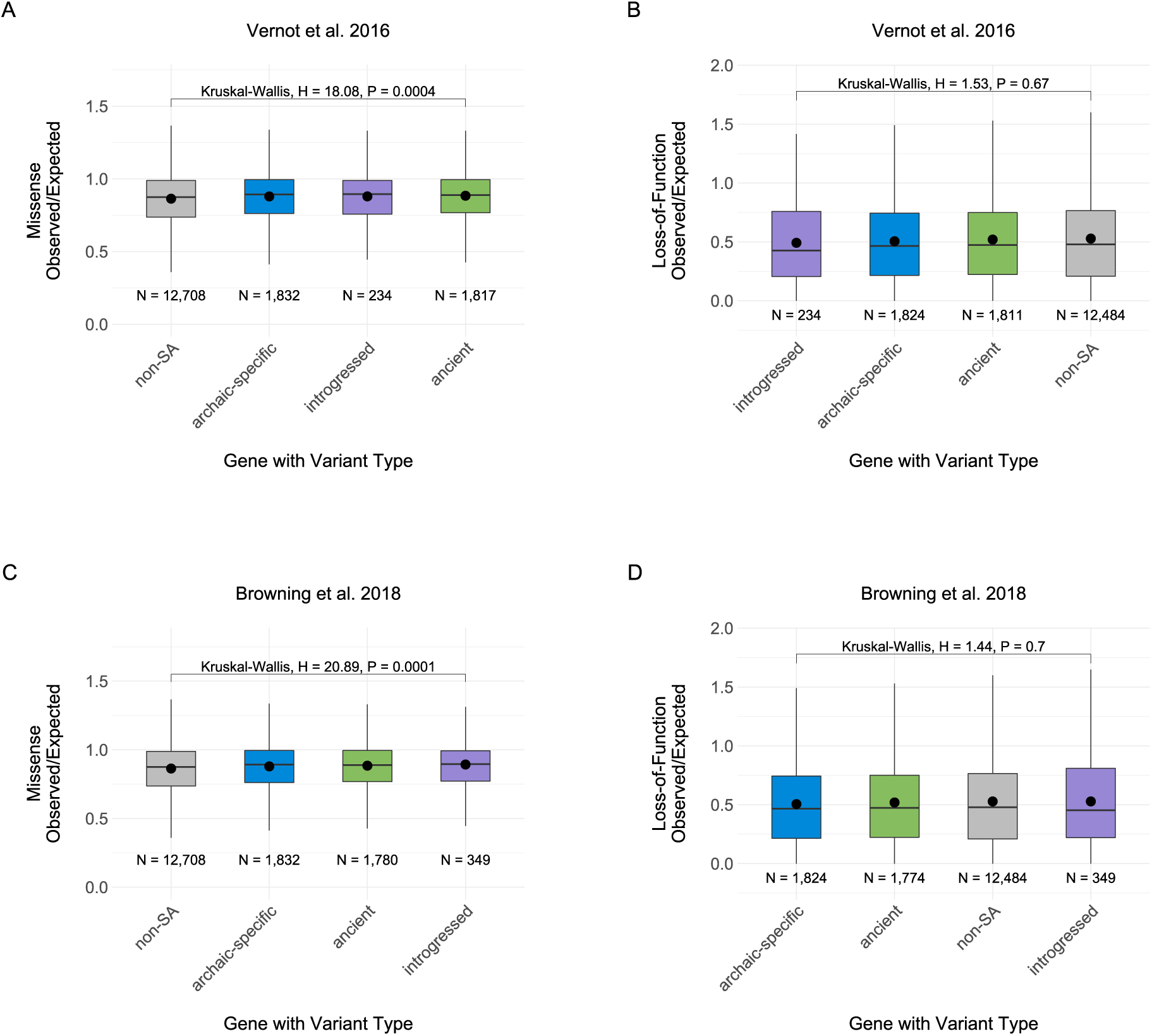
Gene mutational tolerance does not constrain the evolutionary history of SAVs. **(A)** Observed over expected ratio for missense variants per gene from gnomAD among four different sets. The ancient, archaic-specific, and introgressed sets from [39] include any gene that had *≥* 1 SAV at the Δ *≥* 0.2 threshold. Non-SA genes are all genes that do not occur in any of the other three sets. Boxplots indicate the first quantile, median, and third quantile and the mean is noted by the black point. Ns reflect the number of variants per set. **(B)** Observed over expected ratio for loss-of-function variants per gene from gnomAD among four different sets from [39]. The ancient, archaic-specific, and introgressed sets include any gene that had *≥* 1 SAV at the Δ *≥* 0.2 threshold. Non-SA genes are all genes that do not occur in any of the other three sets. Boxplots indicate the first quantile, median, and third quantile and the mean is noted by the black point. Ns reflect the number of variants per set. **(C)** Observed over expected ratio for missense variants per gene from gnomAD among four different sets from [38]. **(D)** Observed over expected ratio for loss-of-function variants per gene from gnomAD among four different sets from [38].

**Supplementary Fig. 8.**
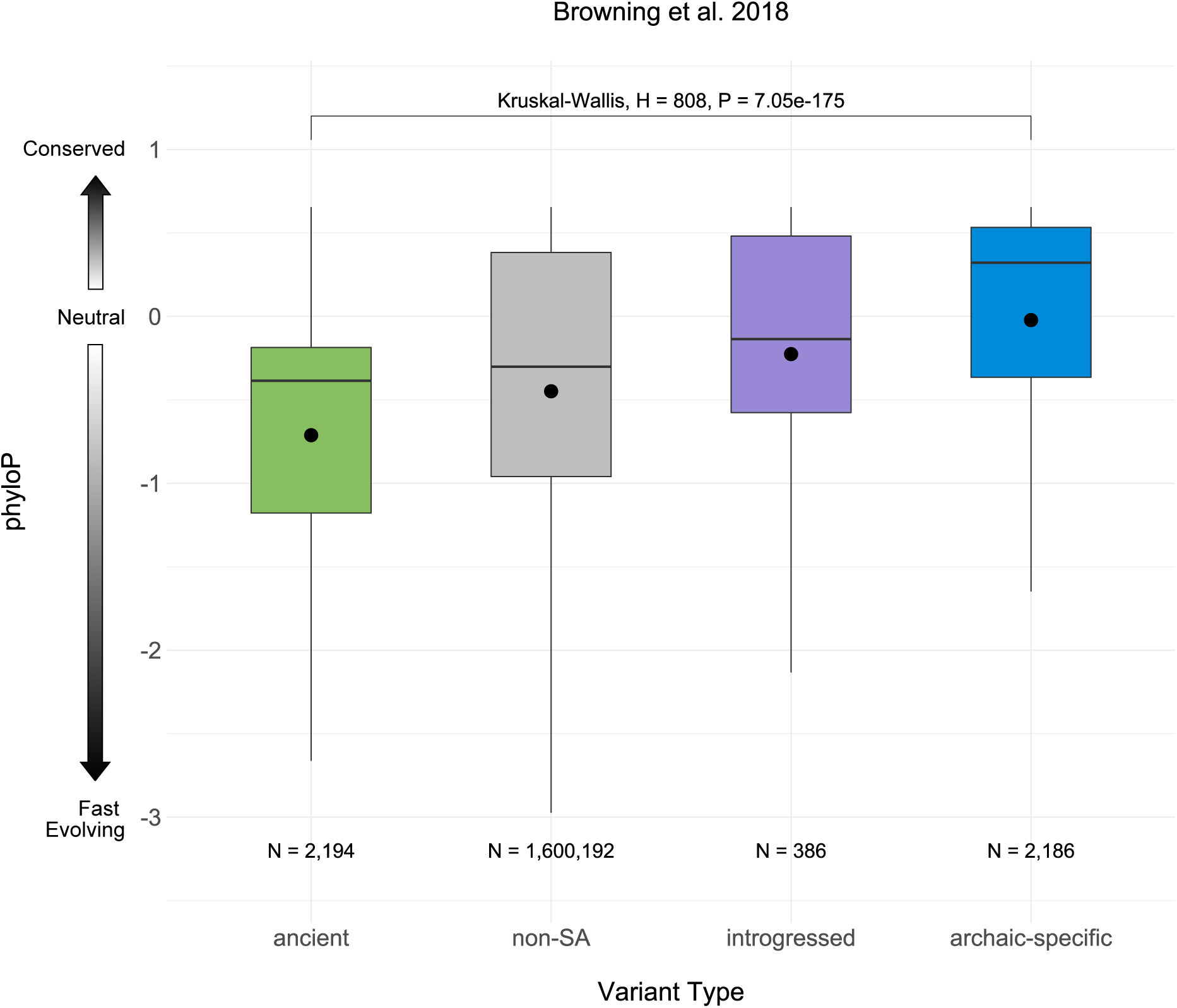
Site-level evolutionary conservation varies across SAVs with different origins. phyloP evolutionary conservation score distributions for archaic SAVs of different origins and non-SAVs. Positive scores indicate substitution rates slower than expected under neutral evolution (conservation), while negative scores indicate higher substitution rates than expected (fast evolution). The boxplots give the first quantile, median, and third quantile of the distributions, and the mean is noted by the black point. Ns are the number of variants per set. SAV classifications were based on [39] introgression calls.

**Supplementary Fig. 9.**
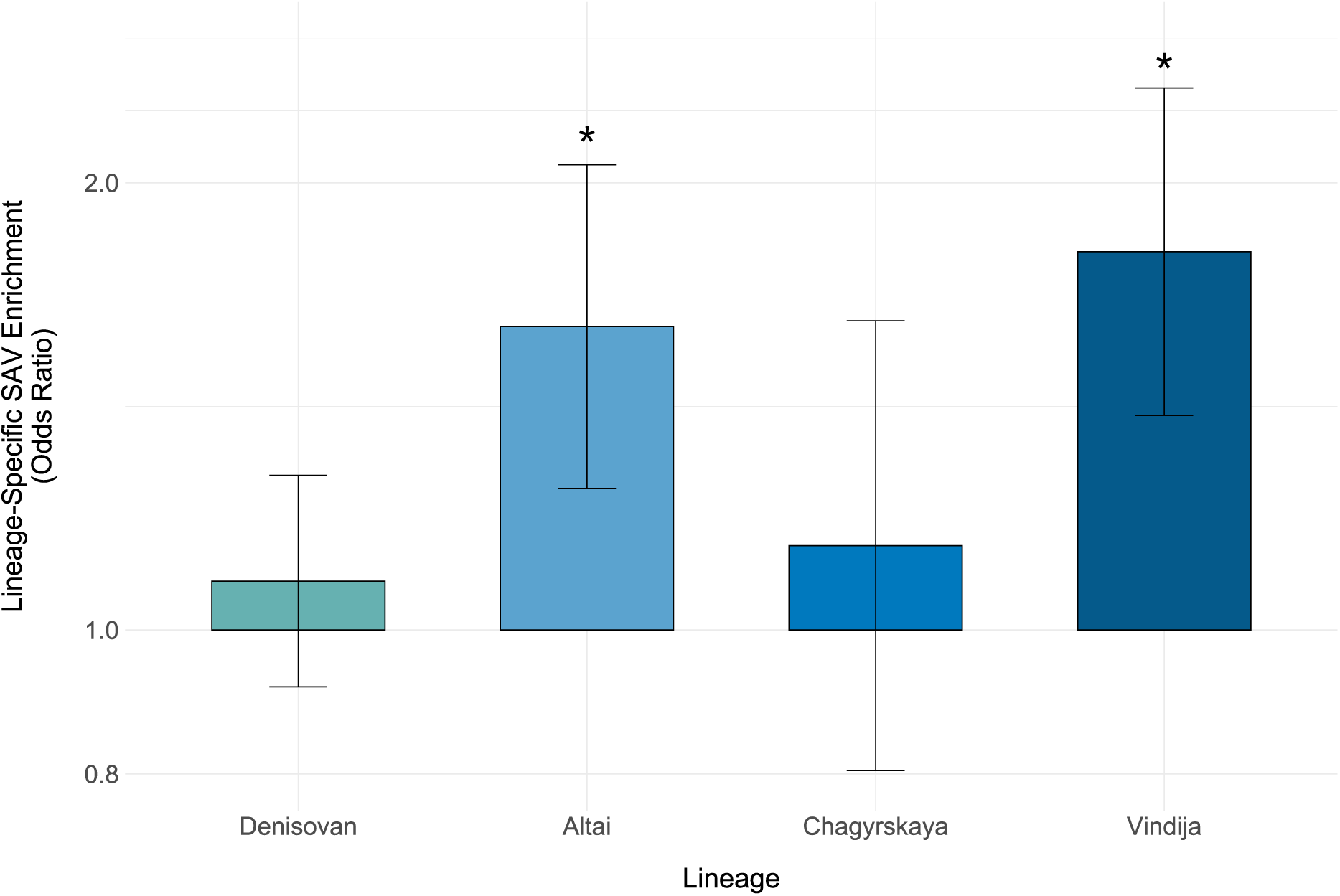
Lineage-specific high-confidence SAVs are enriched in two Neanderthals. Odds ratios from Fisher’s exact test performed for each lineage’s unique high confidence SAVs/non- SAVs compared to those shared among all four individuals. The number of variants used in each enrichment test are listed in **Supplementary Table 9**. Asterisks reflect significance using a Bonferroni corrected *α* (0.0125). Error bars denote the 95% CI. Note the y-axis is log10 transformed.

**Supplementary Fig. 10.**
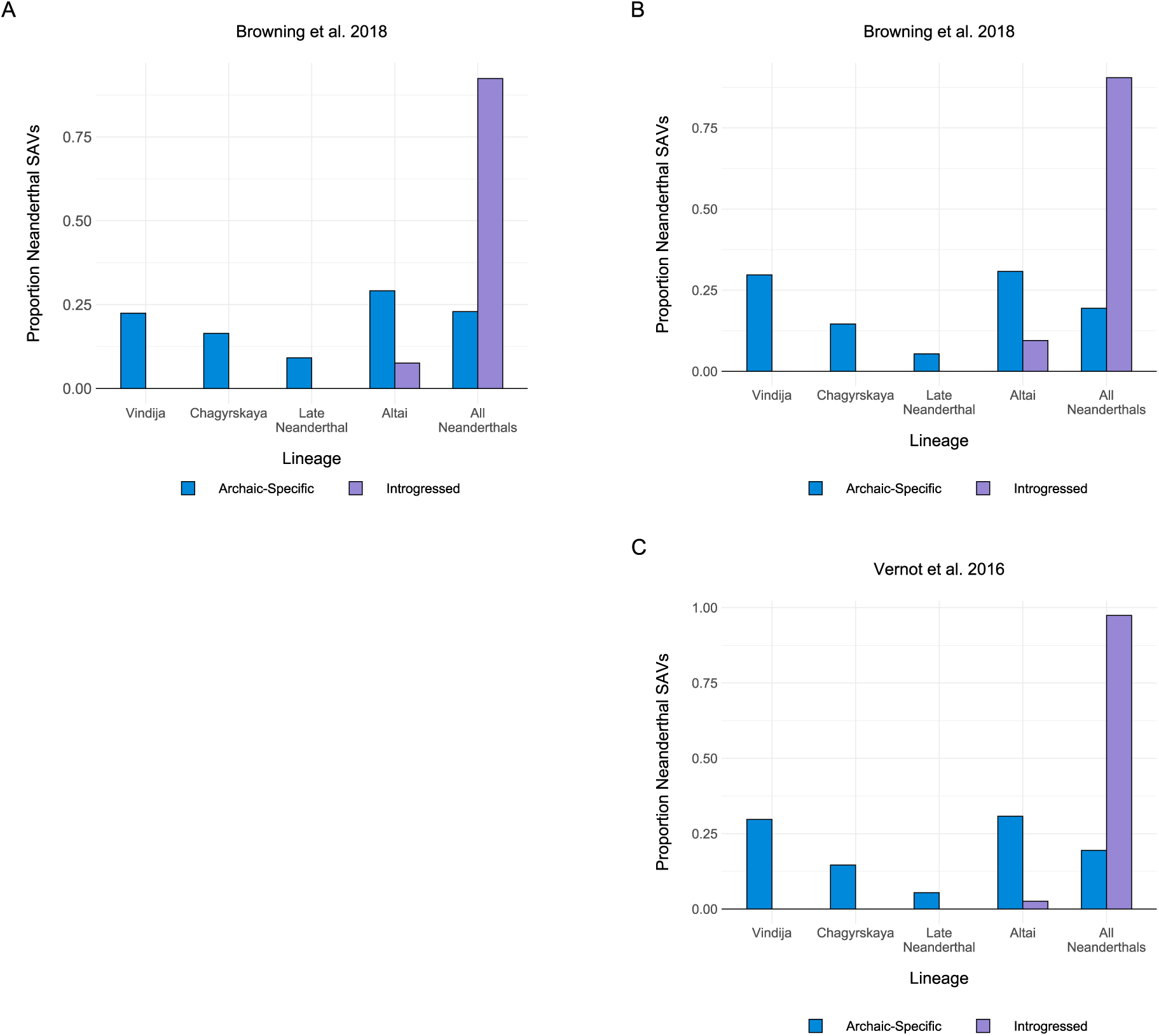
Introgressed SAVs are largely older variants. **(A)** Proportion of SAVs at Δ *≥* 0.2 per Neanderthal lineage among archaic-specific SAVs (expected) and introgressed SAVs (observed) from [39]. Proportions were calculated from the sum of all Neanderthal lineages because power to detect introgressed Denisovan SAVs is low. All data are presented in **Supplementary Table 10**. **(B)** Proportion of SAVs at Δ *≥* 0.5 using introgressed SAVs from [39]. **(C)** Proportion of SAVs at Δ *≥* 0.5 using introgressed SAVs from [38].

**Supplementary Fig. 11.**
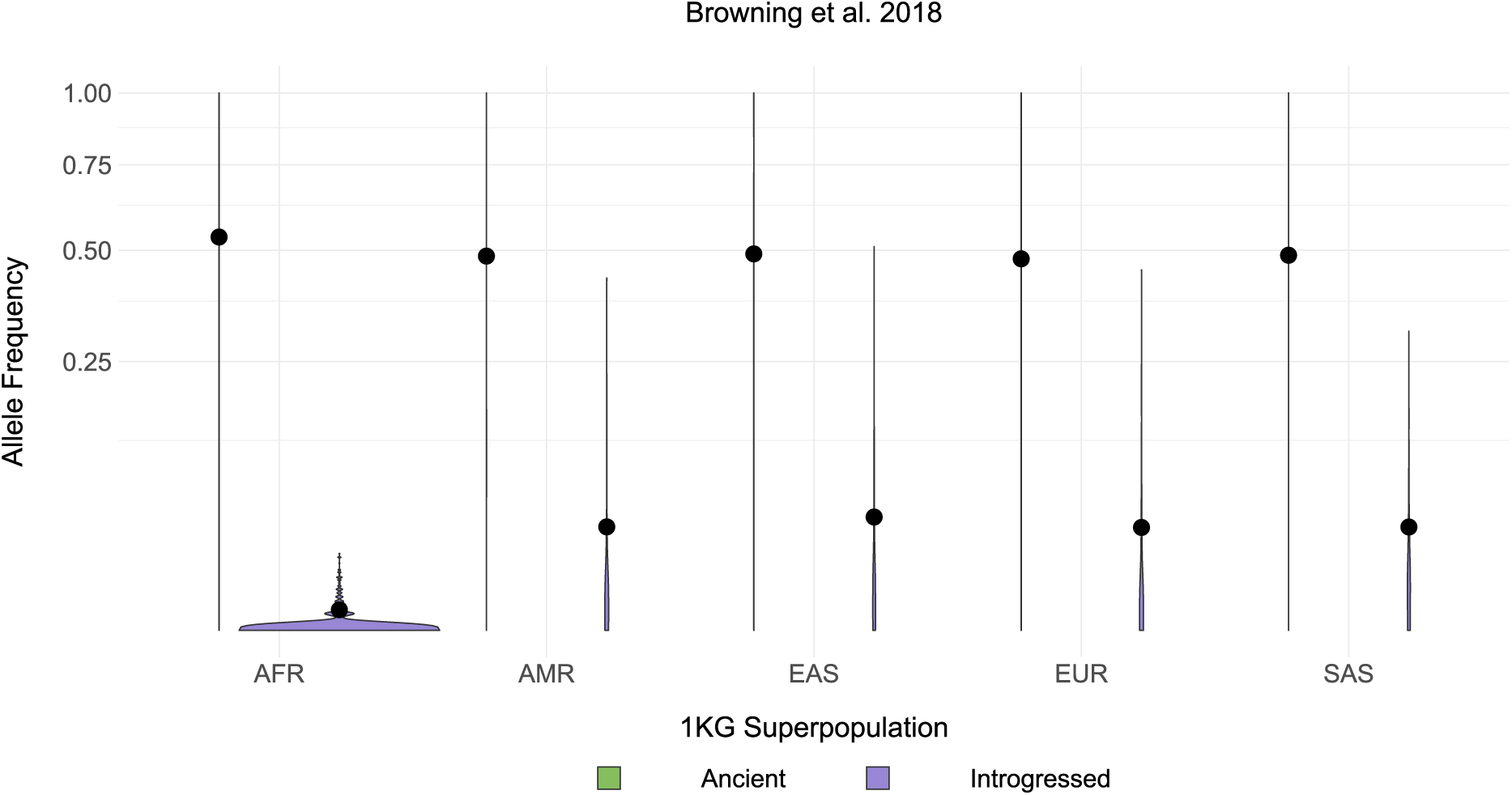
Ancient SAVs occur a moderate to high frequencies, whereas introgressed SAVs occur at lower frequencies. Allele frequency distributions for SAVs at Δ *≥* 0.2 per [39] by 1KG superpopulation. Allele frequencies are from 1KG. If the introgressed allele was the reference allele, we subtracted the 1KG allele frequency from 1. The black dot represents the mean allele frequency. Note the y-axis is square root transformed. AFR = African, AMR = American, EAS = East Asian, EUR = European, SAS = South Asian.

**Supplementary Fig. 12.**
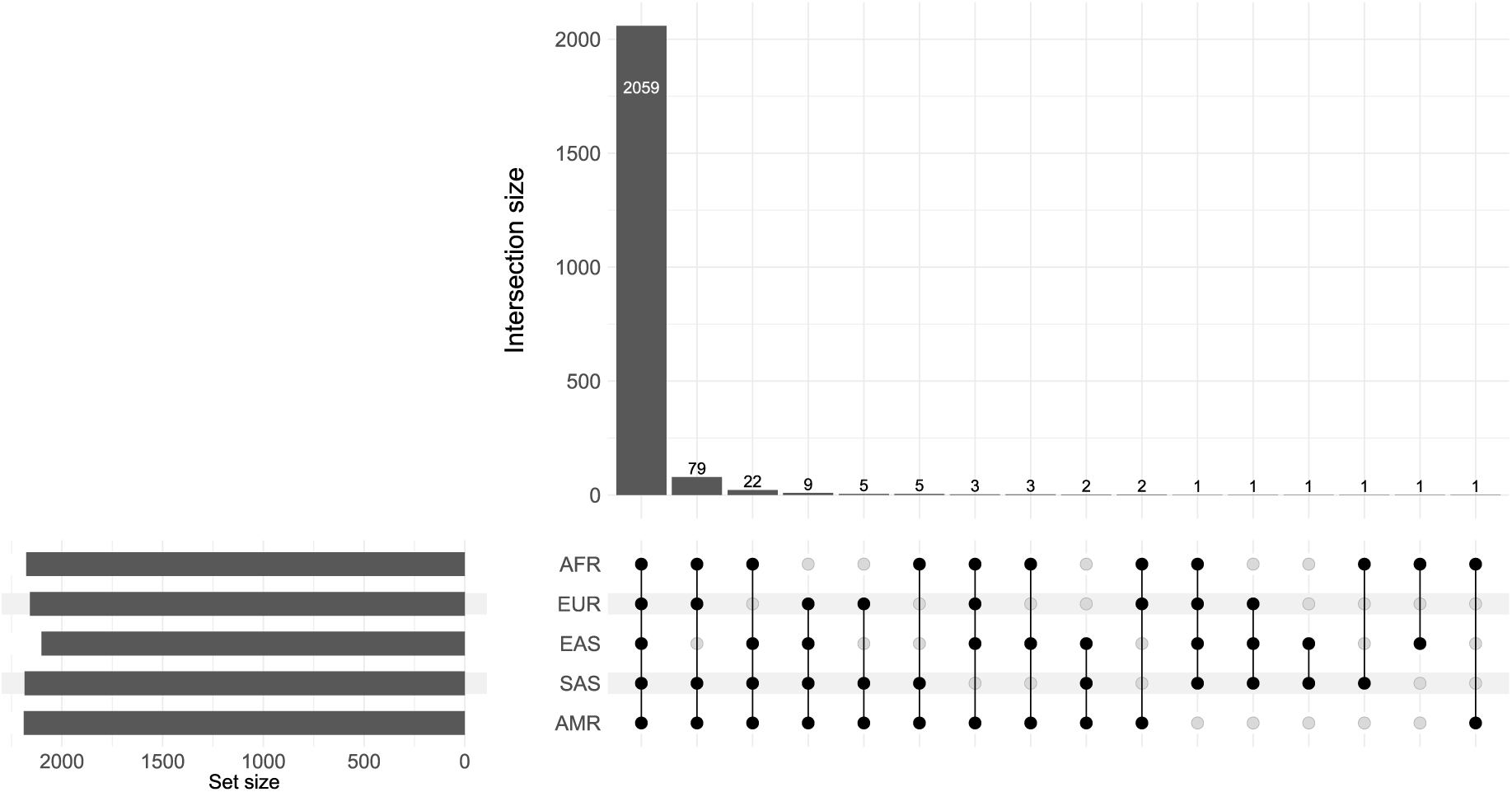
Most ancient SAVs occur in all 1KG superpopulations. Unique and shared ancient SAVs per [39] at Δ *≥* 0.2. Allele frequencies are from 1KG. By definition, each ancient SAV was considered present in a population if the allele frequency was *≥* 0.05 for at least two superpopulations. AFR = African, AMR = American, EAS = East Asian, EUR = European, SAS = South Asian.

**Supplementary Fig. 13.**
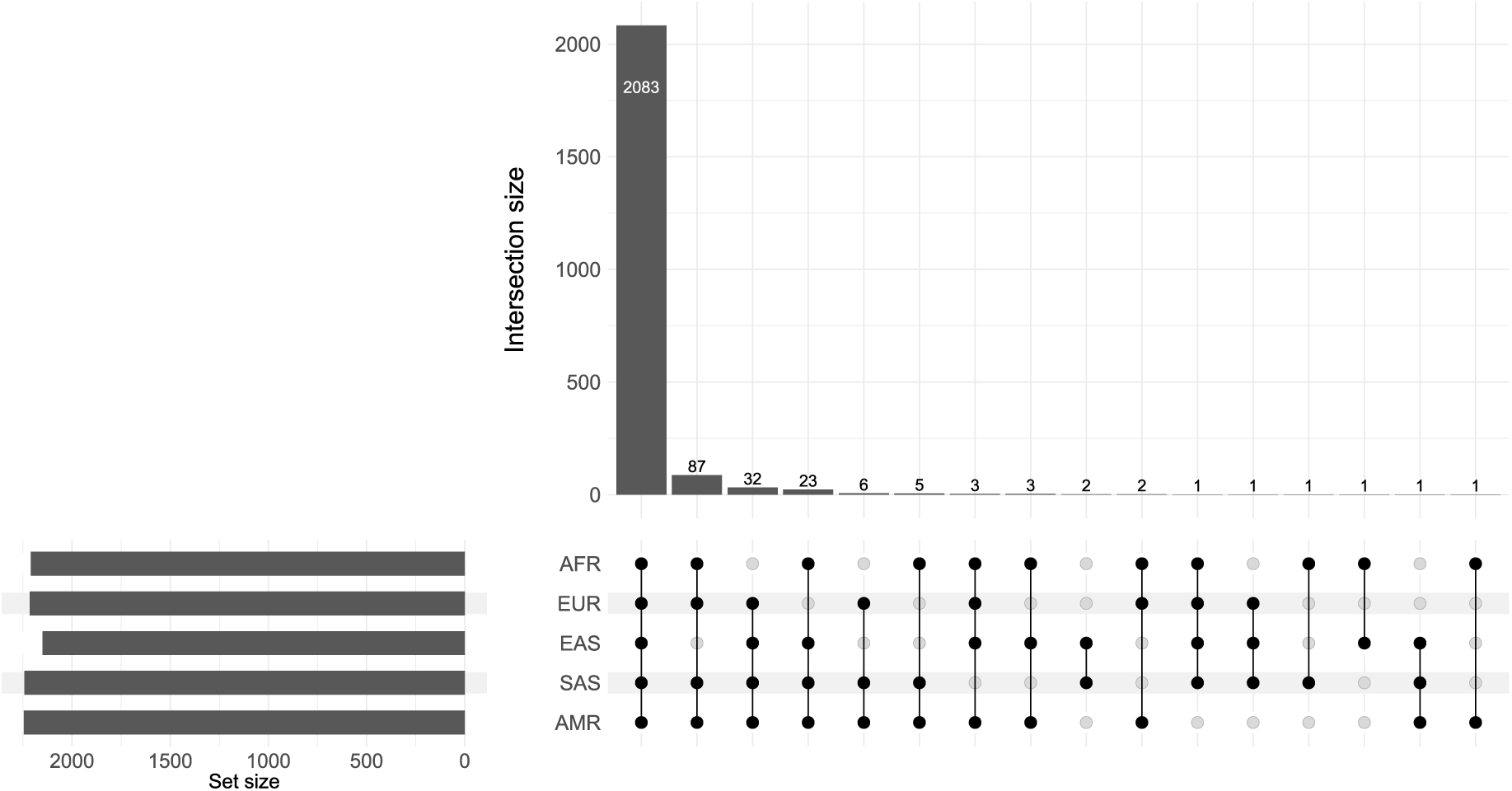
Most ancient SAVs occur in all 1KG superpopulations. Unique and shared ancient SAVs per [38] at Δ *≥* 0.2. Allele frequencies are from 1KG. By definition, each ancient SAV was considered present in a population if the allele frequency was *≥* 0.05 for at least two superpopulations. AFR = African, AMR = American, EAS = East Asian, EUR = European, SAS = South Asian.

**Supplementary Fig. 14.**
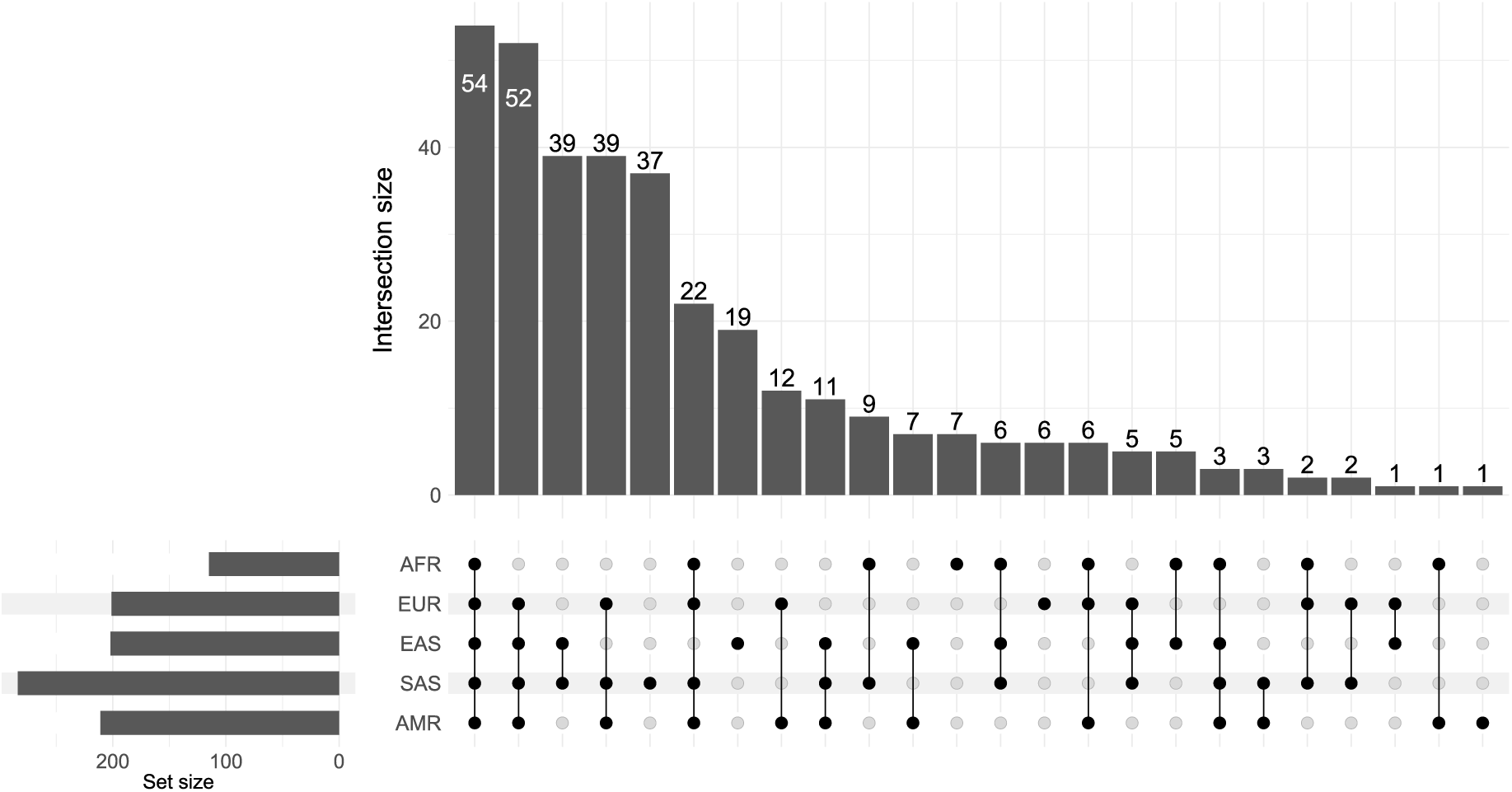
Many introgressed SAVs occur in multiple 1KG superpopulations. Unique and shared introgressed SAVs per [39] at Δ *≥* 0.2. Allele frequencies are from 1KG. If the introgressed allele was the reference allele, we subtracted the 1KG allele frequency from 1. Each SAV was considered present in a population if the allele frequency was *>* 0.01 (N = 349). The remaining 28 variants occurred at *≤* 0.01 allele frequency. AFR = African, AMR = American, EAS = East Asian, EUR = European, SAS = South Asian.

**Supplementary Fig. 15.**
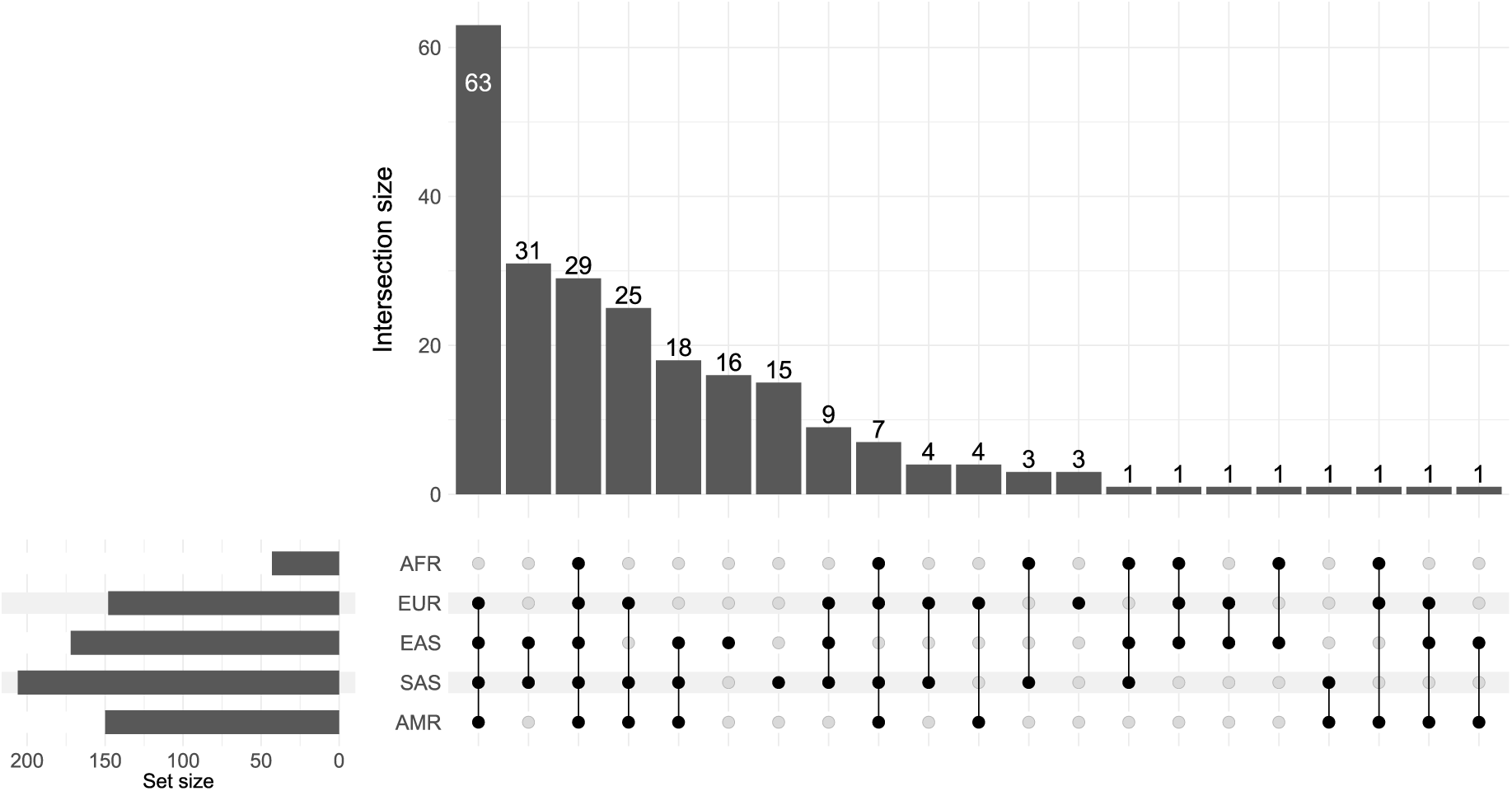
Many introgressed SAVs occur in multiple 1KG superpopulations. Unique and shared introgressed SAVs per [38] at Δ *≥* 0.2. Allele frequencies are from the [38] metadata. Each SAV was considered present in a population if the allele frequency was *>* 0.01 (N = 203). The remaining 34 variants occurred at *≤* 0.01 allele frequency. AFR = African, AMR = American, EAS = East Asian, EUR = European, SAS = South Asian.

**Supplementary Fig. 16.**
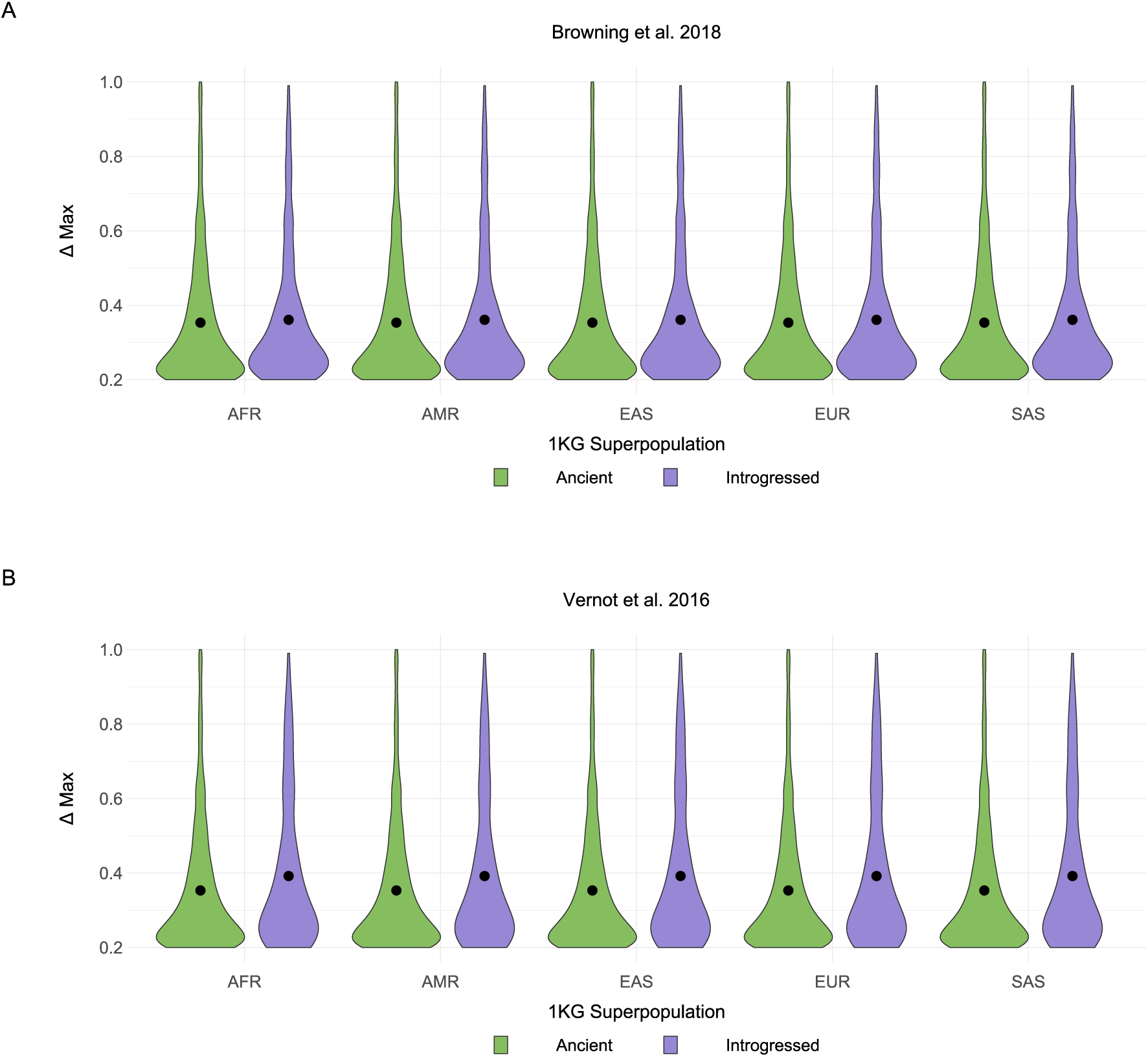
Δ max does not differ between 1KG superpopulations. **(A)** Δ max for SAVs by 1KG superpopulation and allele origin per [39]. AFR = African, AMR = American, EAS = East Asian, EUR = European, SAS = South Asian. **(B)** Δ max for SAVs by 1KG superpopulation and allele origin per [38].

**Supplementary Fig. 17.**
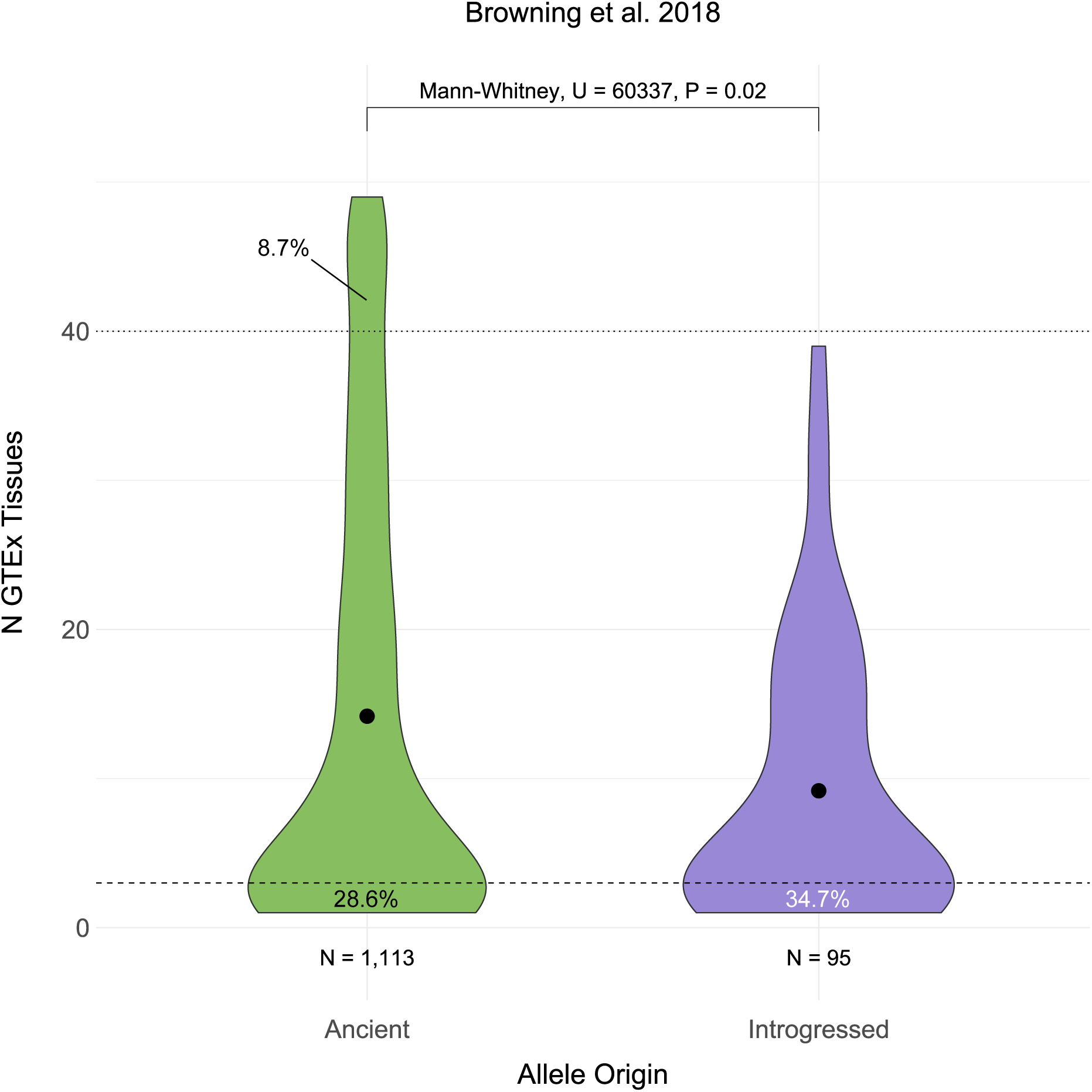
Introgressed sQTL SAVs are more tissue-specific than ancient variants. The distribution of GTEx tissues in which an ancient or introgressed SAV per [39] was identified as an sQTL. We defined “tissue-specific” variants as those occurring in 1 or 2 tissues and “core” sQTLs as those occurring in > 40 of the 49 tissues. The dashed and dotted lines represent these definitions, respectively. The proportion of SAVs below and above these thresholds for both origins are annotated.

**Supplementary Fig. 18.**
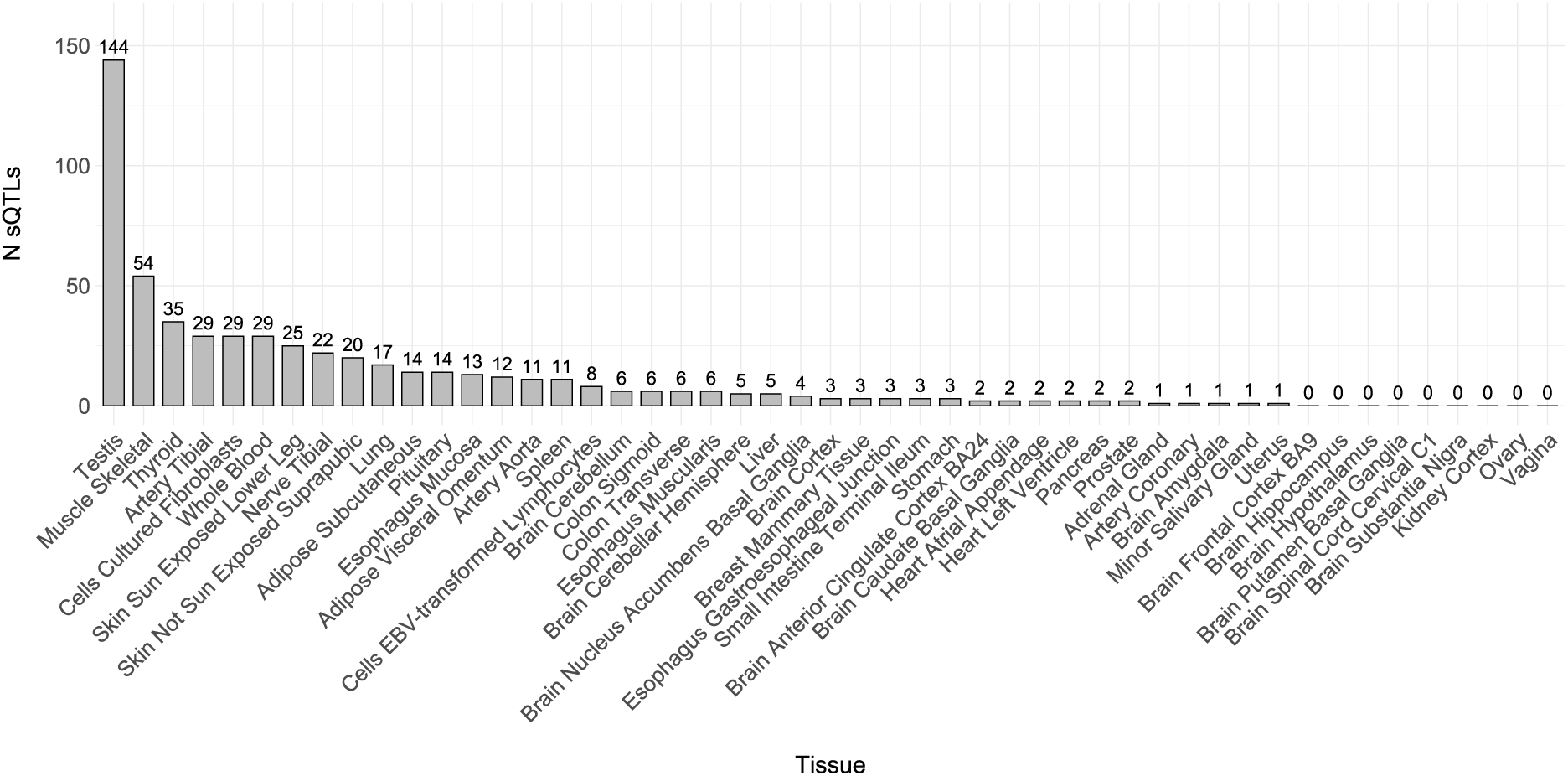
sQTL SAVs predominantly occur in testis, muscle skeletal, thyroid, tibial artery, fibroblasts, skin, and tibial nerve tissues. The number of tissue-specific (N GTEx tissues = 1 or 2) sQTL SAVs per GTEx tissue.

**Supplementary Fig. 19.**
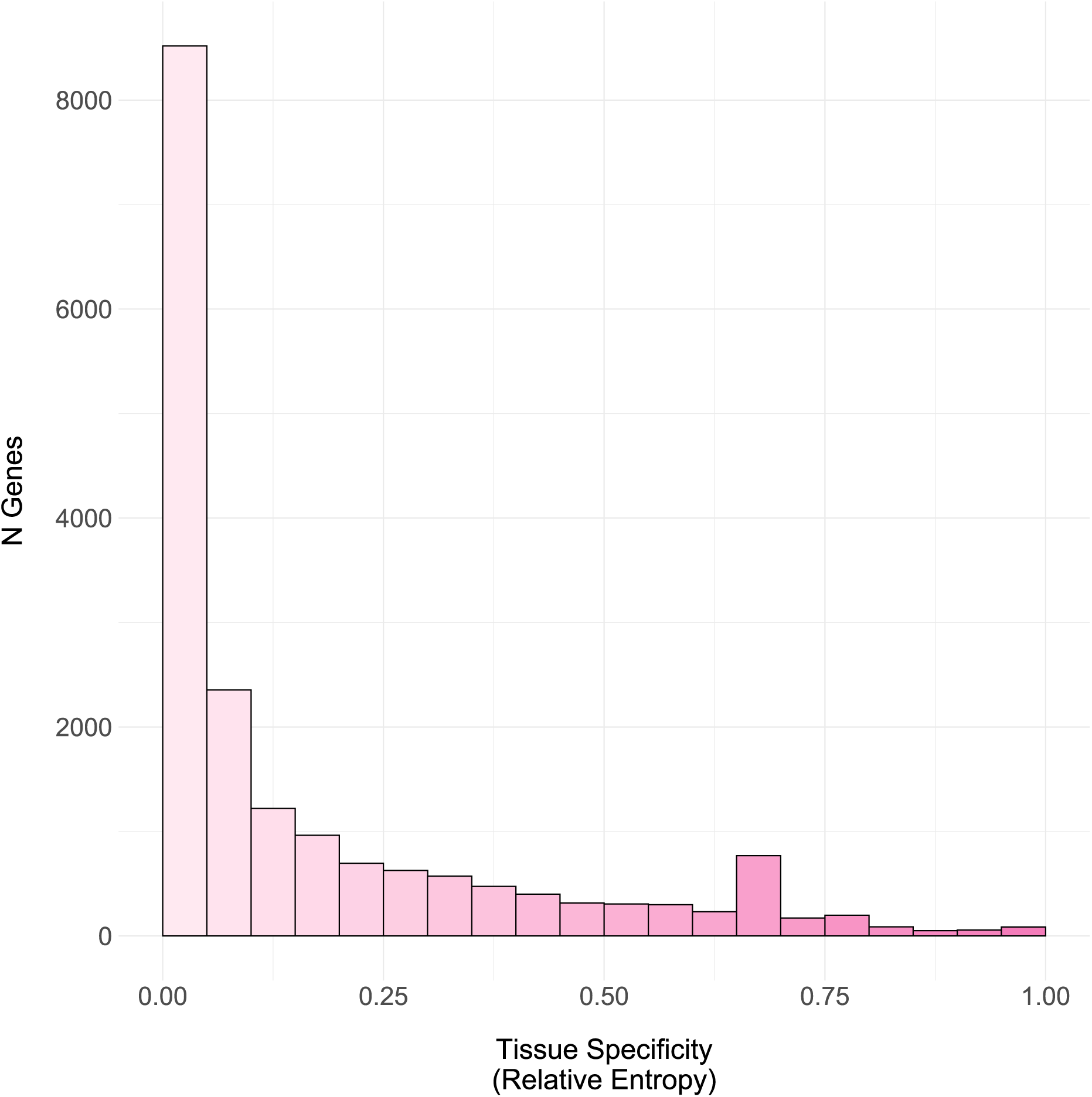
Most gene expression is not tissue-specific. The distribution of relative entropy in 0.05 bins calculated from GTEx TPM counts across 34 tissues for 18,392 genes.

**Supplementary Fig. 20.**
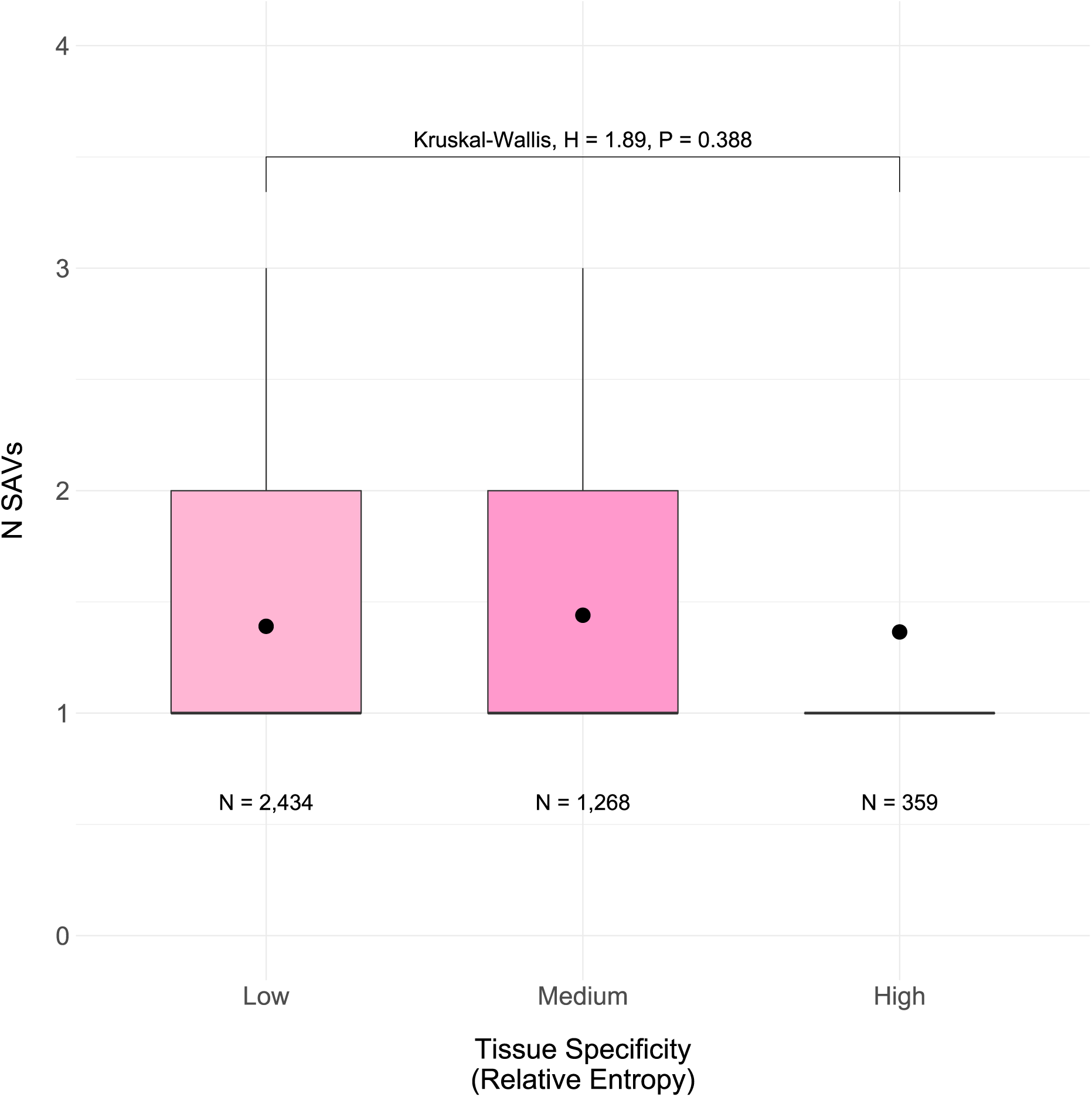
The number of SAVs is unrelated to tissue-specific gene expression. The number of SAVs by binned gene expression calculated from GTEx TPM counts across 34 tissues. We analyzed a total of 4,061 genes with SAVs. Low tissue specificity reflects relative entropy *≤* 0.1, medium tissue specificity reflects relative entropy *>* 0.1 and *≤* 0.5, and high tissue specificity reflects relative entropy *>* 0.5 (**Supplementary** Fig. 19).

**Supplementary Fig. 21.**
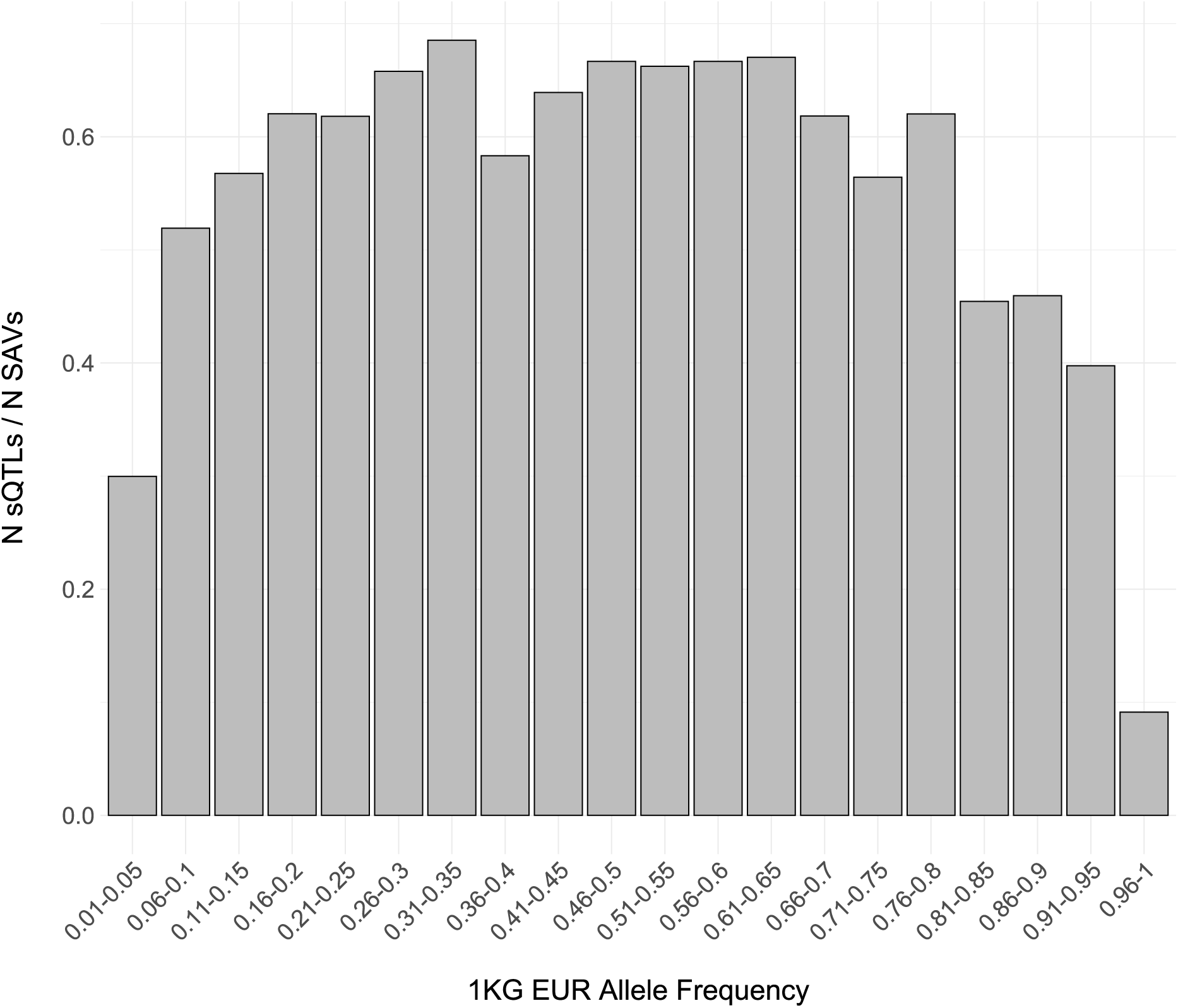
*>* 50% of SAVs are sQTLs. The proportion of sQTLs also identified as SAVs per binned allele frequencies among 1KG European individuals. We used the precalculated frequencies provided in the 1KG VCFs for this analysis.

**Supplementary Fig. 22.**
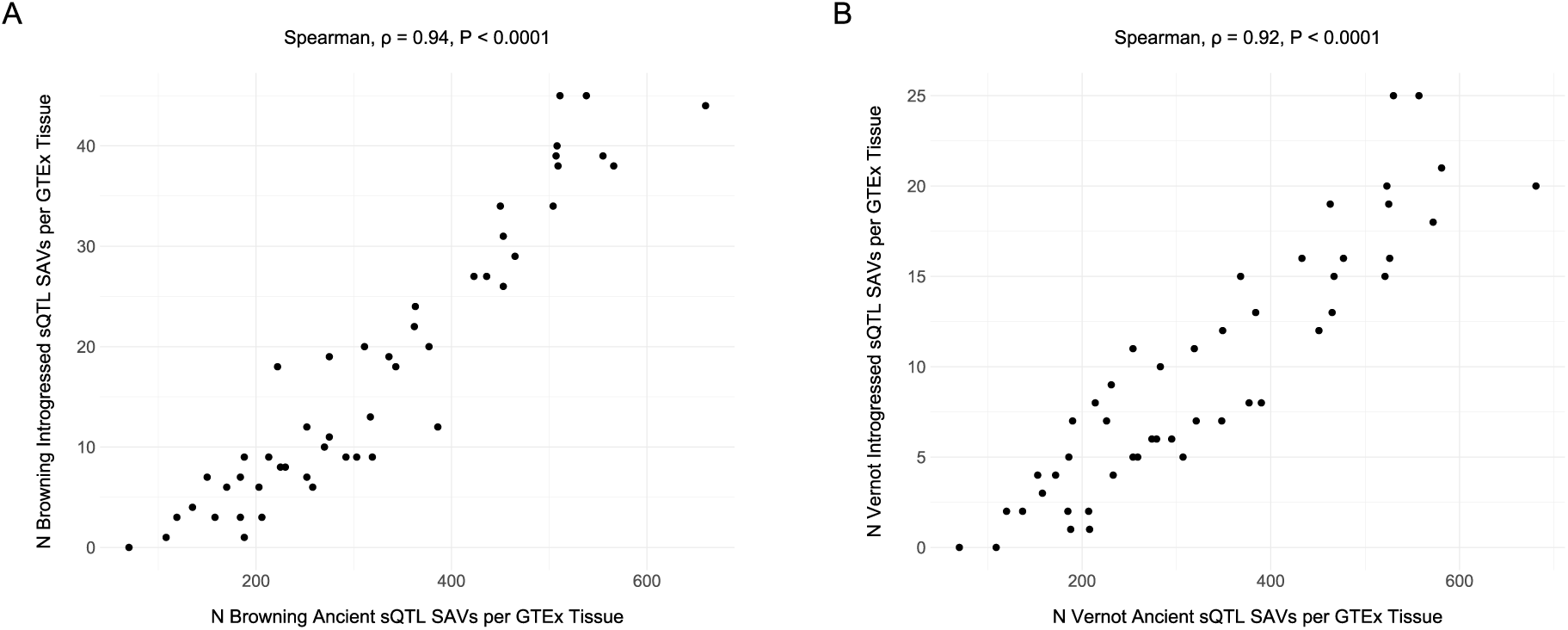
The number of ancient and introgressed sQTL SAVs are associated among GTEx tissues. **(A)** The number of [39] ancient vs. introgressed sQTL SAVs per tissue in GTEx. **(B)** The number of [38] ancient vs. introgressed sQTL SAVs per tissue in GTEx.

**Supplementary Fig. 23.**
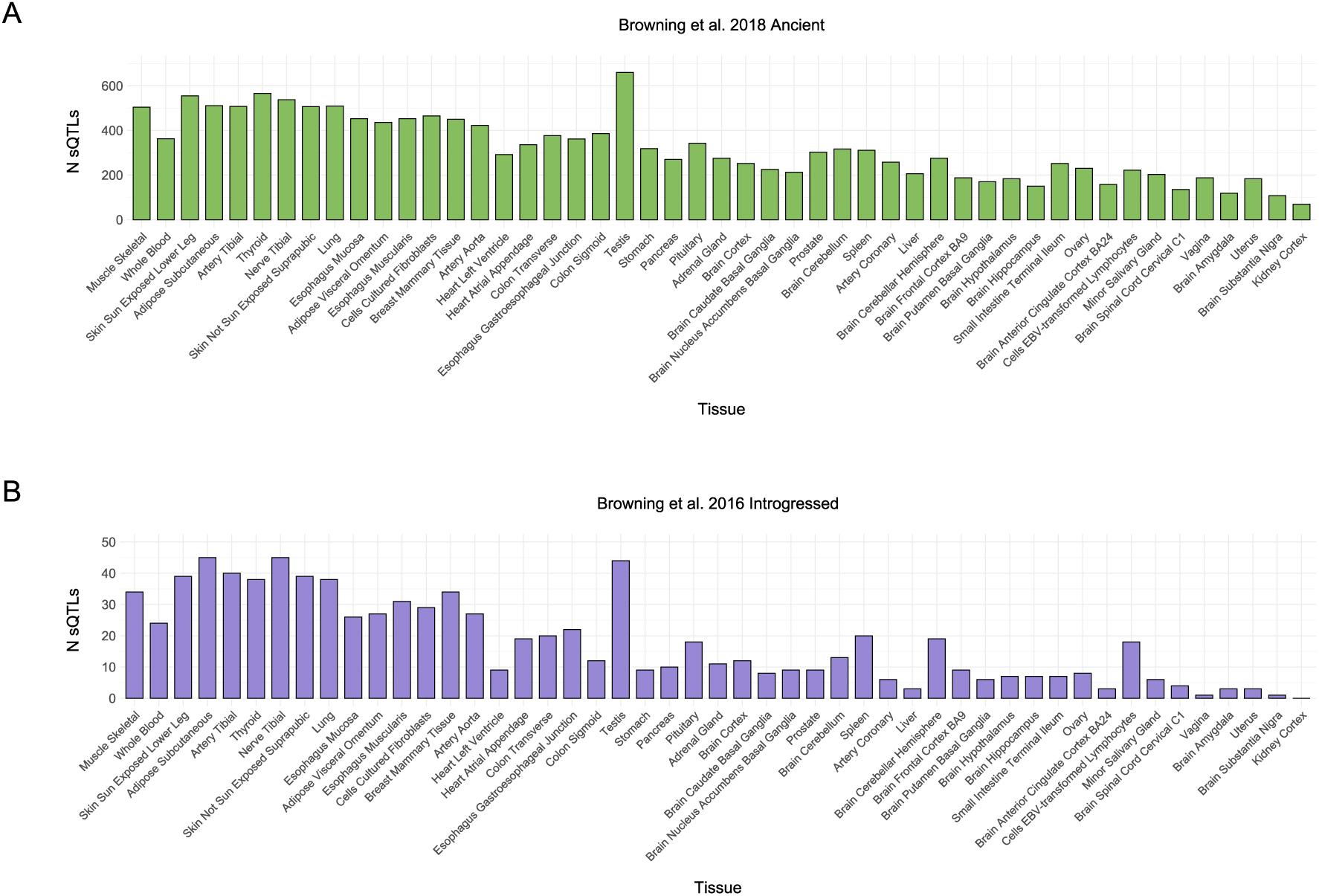
The number of Browning et al. 2018 sQTL SAVs is associated with GTEx sample size. **(A)** The number of sQTL SAVs per tissue among [39] ancient SAVs ordered by decreasing GTEx sample size. **(B)** The number of sQTL SAVs per tissue among [39] introgressed SAVs ordered by decreasing GTEx sample size.

**Supplementary Fig. 24.**
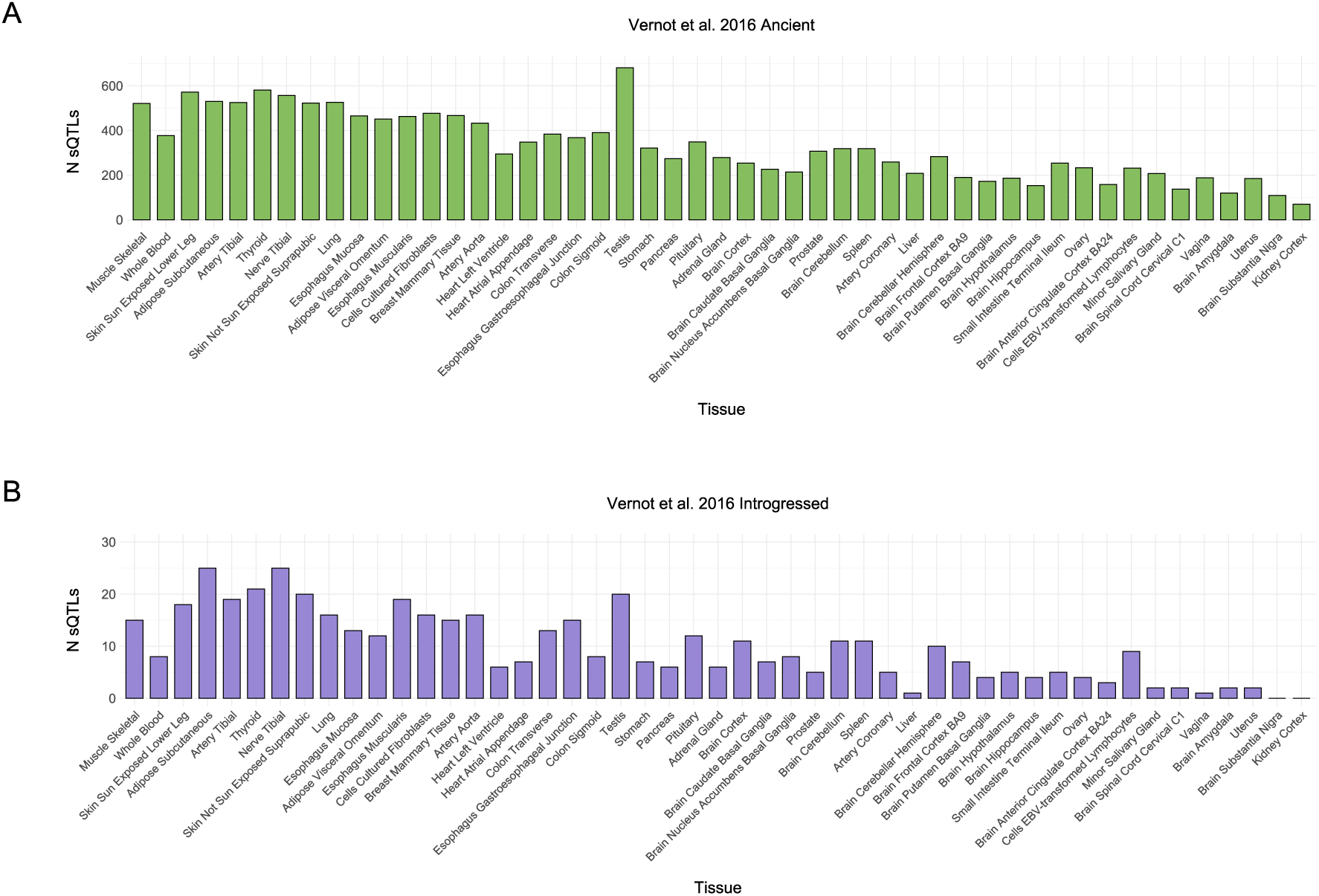
The number of Vernot et al. 2016 sQTL SAVs is associated with GTEx sample size. **(A)** The number of sQTL SAVs per tissue among [38] ancient SAVs ordered by decreasing GTEx sample size. **(B)** The number of sQTL SAVs per tissue among [38] introgressed SAVs ordered by decreasing GTEx sample size.

**Supplementary Fig. 25.**
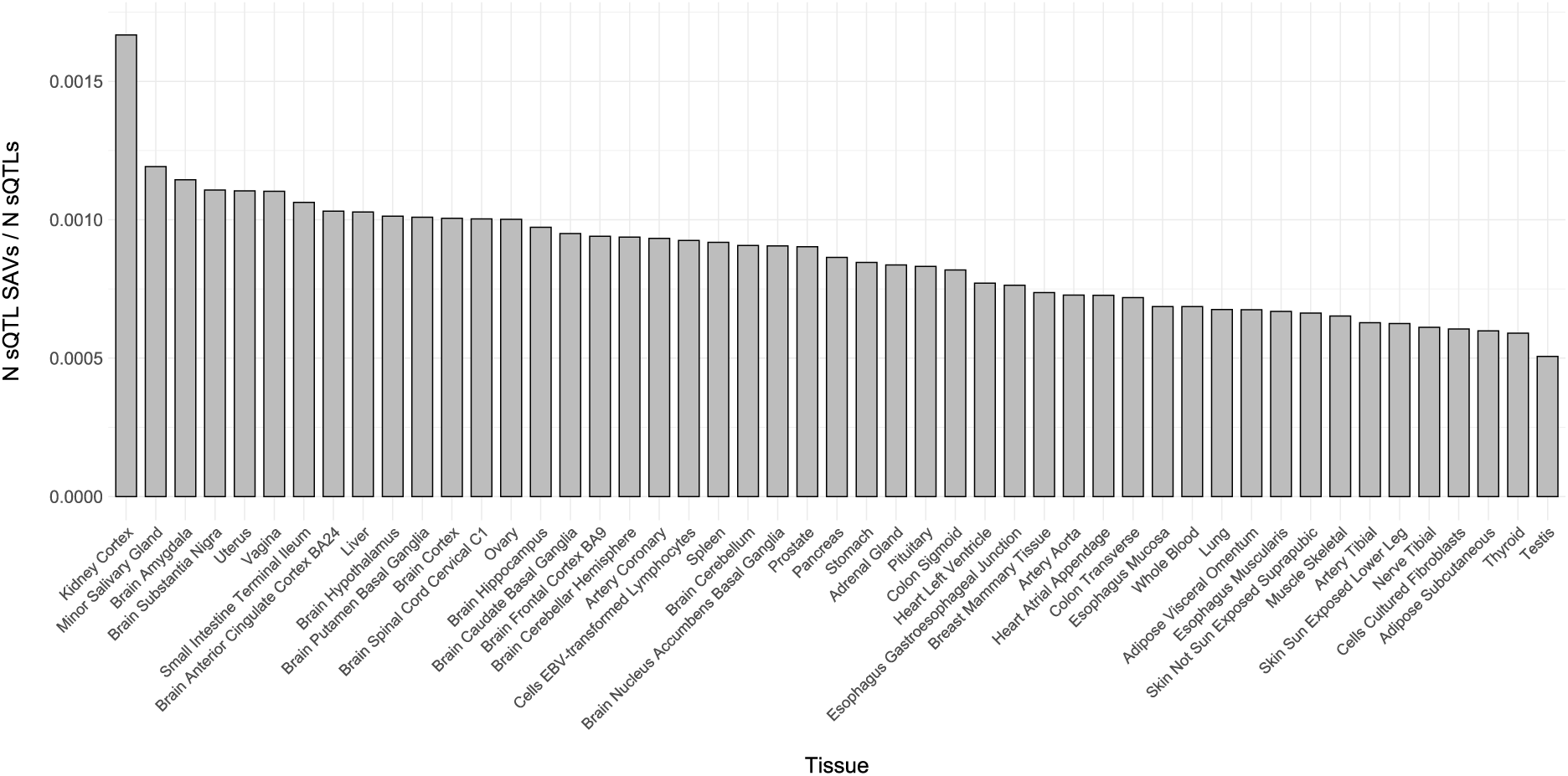
The proportion of sQTL SAVs is similar across tissues. The proportion of sQTLs SAVs from all sQTLs per GTEx tissue, ordered by decreasing proportion.

**Supplementary Fig. 26.**
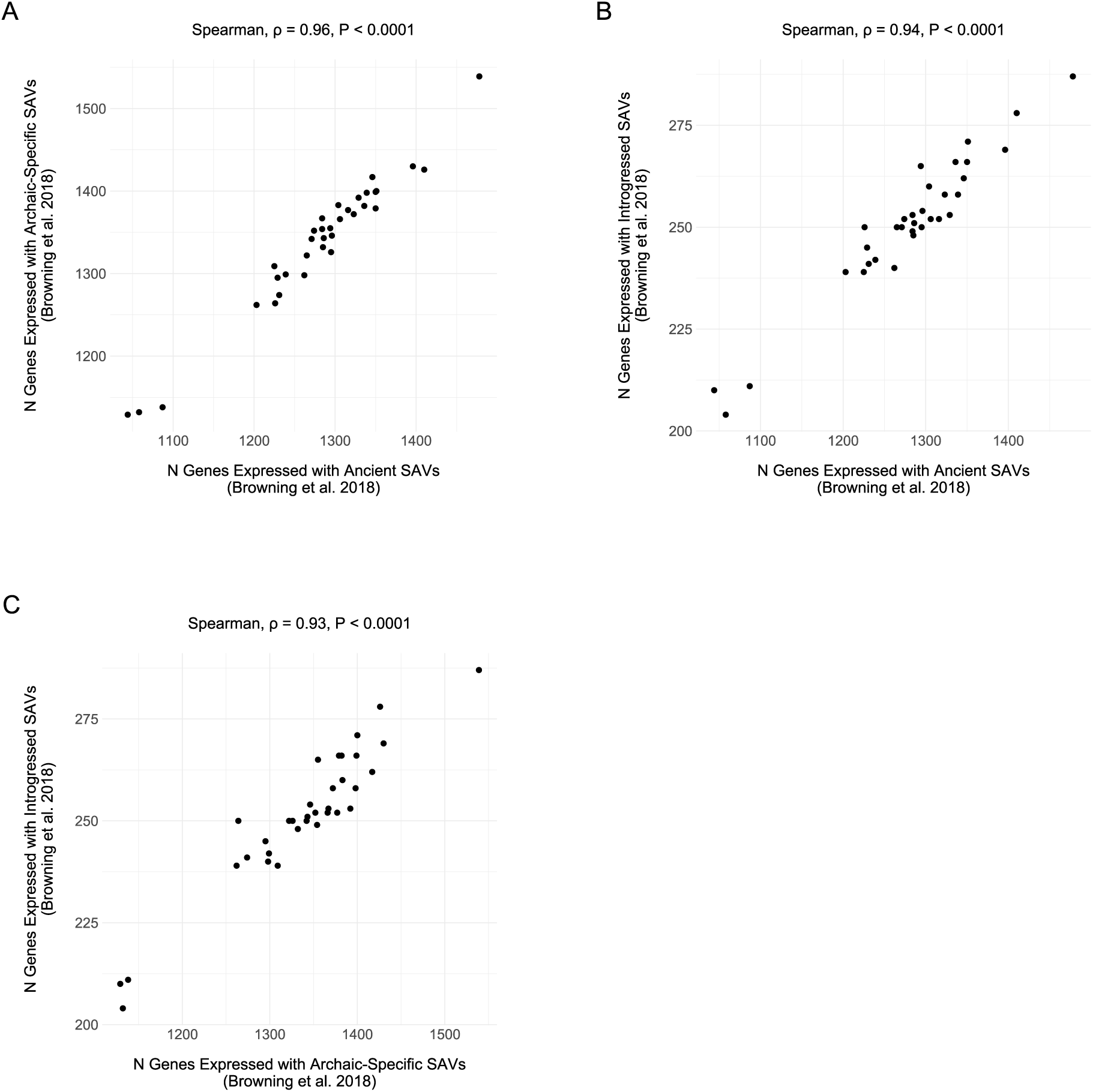
The number of genes expressed per tissue are similar across different [39] allele origins. **(A)** The number of genes with TPM > 1 containing at least one SAV for [39] ancient and archaic-specific SAVs. Each dot represents a GTEx tissue. **(B)** The number of genes with TPM > 1 containing at least one SAV for [39] ancient and introgressed SAVs. **(C)** The number of genes with TPM > 1 containing at least one SAV for [39] archaic-specific and introgressed SAVs.

**Supplementary Fig. 27.**
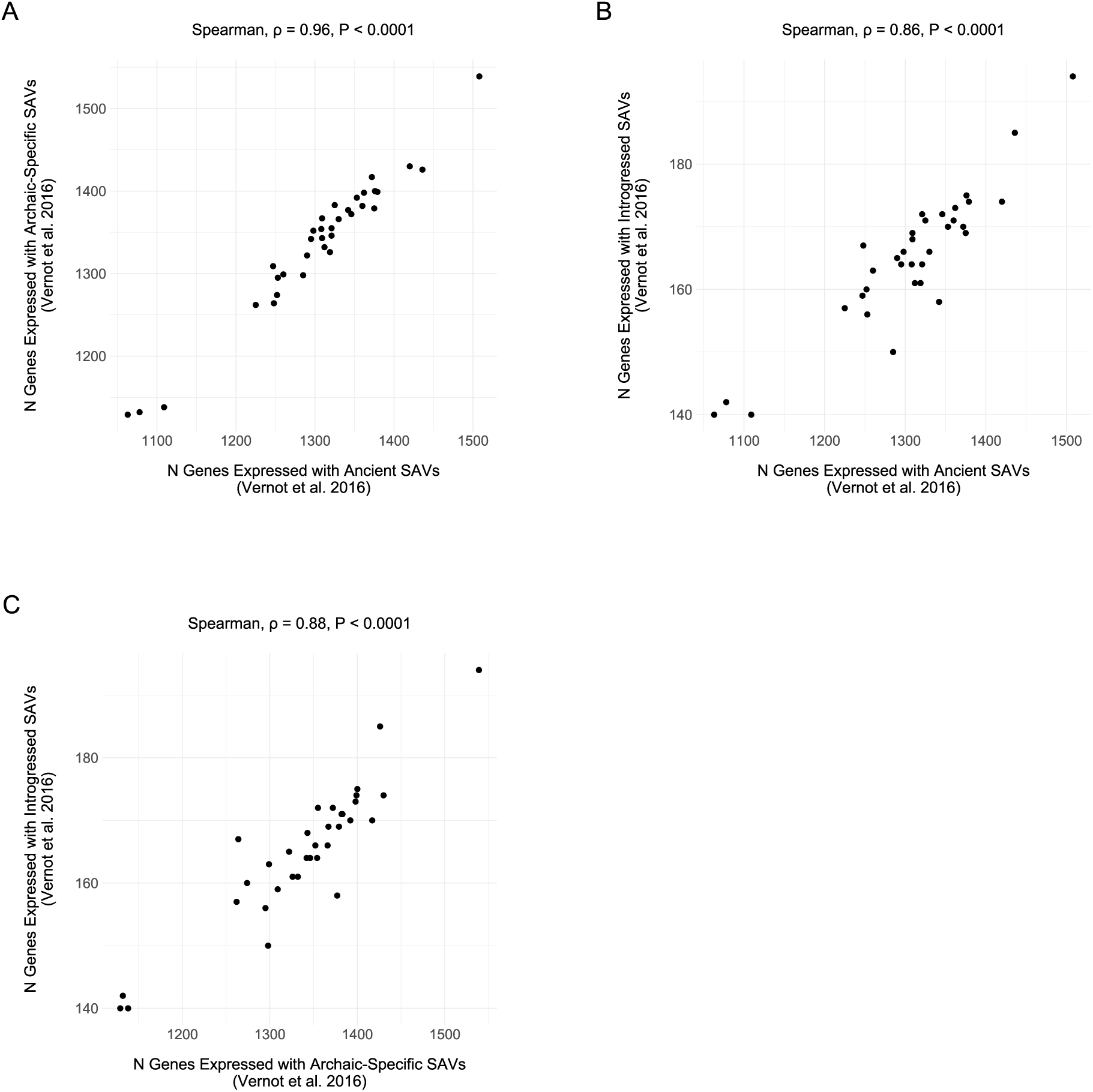
The number of genes expressed per tissue are similar across different [38] allele origins. **(A)** The number of genes with TPM > 1 containing at least one SAV for [38] ancient and archaic-specific SAVs. Each dot represents a GTEx tissue. **(B)** The number of genes with TPM > 1 containing at least one SAV for [38] ancient and introgressed SAVs. **(C)** The number of genes with TPM > 1 containing at least one SAV for [38] archaic-specific and introgressed SAVs.

### 5.4 Supplementary Tables

**Supplementary Table 1.** Distribution of non-reference alleles. The number of variant positions with at least one non-reference allele among the archaics per number of non-reference alleles.

| N Non-Reference Alleles | N |
| --- | --- |
| 1 | 1,566,233 |
| 2 | 1,660 |
| 3 | 1 |

**Supplementary Table 2.**
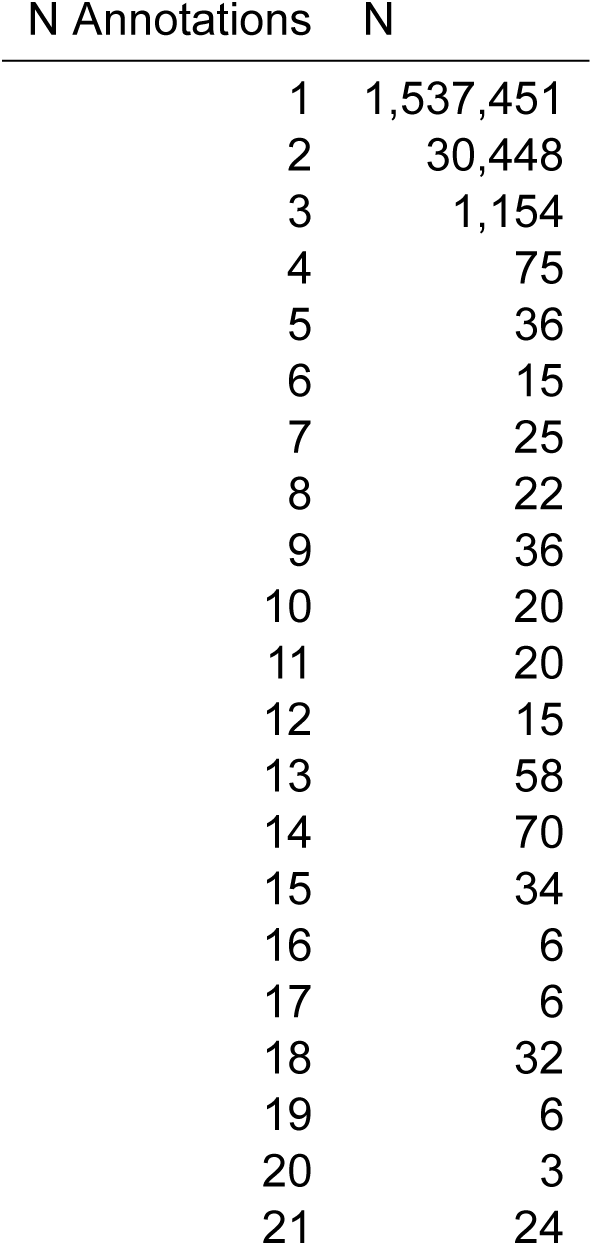
Distribution of multiple annotations. The number of variants with the given number of annotations from GENCODE, Human Release 24, for hg19/GRCH37.

**Supplementary Table 3.**
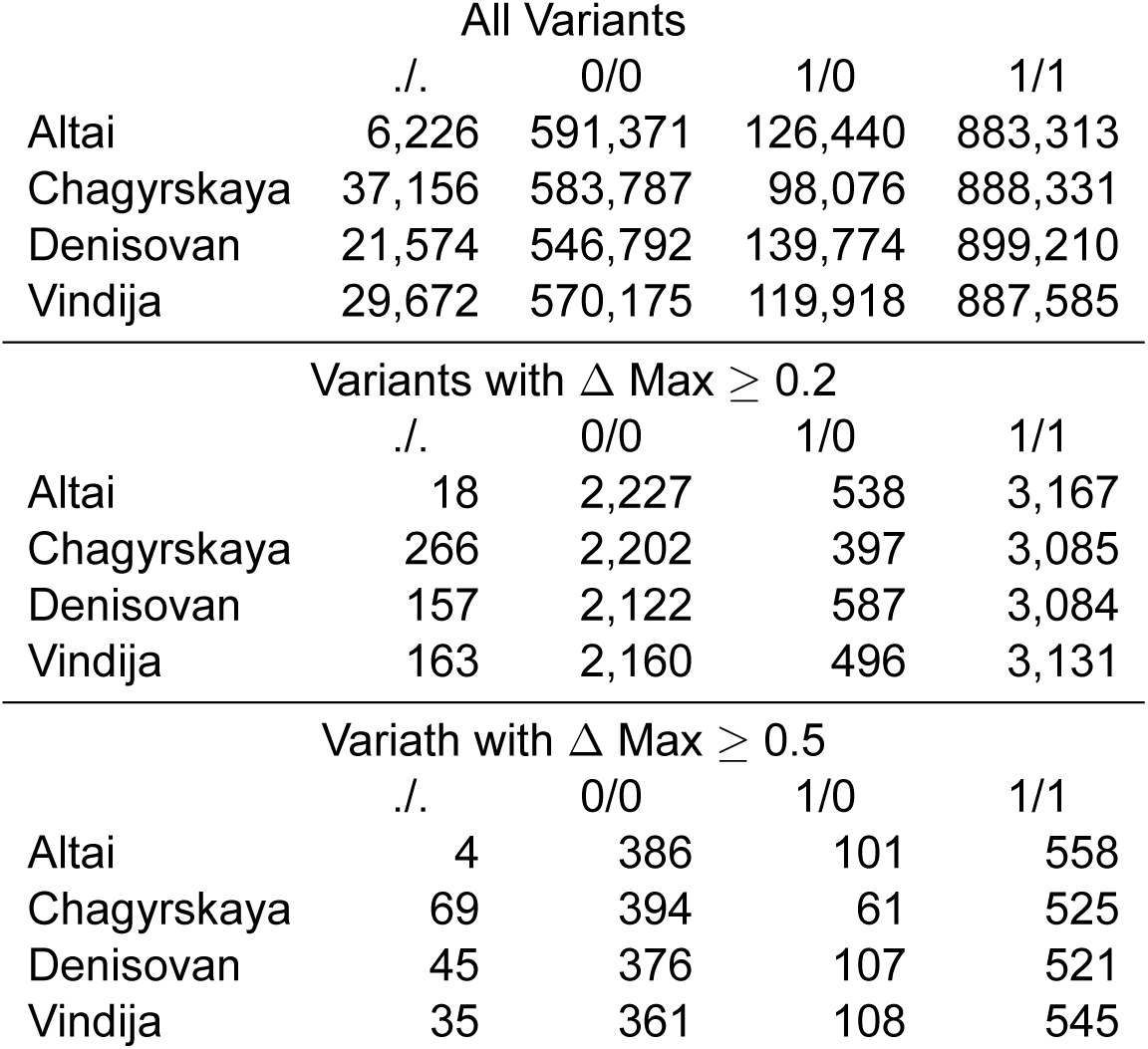
Archaic genotype distribution. The number of genotypes per individual for all variants and both delta thresholds. ./. = missing genotypes, 0/0 = homozygous for reference allele, 1/0 = heterozygote, 1/1 = homozygous for alternate allele.

|  | All Variants |  |  |  |
| --- | --- | --- | --- | --- |
|  | ./. | 0/0 | 1/0 | 1/1 |
| Altai | 6,226 | 591,371 | 126,440 | 883,313 |
| Chagyrskaya | 37,156 | 583,787 | 98,076 | 888,331 |
| Denisovan | 21,574 | 546,792 | 139,774 | 899,210 |
| Vindija | 29,672 | 570,175 | 119,918 | 887,585 |
| Variants with $\Delta \text{Max} \geq 0.2$ | | | | |
|  | ./. | 0/0 | 1/0 | 1/1 |
| Altai | 18 | 2,227 | 538 | 3,167 |
| Chagyrskaya | 266 | 2,202 | 397 | 3,085 |
| Denisovan | 157 | 2,122 | 587 | 3,084 |
| Vindija | 163 | 2,160 | 496 | 3,131 |
| Variants with $\Delta \text{Max} \geq 0.5$ | | | | |
|  | ./. | 0/0 | 1/0 | 1/1 |
| Altai | 4 | 386 | 101 | 558 |
| Chagyrskaya | 69 | 394 | 61 | 525 |
| Denisovan | 45 | 376 | 107 | 521 |
| Vindija | 35 | 361 | 108 | 545 |

**Supplementary Table 4.** The average number of SAVs per randomly sampled individual in 1KG. We randomly sampled one individual per 1KG population and ran SpliceAI on their combined variants. There were 4,582,422 total SpliceAI annotated variants among autosomal SNVs and 14,006 variants with a Δ max *≥* 0.2. These N SAVs reported here include any site where an individual had at least one alternate allele present (i.e., heterozygotes and homozygotes for the alternate allele). Superpopulation and population labels follow 1KG convention.

| 1KG Superpopulation | 1KG Population | Sample Name | N SAVs |
| --- | --- | --- | --- |
| AFR | ESN | HG00137 | 3,266 |
|  | GWD | HG00338 | 3,344 |
|  | LWK | HG00619 | 3,320 |
|  | MSL | HG01198 | 3,514 |
|  | YRI | HG01281 | 3,493 |
| AMR | CLM | HG01524 | 3,363 |
|  | MXL | HG01802 | 3,363 |
|  | PEL | HG02142 | 3,349 |
|  | PUR | HG02345 | 3,385 |
| EAS | CDX | HG02629 | 4,120 |
|  | CHB | HG03060 | 4,160 |
|  | CHS | HG03190 | 4,142 |
|  | JPT | HG03708 | 3,368 |
|  | KHV | HG03711 | 3,379 |
| EUR | CEU | HG03800 | 3,435 |
|  | FIN | HG04014 | 3,421 |
|  | GBR | NA11830 | 3,200 |
|  | GIH | NA18552 | 3,382 |
|  | IBS | NA18868 | 4,102 |
|  | TSI | NA19011 | 3,321 |
| SAS | BEB | NA19452 | 4,078 |
|  | ITU | NA19741 | 3,284 |
|  | PJL | NA20537 | 3,168 |
|  | STU | NA21141 | 3,487 |

**Supplementary Table 5.** Distribution of SAVs per gene. The distribution of SAVs per gene with the specific genes listed for those with *≥* 7 SAVs.

| N SAVs | N Genes | Genes |
| --- | --- | --- |
| 1 | 3,111 |  |
| 2 | 769 |  |
| 3 | 246 |  |
| 4 | 71 |  |
| 5 | 19 |  |
| 6 | 14 |  |
| 7 | 6 | *CDH13, CDH23, CSMD1, DNAH17, HLA-DPA1, LAMA5* |
| 8 | 1 | *ADARB2* |
| 9 | 2 | *SDK1, WWOX* |
| 10 | 1 | *HLA-DPB1* |
| 11 | 2 | *CNTNAP2, SSPO* |

**Supplementary Table 6.** Physical gene characteristic associations with N SAVs per gene. Spearman correlation between four variables: 1) the number of exons, 2) length of coding sequence in bp, 3) gene body length in bp, 4) the number of isoforms and the N SAVs per gene for both Δ thresholds.

| $\Delta$ | Gene Characteristic | $\rho$ | P Value |
| --- | --- | --- | --- |
| 0.2 | N Exons | 0.3161 | 0 |
|  | CDS/Exon Length | 0.1851 | 1.07E-135 |
|  | Gene Length | 0.2920 | 0 |
|  | N Isoforms | 0.1767 | 1.87E-113 |
| 0.5 | N Exons | 0.1330 | 2.20E-70 |
|  | CDS/Exon Length | 0.0625 | 1.01E-16 |
|  | Gene Length | 0.1017 | 8.99E-42 |
|  | N Isoforms | 0.0769 | 1.24E-22 |

**Supplementary Table 7.**
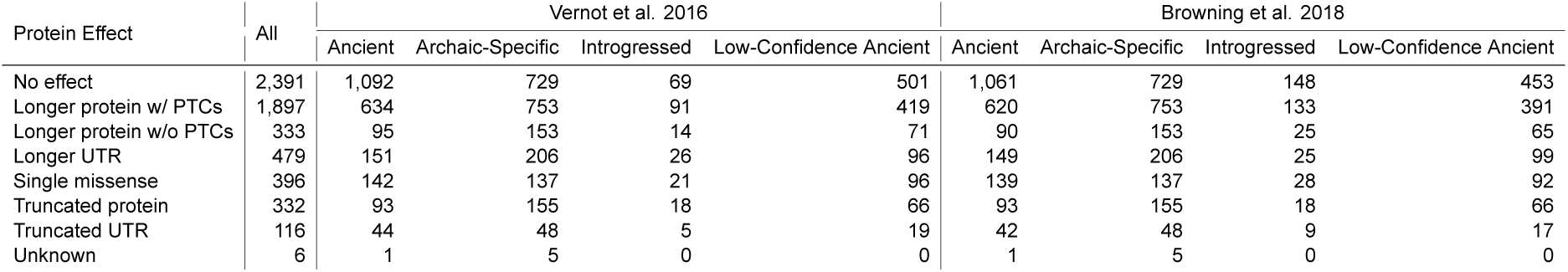
SAV effects on resulting protein. The number of SAVs per effect on the resulting transcript and protein. PTC = premature termination codon, UTR = 5’ or 3’ untranslated region.

**Supplementary Table 8.** Enrichment tests for the sum of lineage-specific SAVs at different Δs. Input data for Fisher’s exact tests. OR = odds ratio; 95% CI Lower Bound and 95% CI Upper Bound = the lower and upper bounds of the 95% confidence interval, respectively; P Value = unadjusted p-value.

| Delta | N Unique SAVs | N Unique Non SAVs | N Shared SAVs | N Shared Non SAVs | OR | 95% CI Lower Bound | 95% CI Upper Bound | P Value | Bonferroni |
| --- | --- | --- | --- | --- | --- | --- | --- | --- | --- |
| 0.2 | 2,416 | 615,666 | 1,933 | 571,264 | 1.160 | 1.092 | 1.231 | 0.000 | Y |
| 0.3 | 1,175 | 616,907 | 945 | 572,252 | 1.153 | 1.059 | 1.257 | 0.001 | Y |
| 0.4 | 685 | 617,397 | 533 | 572,664 | 1.192 | 1.064 | 1.335 | 0.002 | Y |
| 0.5 | 437 | 617,645 | 328 | 572,869 | 1.236 | 1.071 | 1.426 | 0.004 | Y |

**Supplementary Table 9.**
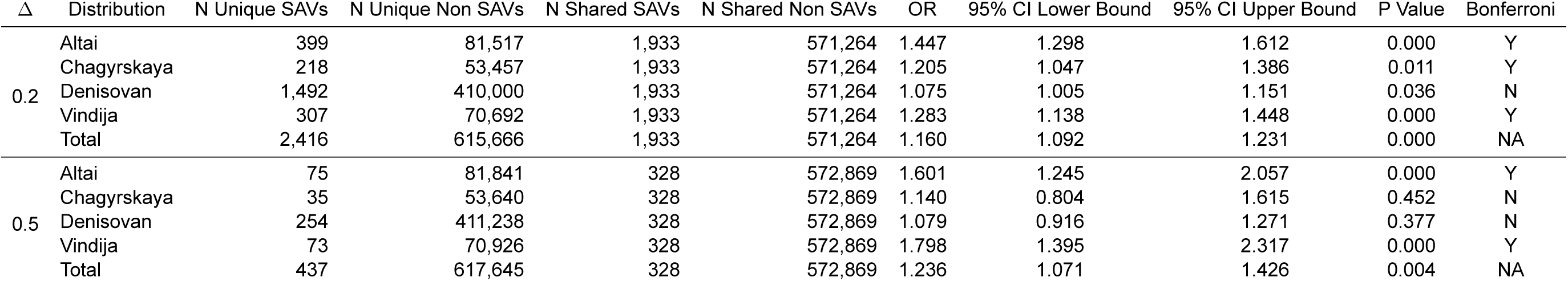
Enrichment tests for lineage-specific SAVs. Input data for Fisher’s exact tests. OR = odds ratio; 95% CI Lower Bound and 95% CI Upper Bound = the lower and upper bounds of the 95% confidence interval, respectively; P Value = unadjusted p-value.

**Supplementary Table 10.**
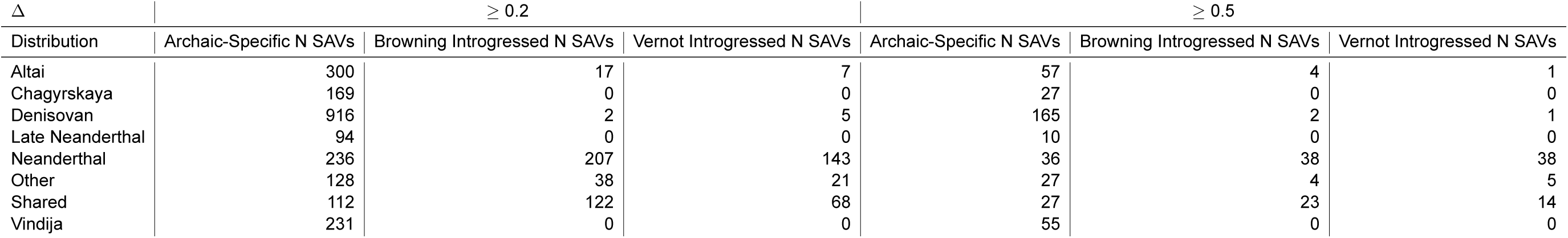
Distribution of archaic-specific and introgressed SAVs. N SAVs for both Δ thresholds among archaic-specific and introgressed SAVs per [39] and [38]. Other denotes any combination of archaics not already listed.

**Supplementary Table 11.** SAVs that exhibit allele-specific expression in modern humans. 16 SAVs from our dataset that matched variants from [59].

| Chromosome | Position | Reference Allele | Alternate Allele | Delta Max | Browning Allele Origin | Vernot Allele Origin | Annotation |
| --- | --- | --- | --- | --- | --- | --- | --- |
| chr1 | 22,174,518 | G | T | 0.98 | introgressed | introgressed | HSPG2 |
| chr1 | 55,537,474 | C | G | 0.33 | low-confidence ancient | introgressed | USP24 |
| chr1 | 161,681,848 | C | T | 0.20 | introgressed | introgressed | FCRLA |
| chr1 | 212,985,592 | G | A | 0.52 | introgressed | introgressed | TATDN3 |
| chr11 | 86,159,859 | G | A | 0.26 | ancient | introgressed | ME3 |
| chr12 | 133,272,470 | G | T | 0.26 | introgressed | introgressed | PXMP2 |
| chr12 | 133,272,470 | G | T | 0.26 | introgressed | introgressed | RP13-672B3.2 |
| chr15 | 85,403,496 | G | A | 0.33 | introgressed | introgressed | ALPK3 |
| chr16 | 88,924,425 | C | G | 0.41 | introgressed | introgressed | TRAPPC2L |
| chr19 | 40,913,595 | G | A | 0.23 | introgressed | low-confidence ancient | PRX |
| chr22 | 50,684,852 | G | C | 0.25 | introgressed | introgressed | HDAC10 |
| chr4 | 170,990,750 | G | A | 0.22 | introgressed | introgressed | AADAT |
| chr5 | 6,753,013 | C | T | 0.24 | introgressed | introgressed | PAPD7 |
| chr5 | 167,919,825 | G | A | 0.38 | introgressed | introgressed | RARS |
| chr6 | 33,264,115 | A | C | 0.34 | low-confidence ancient | introgressed | RGL2 |
| chr8 | 130,763,783 | C | T | 0.87 | introgressed | introgressed | GSDMC |

**Supplementary Table 12.** Core sQTL SAV distribution among the archaics. The distribution of core sQTL SAVs (N = 1,145) among the archaics. Core sQTLs were defined as those variants detected in *>* 40 tissues. All the variants represented here are either low or high-confidence ancient.

| Distribution | N sQTLs |
| --- | --- |
| Altai | 2 |
| Chagyrskaya | 3 |
| Denisovan | 8 |
| Late Neanderthal | 2 |
| Neanderthal | 9 |
| Other | 14 |
| Shared | 74 |

**Supplementary Table 13.** Maximum Δ for SAVs with an introgressed reference allele. Δ maximum for 26 SAVs whose reference allele rather than the alternate allele matched an introgressed tag SNP from [38]. We switched the reference and alternate alleles for these (and non-SAV) loci and reran SpliceAI. Delta Max Alternate Allele = the originally predicted Δ where the reference allele is introgressed, Delta Max Reference as Alternate = the Δ when the reference and alternate alleles are switched. Bolded variants are those whose values did not pass the SAP threshold after switching alleles.

| Chromosome | Position | Reference Allele | Alternate Allele | Delta Max Alternate Allele | Delta Max Reference as Alternate |
| --- | --- | --- | --- | --- | --- |
| chr1 | 52,820,680 | C | T | 0.23 | 0.23 |
| chr1 | 152,944,501 | A | T | 0.23 | 0.23 |
| chr1 | 183,750,131 | C | A | 0.26 | 0.26 |
| **chr1** | **216,945,756** | **C** | **A** | **0.20** | **0.19** |
| chr10 | 17,373,518 | T | C | 0.22 | 0.22 |
| chr11 | 84,191,111 | T | C | 0.40 | 0.40 |
| chr11 | 99,345,359 | C | G | 0.67 | 0.67 |
| chr17 | 14,109,360 | T | G | 0.20 | 0.21 |
| chr18 | 60,003,587 | G | A | 0.31 | 0.30 |
| chr19 | 33,410,289 | T | C | 0.71 | 0.71 |
| chr20 | 62,224,595 | G | A | 0.41 | 0.41 |
| chr3 | 52,486,376 | T | C | 0.34 | 0.35 |
| chr4 | 38,805,942 | G | C | 0.38 | 0.37 |
| chr4 | 100,460,531 | A | G | 0.35 | 0.35 |
| chr4 | 135,122,662 | G | A | 0.80 | 0.80 |
| chr5 | 118,669,326 | G | C | 0.22 | 0.22 |
| chr5 | 126,989,007 | A | G | 0.91 | 0.91 |
| chr5 | 148,430,560 | A | C | 0.32 | 0.32 |
| chr6 | 51,889,358 | T | C | 0.42 | 0.42 |
| chr6 | 52,138,226 | C | G | 0.21 | 0.21 |
| chr7 | 119,524,266 | C | T | 0.30 | 0.30 |
| chr7 | 126,797,408 | T | A | 0.43 | 0.43 |
| chr7 | 140,801,296 | T | C | 0.25 | 0.25 |
| chr7 | 140,958,656 | A | G | 0.26 | 0.26 |
| **chr7** | **157,177,273** | **C** | **T** | **0.31** | **0.16** |
| chr7 | 158,543,832 | C | A | 0.21 | 0.22 |

**Supplementary Table 14.** Missense variants among 147 spliceosome genes. The number of variant positions with at least one non-reference allele among all four archaics and 24 randomly sampled 1KG samples that was identified as a missense variant by Ensembl’s Variant Effect Predictor. Superpopulation and population labels follow 1KG convention. N transcripts = number of transcripts with a missense variant, N Variants = number of missense variant, N Variants PolyPhen = number of damaging variants as measured by PolyPhen, N Variants SIFT = number of deleterious variants as measured by SIFT, N Variants Both = number of variants that were considered both damaging and deleterious by PolyPhen and SIFT, respectively.

| 1KG Superpopulation | 1KG Population | Sample | N Transcripts | N Variants | N Variants PolyPhen | N Variants SIFT | N Variants Both |
| --- | --- | --- | --- | --- | --- | --- | --- |
|  |  | Altai | 34 | 19 | 2 | 5 | 2 |
|  |  | Chagyrskaya | 27 | 15 | 1 | 2 | 0 |
|  |  | Denisovan | 43 | 25 | 2 | 5 | 0 |
|  |  | Vindija | 24 | 14 | 1 | 1 | 0 |
| AFR | ESN | HG00137 | 71 | 23 | 3 | 5 | 2 |
|  | GWD | HG00338 | 57 | 23 | 1 | 3 | 1 |
|  | LWK | HG00619 | 68 | 27 | 2 | 2 | 1 |
|  | MSL | HG01198 | 67 | 28 | 2 | 6 | 1 |
|  | YRI | HG01281 | 56 | 22 | 2 | 2 | 1 |
| AMR | CLM | HG01524 | 69 | 31 | 3 | 7 | 3 |
|  | MXL | HG01802 | 42 | 21 | 0 | 3 | 0 |
|  | PEL | HG02142 | 49 | 19 | 0 | 1 | 0 |
|  | PUR | HG02345 | 43 | 22 | 0 | 3 | 0 |
| EAS | CDX | HG02629 | 70 | 26 | 1 | 6 | 1 |
|  | CHB | HG03060 | 59 | 23 | 1 | 6 | 0 |
|  | CHS | HG03190 | 56 | 23 | 1 | 4 | 1 |
|  | JPT | HG03708 | 66 | 24 | 0 | 3 | 0 |
|  | KHV | HG03711 | 52 | 25 | 1 | 5 | 1 |
| EUR | CEU | HG03800 | 55 | 17 | 1 | 3 | 1 |
|  | FIN | HG04014 | 59 | 24 | 1 | 2 | 1 |
|  | GBR | NA11830 | 83 | 26 | 1 | 5 | 1 |
|  | GIH | NA18552 | 56 | 24 | 1 | 5 | 1 |
|  | IBS | NA18868 | 53 | 21 | 0 | 3 | 0 |
|  | TSI | NA19011 | 59 | 25 | 0 | 4 | 0 |
| SAS | BEB | NA19452 | 65 | 27 | 2 | 6 | 1 |
|  | ITU | NA19741 | 83 | 29 | 1 | 7 | 1 |
|  | PJL | NA20537 | 41 | 23 | 1 | 4 | 1 |
|  | STU | NA21141 | 56 | 23 | 1 | 5 | 1 |

**Supplementary Table 15.** sQTL SAVs by allele origin. The number of sQTL SAVs by allele origin for 49 tissues in GTEx, v8.

| Tissue | N Vernot Ancient | N Vernot Introgressed | N Browning Ancient | N Browning Introgressed | GTEx Sample Size |
| --- | --- | --- | --- | --- | --- |
| Adipose Subcutaneous | 530 | 25 | 511 | 45 | 663 |
| Adipose Visceral Omentum | 451 | 12 | 436 | 27 | 541 |
| Adrenal Gland | 279 | 6 | 275 | 11 | 258 |
| Artery Aorta | 433 | 16 | 423 | 27 | 432 |
| Artery Coronary | 259 | 5 | 258 | 6 | 240 |
| Artery Tibial | 525 | 19 | 508 | 40 | 663 |
| Brain Amygdala | 120 | 2 | 119 | 3 | 152 |
| Brain Anterior cingulate cortex BA24 | 158 | 3 | 158 | 3 | 176 |
| Brain Caudate basal ganglia | 226 | 7 | 225 | 8 | 246 |
| Brain Cerebellar Hemisphere | 283 | 10 | 275 | 19 | 215 |
| Brain Cerebellum | 319 | 11 | 317 | 13 | 241 |
| Brain Cortex | 254 | 11 | 252 | 12 | 255 |
| Brain Frontal Cortex BA9 | 190 | 7 | 188 | 9 | 209 |
| Brain Hippocampus | 153 | 4 | 150 | 7 | 197 |
| Brain Hypothalamus | 186 | 5 | 184 | 7 | 202 |
| Brain Nucleus accumbens basal ganglia | 214 | 8 | 213 | 9 | 246 |
| Brain Putamen basal ganglia | 172 | 4 | 170 | 6 | 205 |
| Brain Spinal cord cervical c-1 | 137 | 2 | 135 | 4 | 159 |
| Brain Substantia nigra | 109 | 0 | 108 | 1 | 139 |
| Breast Mammary Tissue | 467 | 15 | 450 | 34 | 459 |
| Cells Cultured fibroblasts | 477 | 16 | 465 | 29 | 504 |
| Cells EBV-transformed lymphocytes | 231 | 9 | 222 | 18 | 174 |
| Colon Sigmoid | 390 | 8 | 386 | 12 | 373 |
| Colon Transverse | 384 | 13 | 377 | 20 | 406 |
| Esophagus Gastroesophageal Junction | 368 | 15 | 362 | 22 | 375 |
| Esophagus Mucosa | 465 | 13 | 453 | 26 | 555 |
| Esophagus Muscularis | 463 | 19 | 453 | 31 | 515 |
| Heart Atrial Appendage | 348 | 7 | 336 | 19 | 429 |
| Heart Left Ventricle | 295 | 6 | 292 | 9 | 432 |
| Kidney Cortex | 70 | 0 | 70 | 0 | 85 |
| Liver | 208 | 1 | 206 | 3 | 226 |
| Lung | 526 | 16 | 509 | 38 | 578 |
| Minor Salivary Gland | 207 | 2 | 203 | 6 | 162 |
| Muscle Skeletal | 521 | 15 | 504 | 34 | 803 |
| Nerve Tibial | 557 | 25 | 538 | 45 | 619 |
| Ovary | 233 | 4 | 230 | 8 | 180 |
| Pancreas | 274 | 6 | 270 | 10 | 328 |
| Pituitary | 349 | 12 | 343 | 18 | 283 |
| Prostate | 307 | 5 | 303 | 9 | 245 |
| Skin Not Sun Exposed Suprapubic | 523 | 20 | 507 | 39 | 604 |
| Skin Sun Exposed Lower leg | 572 | 18 | 555 | 39 | 701 |
| Small Intestine Terminal Ileum | 254 | 5 | 252 | 7 | 187 |
| Spleen | 319 | 11 | 311 | 20 | 241 |
| Stomach | 321 | 7 | 319 | 9 | 359 |
| Testis | 681 | 20 | 660 | 44 | 361 |
| Thyroid | 581 | 21 | 566 | 38 | 653 |
| Uterus | 185 | 2 | 184 | 3 | 142 |
| Vagina | 188 | 1 | 188 | 1 | 156 |
| Whole Blood | 377 | 8 | 363 | 24 | 755 |

